# Ecological and evolutionary insights into the diversification of Atlantic bluefin tuna

**DOI:** 10.1101/2025.09.15.675172

**Authors:** Chloe S. Mikles, Camille M.L.S. Pagniello, Eyal Bigal, Ben M. Moran, Jay R. Rooker, Aurelio Ortega, Fernando de la Gándara, Hugo Maxwell, Robert Schallert, Michael R. Castleton, Molly Schumer, Michael J. W. Stokesbury, Barbara A. Block

## Abstract

Identifying and preserving biological diversity is fundamental for the management of wild populations. The Atlantic bluefin tuna (*Thunnus thynnus*; ABT) is a vital species to the North Atlantic Ocean ecosystem that is now rebounding from decades of overfishing. This teleost fish is notable for its large body size, unique form of endothermy, and trans-oceanic migrations that enable individuals to move rapidly between spawning and foraging locations. Here we combine high-resolution whole genome sequencing data with spatial and environmental data from electronic tagging to improve our understanding of population structure in ABT. We analyzed whole genome sequences (n=82) from both larvae and adult fish representing the two recognized stocks (western and eastern) of ABT, which originate from geographically distinct spawning grounds. Analyzing these data, we identified 11,181,223 single nucleotide polymorphisms (SNPs), a dataset of unprecedented size and resolution. Coverage across the entire genome resulted in increased power in analyses of population structure, patterns of selection, and demographic history between the two recognized stocks. Notably, we report that both neutral and putatively adaptive SNPs are differentiated between populations, and through analyses of F*_ST_* outlier SNPs, we discovered candidate genes with potentially adaptive roles. We suggest that both demographic history and oceanographic variation of the spawning grounds have contributed to shaping bluefin tuna genomic diversity. Our results characterize adaptive variation that will be consequential for management decisions and critical for preserving locally adapted populations.

## I. INTRODUCTION

Determining biologically meaningful population structure is central to preserving biodiversity and aiding conservation efforts (Funk et al., 2012). In marine fisheries, managed populations are conceptually defined as a “stock,” and stock assessment methods rely on accurate information on life history strategies, migration patterns, and genetic diversity (Hilborn and Walters, 1992; Begg and Waldman 1999). However, understanding these dynamics in highly migratory fish species is challenging due to their extraordinary mobility at an ocean basin scale (Collette et al., 2011). For highly migratory species such as tunas and billfishes, which are characterized by large population sizes, high dispersal capabilities, and continuous distributions, population structure is challenging to detect, if present at all (Graves and McDowell, 2003; Pecoroaro et al., 2018; Mamoozdeh et al., 2019). Furthermore, there is potential for mismatch between definitions of biological populations and managed stocks, where a stock does not necessarily reflect the population genetic structure or spatial dynamics present in the system.

This mismatch can partly be due to unaccounted genetic diversity (either unknown or not incorporated into management strategies), or in some cases a result of changes in the environment that affect population dynamics (Waples et al. 2008; Reiss et al. 2009; Kerr et al. 2017). Thus, it is necessary to incorporate multidisciplinary (i.e., tagging and genetic techniques, see Jorgensen et al., 2010; Heuter et al., 2013) and adaptive management strategies (Maxwell et al., 2015) for studying and managing highly migratory fishes.

The Atlantic bluefin tuna *Thunnus thynnus* (ABT) is an iconic fish species renowned for its endothermic capabilities and specialized morphological and physiological adaptations that facilitate heat retention, thermogenesis, and long-distance migrations (Block et al., 1993, 2001, 2005; Bernal et al., 2001; Watanabe et al., 2015). ABT are a top predator with a key role in global fisheries (Macfadyen et al., 2021); therefore, its biology, migratory patterns, and population structure have been intensely studied (Galuardi and Lutcavage, 2012; Fromentin and Loupaszanski, 2014; Wilson et al., 2015; Arregui et al., 2018). Spatiotemporal variability in migratory behavior, stocks rebounding from decades of overfishing, and an incomplete understanding of population genetic structure, make the designation and delineation of appropriate management units both challenging and contentious (Fromentin and Ravier, 2005; Porch 2005).

ABT are managed by the International Commission for the Conservation of Atlantic Tunas (ICCAT) as two separate stocks, “western” and “eastern,” separated by the 45°W meridian and based on spatially separate and distinct spawning grounds in the Gulf of Mexico (“Gulf”), and the Mediterranean Sea (“Med”) (ICCAT 2018; Porch et al. 2019). These stocks are considered to be reproductively isolated, and although ABT spatially mix extensively on foraging grounds throughout their lifetime (Lutcavage et al. 1999; Block et al., 2005; Galuardi et al., 2010; Rooker at al., 2014, 2019; Wilson et al., 2015), electronic tagging and otolith chemistry studies indicate that adults demonstrate strong fidelity to these spawning areas over multiple years (Block et al., 2005; Rooker et al., 2008). To date, no tagged ABT has been tracked entering both the Gulf and Med spawning grounds. In addition, larval collections and tagging studies indicate temporal isolation in spawning behavior, where fish in the Gulf spawn earlier (April – June) (Teo et al., 2007; Wilson et al., 2015) than in the higher-latitude Med (May – August) (Cort and Loirzou, 1990; Schaefer, 2001).

The management models for ABT have advanced using the multistock age-structured tag-integrated model approach (Taylor et al., 2011), and current ICCAT management models increasingly account for spatial overlap of the two stocks by integrating electronic tag, mark-recapture, catch-per-unit effort, landings, and otolith microconstituent data to determine the spatial and temporal movement dynamics of ABT, as well as the contribution of each stock to the total catch (Carruthers et al. 2016). The eastern stock is estimated to be 10 times more abundant than the western stock (ICCAT, 2017), and although both eastern and western stocks have historically been severely depleted from overfishing over the past 60 years, both are considered to be recovering (ICCAT, 2021). ABT population structure has recently been proposed to be more complex than implied by the current two-stock management strategy (Fromentin et al. 2014; Brophy et al. 2020). Uncertainties remain regarding ongoing gene flow between the two stocks (Alvarado-Bremer et al., 2005; Johnstone et al., 2021; Diaz-Arce et al., 2024), as well as the potential for additional populations within the Mediterranean Sea (Carlsson et al., 2006), and more recently, in the Slope Sea where a newly described spawning area is proposed to exist along the North American slope waters (Richardson et al., 2016; Rodríguez-Ezpeleta et al., 2019; Aalto et al., 2023). The genetic origin of these fish is currently unknown, and fish that spawn here may be a distinct subunit of the population, or potentially western and/or eastern individuals utilizing an alternate spawning ground.

Genetic differentiation within ABT populations is inherently difficult to discern due to high effective population size, relatively recent divergence times of the bluefin lineages within the genus *Thunnus* (Ciezarek et al., 2018), and rapid radiations within Scombrid fishes over time (Block and Finnerty, 1994). Signals of population structure from genetic data in ABT are most pronounced between Gulf and Med populations (Boustany et al., 2008; Puncher et al. 2018, Rodriguez-Ezpeleta et al. 2019), and have been determined using multiple genetic approaches, including microsatellites, mitochondrial DNA and single nucleotide polymorphisms (SNPs).

Within the Med, several microsatellite studies describe population structure between sampling sites (Carlsson et al. 2004; Vella et al. 2009; Riccioni et al. 2010, 2013); however, studies using SNP markers have not to date detected significant genetic structure (Antoniou et al., 2017; Rodríguez-Ezpeleta et al., 2019). Subsets of SNPs serve as valuable markers for population studies due to their broad and abundant distribution across the entire genome and have recently been used to assign stock-of-origin (Puncher et al., 2018; 2022; Rodríguez-Ezpeleta et al., 2019).

The use of whole genome sequencing to survey genomic diversity provides a significant advantage by allowing for SNP discovery across the entire genome, in contrast to microsatellite or reduced-representation sequencing techniques. In addition, the availability of an annotated reference genome assembly for ABT (NCBI accession # GCF_963924715.1; Block et al, *in submission*) has enabled the investigation of genome-wide variation in ABT. While the discriminatory power of whole genome sequencing is not necessary for stock assignment, it is a powerful tool to identify adaptive genetic diversity, perform genome scans, and infer divergence times between populations (Clucas et al., 2019). Together, this makes it particularly useful as a tool for improved management of non-model organisms (Funk et al., 2012; Wilson et al., 2015).

In this study, we combined high-resolution genomic data from whole genome sequencing with spatial and environmental data from electronic tags to improve our understanding of population structure in ABT. We analyzed genomic data from both larvae collected at the specific spawning location with net tows, and adults that traveled to the region (fin clips from tagged fish) to provide representative samples of the Gulf and Med populations (*n* = 82). We characterized the population genomic structure of these individuals, investigated the role of adaptive and neutral processes in shaping differentiation between populations, and inferred demographic history. We also explored the potential for local adaptation to distinct oceanographic conditions of the spawning grounds, to aid in evaluating the adaptive capacity of bluefin tuna.

## II. RESULTS

### Whole genome resequencing results

We generated whole genome resequencing data on a final dataset of 82 Atlantic bluefin tuna representing these two primary spawning grounds, the Gulf of Mexico (Gulf, n=43) and Mediterranean Sea (Med, n=39), split between larvae (Gulf n =26, Med n =13) and adults described above (Gulf n=16, Med n =27). All but four samples were sequenced at a target coverage of 7x, and the remaining at a target coverage of 47x.

The mean percentage of reads mapped to the ABT reference genome was 99.44%, and the mean coverage was 10.60x. (Table S1). After quality filtering and variant calling, we obtained 11,181,223 SNPs for subsequent analyses. The mean missing data was 3.9%. To generate a dataset of likely neutral unlinked SNPs, we pruned for linkage and additionally excluded SNPs within 10kb of exons, resulting in a greatly reduced dataset of 1,208,268 SNPs.

### Summary of adult migratory behavior

To infer subpopulation structure in adult ABT, we generated whole genome sequences from the fin clips of electronically tagged fish (n= 43; see STAR methods and Table S1 for tag deployment metadata). These fish underwent distinct migrations to spawning grounds in the Gulf (n=16) or Med (n=27). No tagged ABT visited both spawning regions, thus we classify individuals as either “Gulf” or “Med” based on entry into their respective spawning ground (Wilson et al., 2015). In addition, six ABT were tagged within the Med and were thus classified as “Med” origin. Satellite tags on ABT recorded data for an average of 271 ± 110.4 days (range 4-466 days). Figure 1 shows the spatial distribution of the fish, and more detailed individual tracks can be found in Supplemental Figure 1.

**Figure 1.**
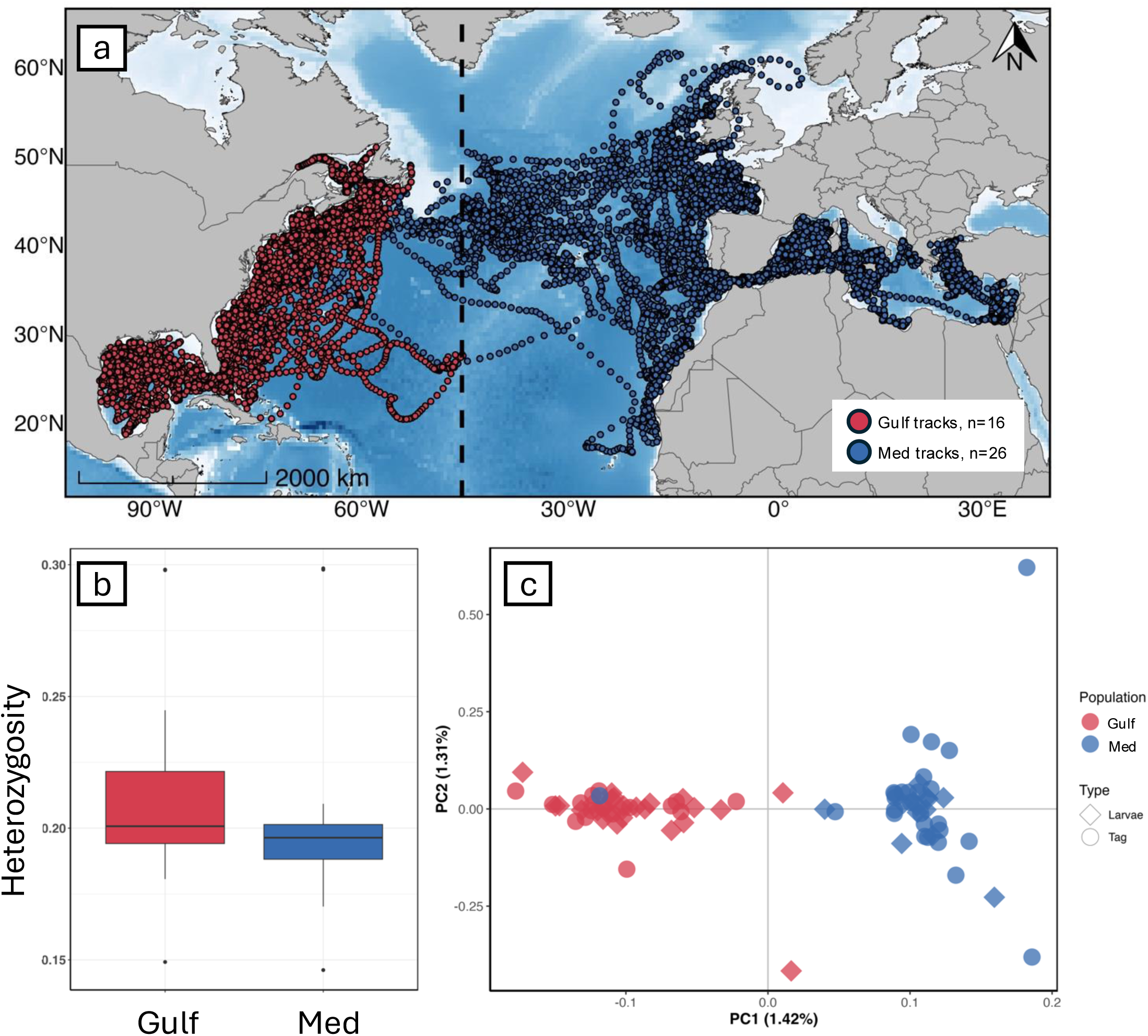
(A) State space modelled tracks for adult Atlantic bluefin tuna tagged with pop-up satellite archival tags demonstrate spatial separation of migration patterns by population (Gulf, red; Med, blue). (B) Individual heterozygosity by population. (C.) Principal component analysis using 11,181,223 SNPs reveals genomic separation of populations.

Gulf ABT were all tagged as adults (mean length 272.4 ± 12.1 cm curved fork length (CFL)) in Nova Scotia, Canada. Geolocation data indicates that following tagging in September, fish departed from the Gulf of St. Lawrence in the late fall or early winter, migrated southward along the U.S. east coast, and entered spawning grounds in the Gulf between November and May (mean entry day: February 17^th^ ± 50 days). ABT resided in the Gulf for an average of 95 ± 43 days, typically departing in spring to early summer (mean exit date May 17^th^ ± 45 days), and returned north to coastal and offshore waters off New England and Canada. Eight of the tagged ABT displayed fidelity to their relative tagging locations in Nova Scotia and six tags were recovered, providing high resolution archival datasets. All fish remained west of the 45°W meridian in the northwestern Atlantic during the course of their migrations.

Med adults examined for this study were electronically tagged in either Canada (n=8), Ireland (n=12), Spain (n=1), or the Med (n=6). All fish tagged outside of the spawning ground crossed through the Straights of Gibraltar and entered the Med with a mean entry date of May 13^th^ ± 38 days. At tagging, these ABT had a mean length of 234 ± 26.1 cm CFL. The five adults tagged inside of the Med had a mean length of 139.8 ± 8.5 cm CFL. These fish resided in the Med for the entirety of the tagging period (Med “residents”). Non-resident Med fish spent on average 67.5 ± 43 days in the Med, and typically exited in late summer (August 14^th^ ± 24 days). Med adults undertook diverse migratory pathways that varied both between and within their original tagging locations.

### Population structure and genomic diversity

Whole genome sequences allowed us to characterize the contributions of both adaptive and neutral markers in shaping differences between populations. Principal component analysis (PCA) revealed subtle, but clear genome-wide differentiation between Gulf and Med ABT (Figure 1c). Using the full SNP dataset, the first principal component axis (PC1) explained 1.42% of the variation and separated the majority of Gulf and Med ABT, with no differences in clustering between the two types of sample groups (adults and larvae). The putatively neutral dataset showed a similar pattern (Figure S2). In a PCA conducted with the tagged fish separately, PC1 explains 2.56% of the differentiation, and PC2 explains 2.46%, suggesting increased differentiation within the Med samples and not the GULF (Figure S3). Admixture was run using the thinned SNP dataset identified K=1 as the best supported cluster. We did not detect sub-structuring in higher K-values (Figure S4).

F*_ST_* estimates based on the full dataset suggest very low levels of divergence between Gulf and Med (mean F*_ST_* = 0.0026 ± 0.009; weighted mean F*_ST_* = 0.0032 ± 0.048). Genome-wide estimates of *d_xy_* were also very low (mean 0.006 ± 0.003). Individual SNP F*_ST_* values ranged from 0 to 0.77. There were very slight differences in heterozygosity between Gulf (mean ± SD = 0.21 ± 0.03) and Med (0.20 ± 0.03) (Figure 1b). Genome-wide patterns of nucleotide diversity and Tajima’s D were similar between Gulf and Med (Figure S5 and S6).

### Demographic history

To investigate long-term demographic history of ABT, we estimated effective population size (N_e_) using PAIRWISE SEQUENTIALLY MARKOVIAN COALESCENT (PSMC). We ran PSMC on the high coverage genomes from one GULF adult (GULF-24) and one Med adult (MED-7). Because there are a range of estimates for generation length of the two populations (13-16 years for Med and 13-19 years for Gulf; Colette et al., 2021), we scaled the population using a generation time of 13 for both populations and a mutation rate of 3×10^-9^ per base per year (Figure 2). We additionally scaled the population using generation lengths from 8-16 (Figure S15, S16). The N_e_ of both populations steadily declined during the mid Pleistocene, plateauing at an estimate of 350,000 before splitting to an expansion of the Med a bottleneck of the Gulf. Our results show that trend reverse, where the Gulf expands and the Med declines. Both populations maintained a steady population size through the Last Glacial Maximum, with the Gulf N_e_ remaining higher than the Med.

**Figure 2.**
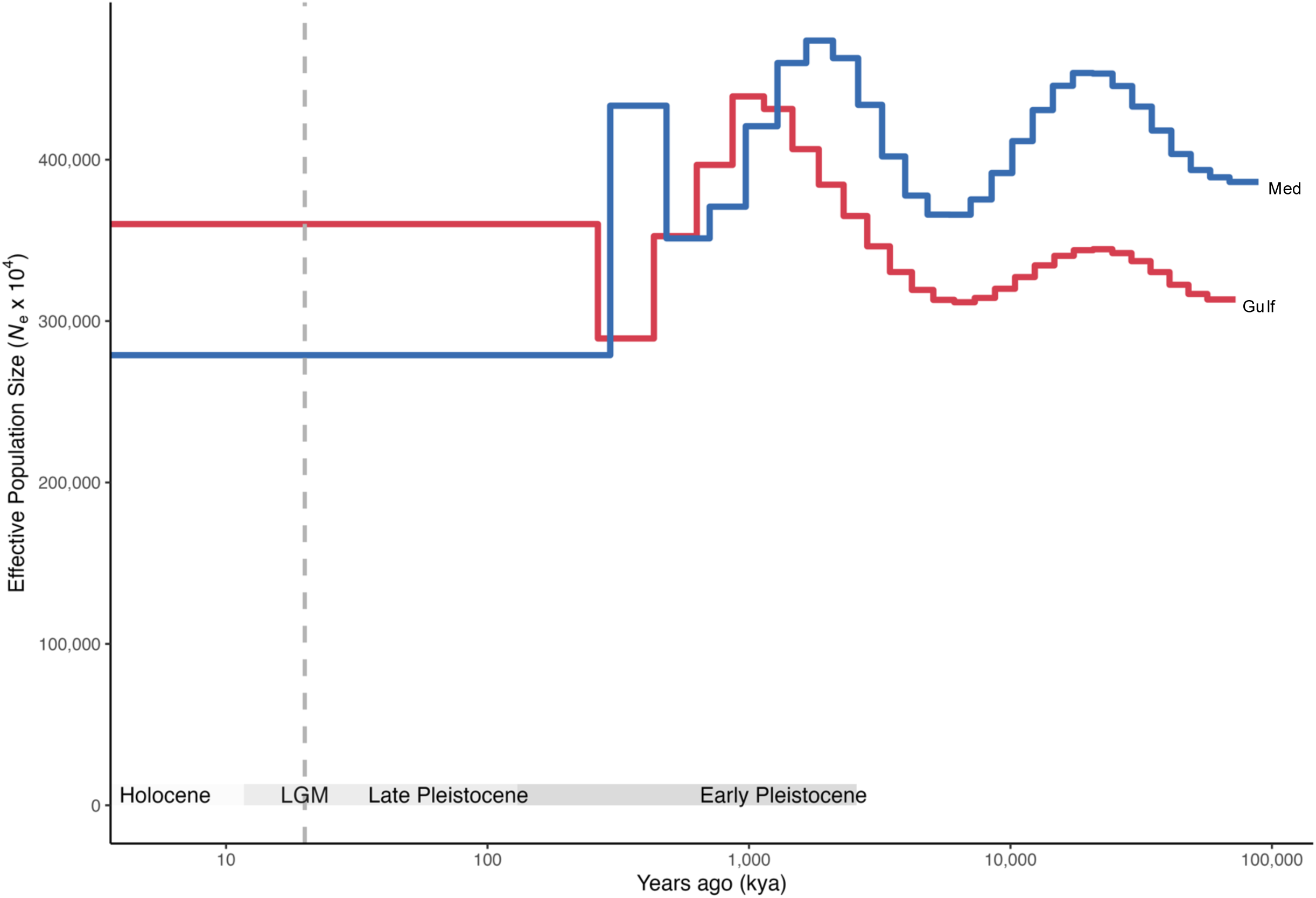
Demographic history of Atlantic bluefin tuna. Effective population sizes were reconstructed using pairwise sequentially Markovian coalescent (PSMC) assuming a generation time, g, of 13 years and a mutation rate, µ of 3×10-9.

### Patterns of selection

Manhattan plots of pairwise F*_ST_* in 25 kb windows revealed very low levels of genome-wide divergence with several distinct elevated peaks occurring on chromosomes 1-12, 14-24. (Figure 3a). Genome-wide patterns of *d_xy_* were overall very low, though peaks in F*_ST_* corresponded with elevated values for *d_xy_* (Figure S7). PCA of the outlier regions revealed very strong separation of the Gulf and Med populations, with no individual overlapping into the opposite population’s cluster (Figure S8). The first PC axis explained 18.5% of the variation.

**Figure 3.**
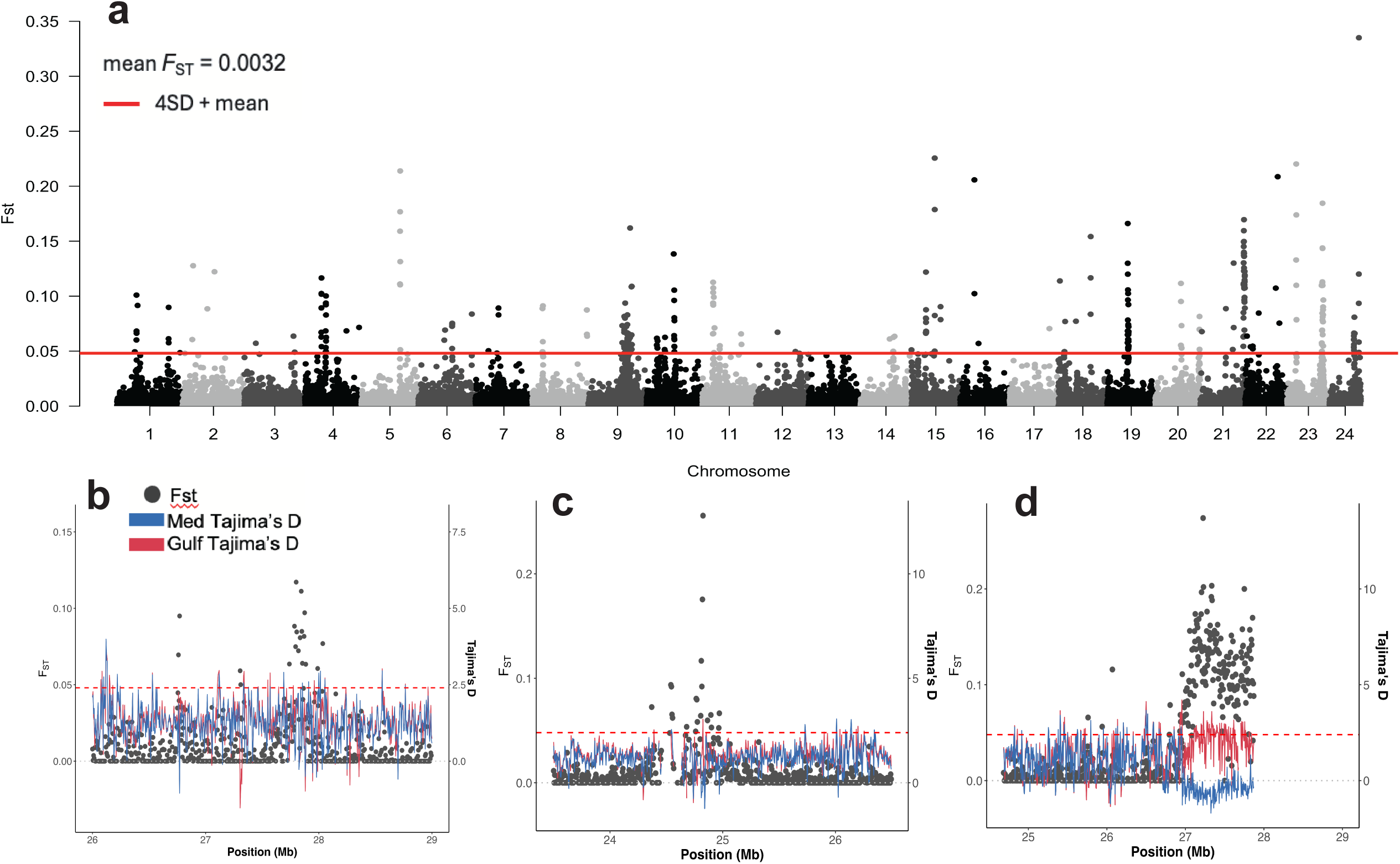
(A) Genome-wide estimates of pairwise Fst between Gulf and Med populations of Atlantic bluefin tuna. Pairwise Fst (grey dots) and Tajima’s D by population (Gulf, red; Med, blue) plotted in 5kb windows and centered on significant peaks for Chromosome 1 (b), Chromosome 8 (C), and Chromosome 21 (d).

We observed 298 outlier SNPs with F*_ST_* values greater than four standard deviations above the mean. The outlier SNPs had a mean F*_ST_* of 0.20 ± 0.02, and we identified 219 unique genes that overlapped within ±20kb windows of these SNPs (Table S2). The candidate genes associated with SNPs exhibiting elevated divergence between GULF and Med populations are linked to several biological functions with potential adaptive relevance. These include 10 elevated windows, three of which occur on Chromosome 21, associated with voltage-gated calcium ion membrane transport proteins (*trpc6a, guca3e, MLNR*), and genes with biological functions that include visual perception and organ development (*rcbtb1*). Additionally, eight windows contain genes related to the solute carrier family (SLC), four are related to cardiac function (*mylk4a, smad4a, foxo1a, LUZP1*), and two are related to reproduction (*spata16, fshr*).

Other notable gene functions include development of retinal vasculature and camera-type eye (*med27*), fin development (*fndc3a, fgfr1a*), striated muscle contraction (*MYH13, tnnt3*), and circadian behavior (*nmur3*). A single gene, *dnmt3aa,* located on Chromosome 8 functions in DNA methylation, response to hypoxia and temperature stimulus, epigenetic memory and learning, and swimming behavior.

On Chromosome 21, the peak in F*_ST_* corresponds with a population-level split in Tajima’s D, where negative values of Tajima’s D were observed for the Med population, and positive for the Gulf population (Figure 3d). Preceding the peak, Tajima’s D values were positive and patterned closely for both populations. Values for *d_xy_* were overall low, reaching near-zero values under the peak in F*_ST_* (Figure S9). This F*_ST_* peak was associated with genes related to respiration, vascularization, and reproduction, among others (Table S2).

### Oceanographic conditions of distinct spawning grounds as a predictor of divergent genotypes

We conducted genome-wide association analyses using GEMMA to explore correlations between genetic signatures and environmental and oceanographic variation between the two spawning grounds. We utilized measurements from tag-derived positions when adult ABT occupied either the Gulf or Med, including daily mean values of surface temperature (°C), sea surface salinity (PSU), mixed layer depth (m), sea surface height (m), and eddy kinetic energy (m^2^/s^2^). We also included the calendar day of entry to the respective spawning ground. ABT experienced significantly lower sea surface temperature, higher sea surface salinity, deeper mixed layer depth, and shallower sea surface height in the Med compared to the Gulf (Table 1, Kruskal-Wallis p-values < 0.001). Eddy kinetic energy was the same for both regions (p-value = 0.8). Mean daily values are plotted by individual in Figures 4 and S11-14. Two Med ABT resided in the Med for ∼1 year (MED-10 and MED-41; 351 and 367 days, respectively); the environmental variables experienced by these fish were not necessarily representative of spawning conditions and differed substantially from the migratory Med fish, thus these individuals were given NA values for GEMMA.

**Figure 4.**
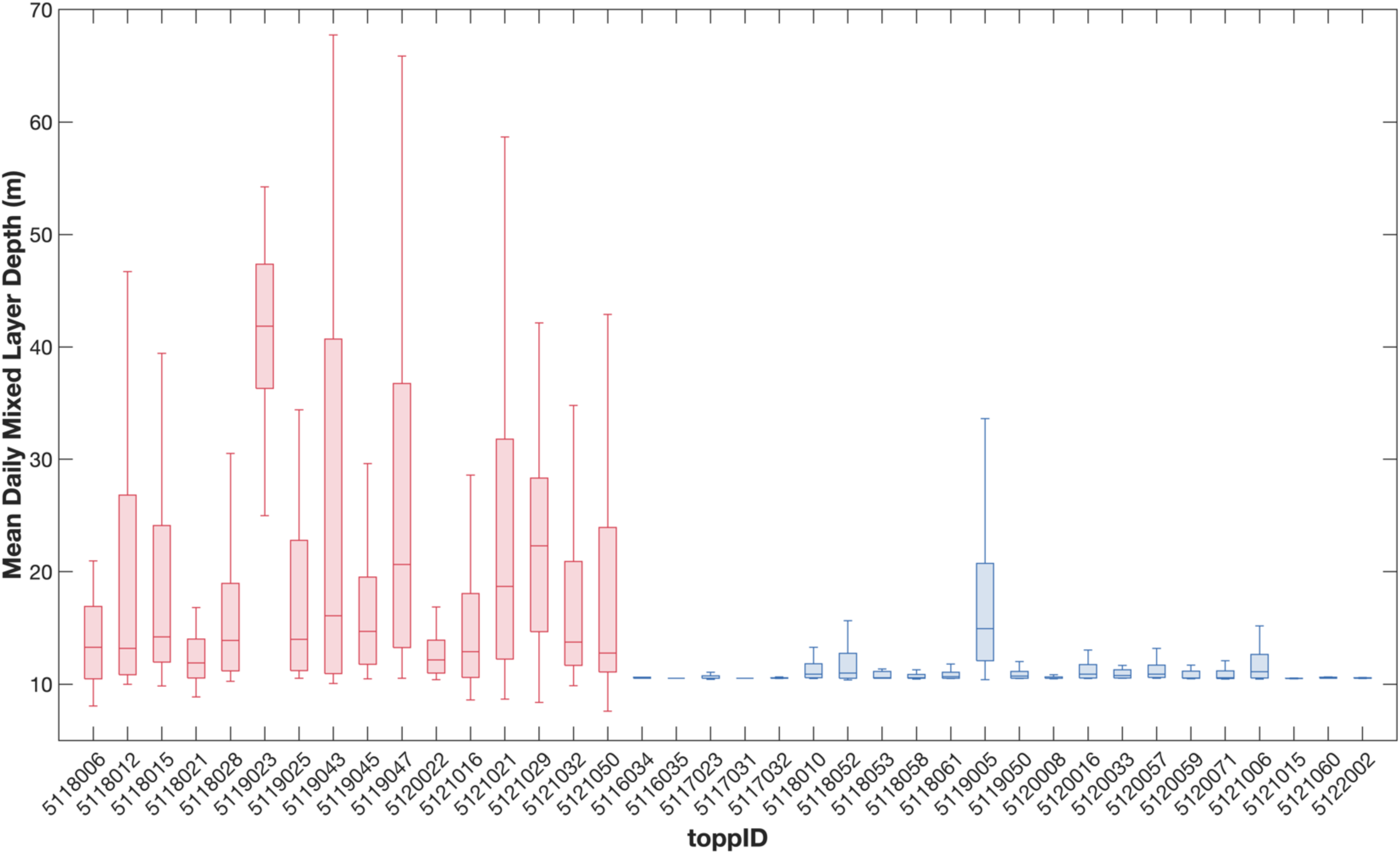
Mean mixed layer depth (MLD) in meters for adult bluefin tuna while occupying spawning grounds in Gulf of Mexico and Mediterranean Sea. Gulf adults are coloured red, Med adults blue.

**Table 1.** Oceanographic conditions experienced by adult ABT while occupying distinct spawning grounds in the Gulf of Mexico (Gulf) or Mediterranean Sea (Med). Values represent daily means of sea surface temperature (SST), sea surface salinity (SSS), mixed layer depth (MLD), sea surface height (SSH), and eddy kinetic energy (EKE).

|  | GOM | Med | P-value<br>(difference<br>between<br>means) |
| --- | --- | --- | --- |
| SST (°C) | 25.2 +/- 1.1 | 22.3 +/- 1.5 | **p < 0.001** |
| SSS (PSU) | 35.9 +/- 0.4 | 37.6 +/- 0.5 | **p < 0.001** |
| MLD (m) | 20.6 +/- 7.0 | 11.4 +/- 1.5 | **p < 0.001** |
| SSH (m) | -0.05 +/- 0.04 | -0.4 +/- 0.06 | **p < 0.001** |
| EKE (m<sup>2</sup>/s<sup>2</sup>) | 0.04 +/- 0.03 | 0.04 +/- 0.03 | 0.8 |

**Table 2.**
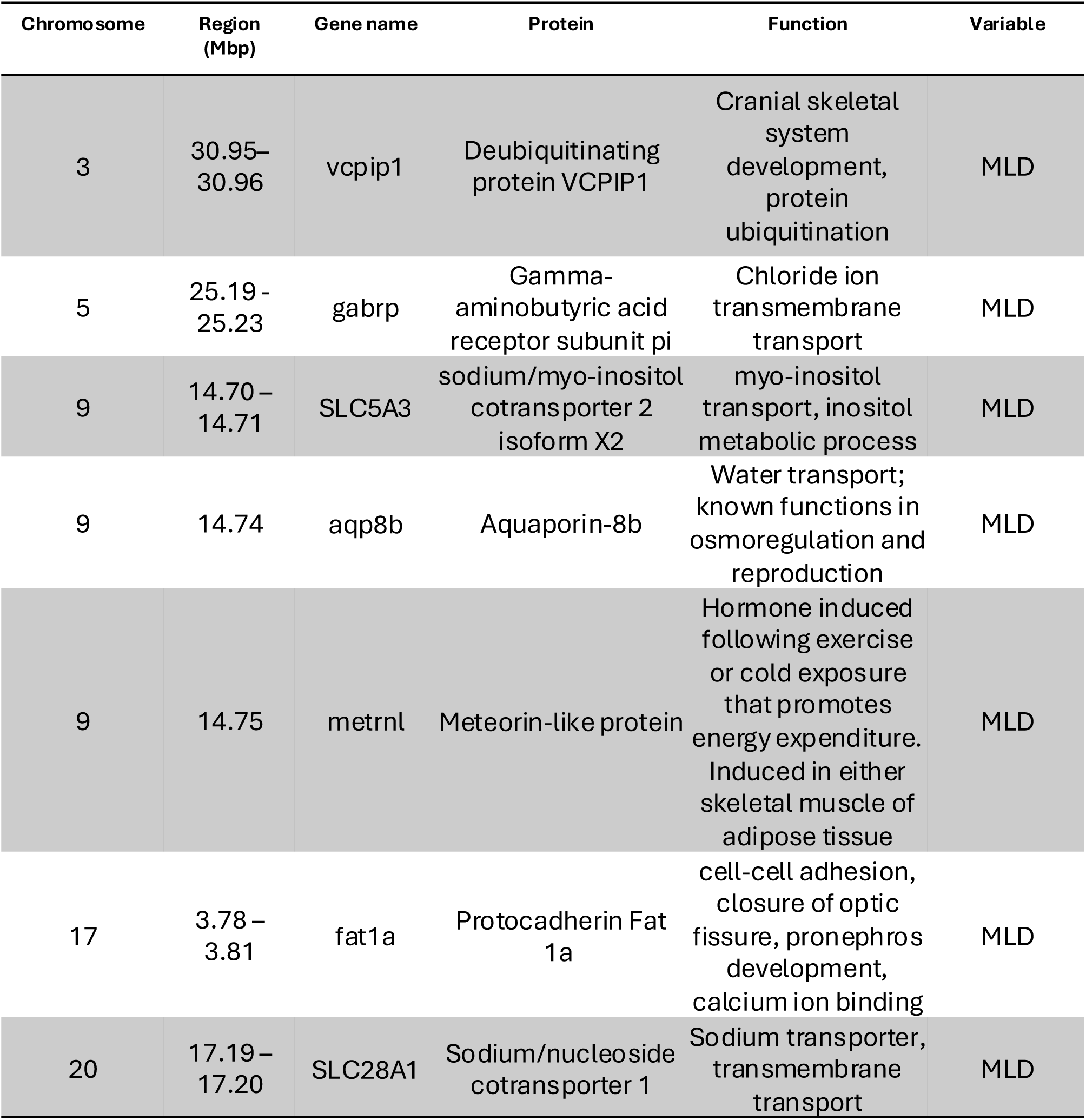
Candidate genes that are significantly associated with mean MLD. Subset of genes with putative physiological relevance; all genes are detailed in Table S3.

| Chromosome | Region (Mbp) | Gene name | Protein | Function | Variable |
| --- | --- | --- | --- | --- | --- |
| 3 | 30.95–30.96 | vcpip1 | Deubiquitinating protein VCPIP1 | Cranial skeletal system development, protein ubiquitination | MLD |
| 5 | 25.19 - 25.23 | gabrp | Gamma-aminobutyric acid receptor subunit pi | Chloride ion transmembrane transport | MLD |
| 9 | 14.70 – 14.71 | SLC5A3 | sodium/myo-inositol cotransporter 2 isoform X2 | myo-inositol transport, inositol metabolic process | MLD |
| 9 | 14.74 | aqp8b | Aquaporin-8b | Water transport; known functions in osmoregulation and reproduction | MLD |
| 9 | 14.75 | metrnl | Meteorin-like protein | Hormone induced following exercise or cold exposure that promotes energy expenditure. Induced in either skeletal muscle of adipose tissue | MLD |
| 17 | 3.78 – 3.81 | fat1a | Protocadherin Fat 1a | cell-cell adhesion, closure of optic fissure, pronephros development, calcium ion binding | MLD |
| 20 | 17.19 – 17.20 | SLC28A1 | Sodium/nucleoside cotransporter 1 | Sodium transporter, transmembrane transport | MLD |

Association tests in GEMMA identified 35 SNPs that were significantly correlated with the environmental variables of interest based on a cutoff of PIP >0.01. This included 13 SNPs associated with mean mixed layer depth, 11 with mean sea surface height, 3 with sea surface salinity, 3 with sea surface temperature, and 5 with spawning ground entry day (Table S3). The mean PIP value for these outlier SNPs was 0.22 ± 0.32. We identified 42 unique outlier genes that overlapped within ±20kb windows of these SNPs (Table S4). Candidate genes identified through GEMMA included several with potential functional relevance: thermoregulatory gene *metrnl,* osmoregulation *aqp8b,* and ion transport (*fat1a, SLC28A1*) associated with mixed layer depth, swimming and neuroregulatory gene *nrcama* and ion transport gene *hcn2b* associated with SSH, and ion transport gene *kcn13b* associated with sea surface temperature.

## V. DISCUSSION

Our novel integration of whole genome data with marine spatial data from electronic tags provides a robust analysis of ABT population structure. Utilizing over 11 million SNPs, we demonstrate the utility of incorporating whole genome data to delineate and characterize populations, and we posit that oceanographic variation of the spawning grounds has contributed to differentiation of the two primary populations of Atlantic bluefin tuna. We additionally discuss candidate genes that are associated with regions putatively under selection.

### Spatial and genomic population delineation

Integrating spatial data from electronic tagging with whole genome data provides valuable insights into the population structure of ABT. Our use of genome-wide data to describe population structure of ABT unequivocally distinguishes Gulf and Med populations (Figure 1c), revealing a more defined pattern compared to previous genetic studies utilizing reduced-representation SNP datasets and microsatellites (Boustany et al., 2008; Rodriguez-Ezpeleta et al., 2019; Díaz-Arce et al., 2024). The population structure detected from whole-genome data is concordant with the spatial structuring of adult movement patterns revealed from electronic tags, which are herein restricted west of 45°W for Gulf, and primarily concentrated east of 45°W for the Med (Figure 1a). Prior tagging and otolith chemistry studies have established that the two populations overlap on foraging grounds in the Northwest Atlantic (Galuardi et al., 2012; Rooker et al., 2014, 2019; Kerr et al., 2020), and as adults are reproductively isolated in the spawning season through repeat use of their natal spawning grounds (Block et al., 2005). Consistent with other SNP datasets, we did not detect any population sub-structuring within the Med (Puncher et al., 2018; Rodriguez-Ezpeleta et al., 2019), however our sampling design did not have enough representatives from the three different regions within the Med to fully test this hypothesis. To date there are conflicting studies of Med population structure, with several studies reporting genetic differentiation within the Med (Carlsson et al., 2004, 2007; Vella et al., 2006, 2009; Riccioni et al., 2010, 2013), while others detect none (Alvarado-Bremer 1999, 2003; Boustany et al., 2008; Viñas et al., 2011; Antoniou et al., 2017; Puncher et al., 2018; Rodriguez-Ezpeleta et al., 2019). Future work should prioritize whole genome approaches that increase the larval sample size to include adequate representatives from the three Med regions, western, central, and eastern, given that adults travel through regions and do not necessarily spawn there.

Given the complexities of ABT population structure (Fromentin et al. 2014; Brophy et al. 2020), it is beneficial to include electronic tagging data with genomic analyses to contextualize results. Additionally, distinct life history strategies and migratory patterns may have a genomic basis, as has been observed in Pacific salmon (*Oncorhynchus* spp., Hale et al., 2013; Prince et al., 2017) and Atlantic cod (Nordeide et al., 2011; Clucas et al., 2019). In ABT, electronic tagging data has demonstrated that the Med hosts both migratory and resident populations (Fromentin and Lopuszanski 2013; Cermeño et al., 2015), and two ABT included in our study demonstrated residential behavior to the Med. It will be advantageous in the future to increase electronic tag deployments (as well as the duration of the tag deployment) to investigate migratory vs. residential behavior from a genomic perspective.

Interestingly, one spatially-assigned Med fish clustered with the Gulf population in the PCA (Figure 1c). This fish was tagged in the Gulf of St. Lawrence, Canada, and traveled to presumably spawn in the far eastern Med. This anomaly is notable for several reasons. First, the Gulf of St. Lawrence was historically comprised of ∼100% GULF fish, determined using otolith δ^18^O and δ^13^C (Rooker et al. 2008, Schloesser et al. 2010). SNP data revealed an increasing proportion of Med-origin fish in this region in recent years, as well as higher proportions of younger fish (Puncher et al., 2022). Second, tagged Med fish primarily travel to the western and central Med to spawn (Block et al., 2005; Cermeño et al., 2015), and it is extremely rare that a western-tagged fish reaches the eastern Med (Walli et al., 2009). Even among large adults tagged in the eastern Atlantic, travelling to the eastern Med is rare (Pagniello et al., 2023; Ferter et al., 2024), and movement patterns of fish that reside in the eastern Med are poorly understood (Arrizabalaga et al., 2019). Increased attention to studying the movement patterns and genomic diversity of ABT from the eastern Med would be beneficial to fill important data gaps and improve our understanding of ABT biodiversity.

### Evolutionary forces driving diversification in ABT

Our results provide new insights into the drivers of divergence in ABT populations. Our PSMC analyses demonstrate that the Gulf and Med populations have undergone a series of population expansions and contractions that likely shaped present-day population structure. The exact timing of these changes in N_e_ are uncertain without knowledge of the true mutation rate for ABT; however, the Pleistocene era can be characterized by transitions from glacial to interglacial periods that drastically altered global sea levels and thus fragmented marine habitats (Ayalon et al., 2002). Glacial periods reduced marine habitat and shifted ocean currents, which has been shown to cause population bottlenecks and result in population genetic structure in many marine species (Ludt and Rocha, 2015). As far as inferring recent demographic history, PSMC lacks the resolution to estimate N_e_ since bluefin tuna have been exploited by humans, though Andrews et al. (2021) found no evidence of recent bottlenecks comparing ancient and contemporary samples of ABT.

We highlight the utility of investigating both neutral and potentially adaptive variation in ABT. While neutral variation has historically been used to delineate fisheries stocks (f et al., 1997; Shaklee et al. 1999; Beacham et al. 2004), it is known that adaptive variation is a crucial part of species’ diversity and can be highly relevant to management issues (Crandall et al. 2000; Funk et al. 2012; Pecoraro et al. 2018). For example, an increasing number of studies on Atlantic cod using next generation sequencing techniques have identified adaptive divergence among populations with high levels of gene flow, despite weak or undetected differentiation at neutral loci (Nielsen et al. 2009; Therkildsen et al. 2013; Clucas et al. 2019). In our study, the full and neutral SNP datasets describe similar population structure, though the full SNP dataset explains slightly more differentiation. Despite very low levels of F*_ST_* and dxy genome-wide, there were several F*_ST_* outlier peaks with moderate levels of differentiation. Future studies should investigate potential structural variation of the bluefin tuna genome (i.e., presence of chromosomal inversions) and how that may affect recombination rates (Akopyan et al., 2024). Our results suggest multiple evolutionary forces have played a role in shaping the population genetic variation of ABT, which could be disentangled in future genomic studies.

We identified 219 unique candidate genes (Table S2) associated with these putatively adaptive SNPs, with functions related to sodium and calcium ion transport, cardiac function, visual perception, reproduction, swimming, and circadian behavior. One interesting gene family are the solute carriers (SLC genes), which are essential in transporting compounds across membranes and maintaining electrochemical gradients. For example, *slc16a1* facilitates the transport of lactate, which is essential for anaerobic metabolism and has elevated expression in zebrafish *Danio rerio* stressed with hypoxic conditions (Ton et al., 2003). This mechanism is likely very important in bluefin tuna, which produce extremely large quantities of lactic acid during burst swimming (Brill, 1996). Another gene of interest is *SLC25A14,* is a mitochondrial proton carrier of the SLC25 family (also known as uncoupling proteins). Uncoupling proteins can play an important role in dissipating the proton gradient for thermogenesis (Kunji et al., 2021). While present in zebrafish and carp *Cyprinus carpio,* these genes can have alternative functions not related to thermogenesis, as they are ectothermic fishes (Stuart et al., 1999).

However, the uncoupling proteins may have a role in thermogenesis of the endothermic bluefin tuna. These proteins have been demonstrated to be critical for regulating the mechanism of mitochondrial proton leaks (Stuart et al., 2001), a key source of thermogenesis in red muscle tissue (Block 1994, Duong et al. 2006).

We discovered several genes that could be useful in broader studies regarding ABT biology. There were three genes associated with reproduction, *pgr* (ovulation), *dcaf13* (oocyte growth), and *spata16*, which could be used to aid the identification of sex-determining regions in ABT. Additionally, the gene *dnmt3aa*, which has interesting functions related to swimming behavior in zebrafish, is better known for its control of DNA methylation patterns (Lai et al., 2020). This gene specifically can be used for epigenetic aging by analyzing patterns of DNA methylation across the genome and can be used as a non-invasive tool to age fishes (Piferrer and Anastasiadi, 2023). These genes and many others require further study to examine amongst *Thunnus* species, particularly for understanding resiliency to external stressors such as fishing pressure and climate change.

### Environmental drivers

Environment-associated traits contribute to species’ future adaptive capacity, though can be difficult to assess in migratory species that face different selective pressures during both their annual migratory cycle and different life stages (Thurman et al., 2020; Bay et al., 2021). For fish such as ABT, an annual migration cycle may take a single individual across a variety of oceanographic conditions, from the cold, fresher waters of the Gulf of St. Lawrence to the warmer and more saline waters of the Med. To investigate the role of the environment in shaping genomic diversity, we leveraged our unique dataset that includes individual-level genomic, behavioral, and environmental data. As bluefin tuna spawning behavior is closely linked to oceanographic conditions that provide optimal success for eggs and larvae (Teo et al., 2007; Muhling et al., 2010, 2013; Reglero et al., 2018), we hypothesized that there may be evolutionarily significant variation as a result of environmental differences of these two spawning grounds. Analyses of tagging data from adults indicate there exist slight differences in preferred temperature for spawning between the Gulf and Med, where ABT in the Gulf prefer 24 – 27°C (Block et al., 2005; Teo et al., 2007), and in the higher-latitude Med, prefer slightly cooler waters between 23.5 – 25°C (Alemany et al., 2010). Metrics such as primary production, sea surface currents, and SSH are also important for predicting suitable spawning habitat for ABT (Druon et al., 2016).

Using the electronic tagging data on adult ABT, we extracted oceanographic variables experienced by the fish while occupying their spawning grounds. Rather than comparing general metrics of the Gulf and Med, we utilized daily data points from the tracks during the exact period the tagged animals occupied each spawning region, aiming to hone in on the specific environmental conditions for which animals potentially are selecting. We found that ABT in the Gulf and Med experienced significant differences in mean SST, SSS, SSH, and MLD, all important metrics for spawning bluefin tuna and the survival of their larvae (Teo et al., 2007; Alemany et al., 2010; Muhling et al., 2013; Russo et al., 2022). Through GEMMA, we detected SNPs associated with these environmental metrics, including 13 SNPs associated with mean mixed layer depth, 11 with mean sea surface height, 3 with sea surface salinity, and 3 with sea surface temperature (Table S3). We additionally detected 5 SNPs related to migration timing (calendar day of entry to the spawning ground). Interestingly, we detected several genes with functional relevance, particularly in ion transport. For example, *Aqp8b,* aquaporin, which functions in osmoregulation, was significantly associated with mixed layer depth. *Apq8b,* along with other aquaporin orthologs, are important for water homeostasis and kidney function in teleost fishes. The closely related Aquaporin 1 is an important gene for oocyte hydration, which is particularly important for the buoyancy and thus survival of pelagic fish larvae (Finn and Fyhn, 2000). In ABT, aquaporin has been demonstrated to be differentially expressed in male versus female gonads (Gardner et al., 2012) and may be an important gene for understanding evolutionarily significant variation.

To our knowledge, this is the first analysis of genotype by environment interactions in ABT. We utilized GEMMA to discover SNPs associated with the environmental differences experienced (or selected for) by ABT, which may indicate local adaptation. Our results make a compelling case for possible selection for specific oceanographic conditions by ABT. If this is the case, characterizing the influence environment has on divergent ABT genotypes is particularly important as conditions of the spawning grounds may shift in the future with climate change (Muhling et al., 2011). While these results are interesting, there are important limitations. First, our sample size of spawning adults is low and does not include all spawning locations of ABT. Future sampling should include additional spawning locations such as the Slope Sea (Richardson et al., 2016), as well as include broader and more finite sampling within the Gulf and Med to make more robust conclusions about genotype environment associations (Alvarado et al., 2022). The satellite tag data also is not transmitted at a resolution high enough to analyze exactly where and when fish are spawning. Thus, we cannot with certainty differentiate the specific environmental variables where fish are spawning versus traveling within the Gulf and Med basins. Second, we cannot truly determine which mutations are physiologically relevant in ABT without functional studies; however, future studies could combine RNA-sequencing data with programs such as AlphaFold (Jumper et al., 2021) to enable investigation of gene function in ABT.

### Management implications

Identifying and delineating populations often benefits from the utilization of multidisciplinary approaches (see Adams et al. 2003; Boustany et al. 2008; Pearce et al. 2008; Jorgensen et al. 2010; Hueter et al. 2013). Our study integrates spatial data from electronic tagging with genomic data, and together, these techniques can increase the confidence behind inference of population structure. Our identification of highly-differentiated SNP markers that separate Gulf and Med populations will be a powerful tool for genotyping fish of unknown origin, particularly as stock assessment models adapt to include genomic data. Previously, a 96-SNP panel was used to assign ABT of known origin to their stock, with a success rate of 81% and 83% for Gulf and Med, respectively (Rodríguez-Ezpeleta et al., 2019). Similar methods have been used to genotype fish of unknown origin on mixed foraging grounds in the Northwest Atlantic Ocean (Puncher et al., 2022). Future work utilizing SNP markers could improve assignment probabilities of catch data, assign stock to spatial data for juveniles, and assign sex to investigate stock- and sex-specific movement patterns (Nielsen et al., 2024). However, any further work on stock assignment to either “Gulf” or “Med” is complicated also by uncertainties surrounding fish spawning in the Slope Sea region (Richardson et al., 2016; Rodríguez-Ezpeleta et al., 2019; Aalto et al., 2023). Genetic analysis of samples from the region suggests potential for an admixed population (Díaz-Arce et al., 2024), therefore stock structure of ABT (and future management) may be more complex than Gulf or Med assignment. Additional work utilizing various techniques and additional sampling sites is necessary to improve understanding of ABT population structure and dynamics.

## Acknowledgments

We extend our thanks to members of the Block Lab Drs. Shaili Johri and Emil Aalto for advice and discussions throughout the project. We thank Ted Reimer for assistance in preparing tags, tagging, fin clip sample collection, and data management at sea. Erik Hanson, Brendan Cornwell, and Victoria Grant for helpful advice on bioinformatic data processing and analyses. We are grateful to Dr. Francisco Alemany of ICCAT for support of the genetic work and acquisition of samples, as well as for providing some of the satellite tags. Tagging of Atlantic bluefin tuna has been going for many years and we thank all of the captains, crew, and anglers who have enabled the deployment of electronic tags. All tagging was conducted under Stanford University Institutional Animal Care and Use oversight to BAB (#10786), and with national permits as follows: Animal Welfare License (Project AE19121/P003) as required under Directive 2010/63 /EU and Irish Government S.I. No. 543 of 2012; Fisheries and Ocean Scientific License to Fish #SG-RHQ-19–120 and Acadia Animal Care Protocol # 12–16R#3.

## Author contributions

CSM and BB conceived and designed the study and assisted with fieldwork and data collection. CSM conducted molecular work, data analyses, and wrote the manuscript with input from all authors. CMLP, BMM, MC, and MS assisted with data analyses and interpretation. RS and MJWS conducted fieldwork and sample collection. AO, EB, FG, HM, JRR, and FG contributed data.

## STAR METHODS

KEY RESOURCES TABLE

## RESOURCE AVAILABILITY

### Lead contact

Further information and requests for resources should be directed to and will be fulfilled by lead contact Barbara Block.

### Materials availability

This study did not generate new unique reagents.

### Data and code availability

All data generated in this paper will be made available through Dryad.

## EXPERIMENTAL MODEL AND SUBJECT DETAILS

We analyzed 82 Atlantic bluefin tuna (ABT) samples representing Gulf of Mexico (Gulf, n=43) and Mediterranean Sea (Med, n=41) populations (Table S1). Fin clips were obtained from adult ABT that were tagged and released with electronic tags between 2017 and 2022, as part of tagging programs directed by BAB (see Block et al. 1998 for additional details on tagging procedures). Electronic tags included in this study are pop-up satellite archival tags (PSAT) models miniPAT (Wildlife Computers, USA), PSATFLEX, PSATLIFE, and Archival-X (Lotek Wireless, Canada), and acoustic tag models V16-4H and V16-6H (Innovasea, USA). Fin clips were preserved in RNAlater and stored at −80°C. In addition to fin clips, we obtained position and environmental data from the electronic tags that allowed for spatial assignment to either the western or eastern ABT stocks, based on visitation to either the Gulf of Mexico or Mediterranean Sea spawning grounds. Larvae were provided from existing collections of JRR from the Gulf of Mexico, and AO and FG from the Mediterranean Sea.

## METHOD DETAILS

### Tagging and sample collection

Adult ABT were caught in waters off Canada (n=25), Ireland (n=11), Spain (n=2), and Israel (n=5). Full detailed summary of capture and tagging methods are described in Block et al. 1998 and Block et al. 2005. In summary, ABT were caught on rod and reel by permitted vessels using slightly different fishing techniques that are specific to each geographic region; generally using trolled lure/bait combos or live bait. Fish were brought onto the vessel where they are handled on a wet vinyl mat, gills irrigated with a high-pressure saltwater hose, and eye covered by a wet cloth. Pop-up satellite archival tags (PSATs) and external acoustic tags (hereafter simplified to “electronic tags”) were placed externally, using two 15 cm tethers with titanium darts that were inserted into the dorsal musculature next to the second dorsal fin of the animal. 19 individuals were outfitted with both a PSAT and an acoustic tag. While on the vessel, fish were measured for length (curved fork length; CFL) and half girth, and fin clips were collected and preserved in RNAlater® (Thermo Fisher Scientific, MA, USA).

Tag models placed in fish included in this study were: PSAT models miniPAT (Wildlife Computers, USA), Archival-X (Microwave Telemetry, USA), and PSATFLEX and PSATLIFE (Lotek, Canada), and acoustic models V16-4H and V-16-6H (Innovasea, USA).

### Geolocation methods

Movements of tagged ABT were reconstructed using a three-step method described in Wilson et al. (2015). To summarize, a Bayesian state space model (SSM) used light-level longitudes from Wildlife Computers’ Global Position Estimator v. 2 algorithm and sea surface temperature (SST)-derived latitudes, derived as described in Teo et al. (2004), as inputs. Two Markov chain Monte Carlo (MCMC) chains were run and 20,000 position estimates at 6 hr time steps were saved. The mean of the posterior distributions of both longitude and latitudes were used to generate the most probable tracks that are shown in Figure 1a, which were sub-sampled back to 24 hr for the analyses. Similarly to Ferter et al. (2024), we generated 99% likelihood surfaces from the posterior distribution for each estimated position (i.e., daily 99% likelihood surfaces rather than a single over the entire tag deployment from all estimated positions).

### DNA extraction and sequencing

We extracted genomic DNA using the DNeasy blood and tissue kit (Qiagen, CA, USA) following the manufacturer’s protocol. We quantified DNA concentrations using the Nanodrop Lite Spectrophotometer (Thermo Fisher Scientific, MA, USA). Libraries were prepared and sequenced at Texas A&M Genomics and Bioinformatics Service. 92 samples were sequenced in paired-end 2×150 bp mode on a single Illumina NextSeq 6000 lane to a depth of coverage of approximately 10x. Four samples (two from each population, one larva and one adult) were sequenced at approximately 45x depth of coverage.

### Mapping

We assessed library quality using FASTQC (Andrews, 2010) and performed sequence trimming and quality filtering using ADAPTERREMOVAL version 2.1.1 (Schubert et al. 2016). We trimmed low-quality bases and Ns from the ends of reads with a minimum quality threshold of 10. We mapped filtered reads to Haplotype 1 of the Atlantic bluefin tuna reference genome (Block et al. *in submission*) using bowtie2 version 2.5.1 (Langmead and Salzberg 2012) and subsequently used SAMTOOLS version 1.18 (Li et al. 2009) to convert SAM files and sort the resulting BAM files. We obtained alignment statistics using QUALIMAP version 2.2.2.

## QUANTIFICATION AND STATISTICAL ANALYSES

### Variant calling and data filtering

To prepare samples for variant calling, we used Picard Tools version 2.18.11 to add read groups, mark and remove duplicates, and index bam files. We called variants using GATK’s *Haplotype Caller* v. 3.8.1(McKenna et al. 2010) with the following parameters: QD < 2, FS > 60.0, MQ < 30.0, and ReadPosRankSum < −8.0. We additionally filtered out variants that were not biallelic, had minor allele frequencies less than 5%, mean coverage less than 2X or more than 50X, and more than 20% missing data using VCFTOOLS v. (Danecek et al. 2011). We pruned for linkage using PLINK v. 2.0 (Purcell et al. 2007) with a window size of 100kb, a step size of 100kb, and a variance inflation factor of 0.2.

### Sample quality filtering

We estimated relatedness among all samples using the “relatedness” statistic (Yang et al. 2010) in VCFTOOLS. We subsequently removed seven samples (Med larvae) that were closely related, either full (relatedness coefficient ≥0.5) or half (relatedness coefficient 0.2 – 0.35) siblings. We removed two samples (GULF larvae) for high heterozygosity (>0.5), and one sample (GULF tag) for inconsistencies with tag geolocation models. Removed samples were excluded from all analyses, leaving a final dataset of 82 individuals.

### Descriptive Statistics

We estimated basic descriptive statistics using the full SNP dataset and a putatively neutral SNP dataset, which was pruned to avoid linkage and exclude SNPs within 10kb of exons. For both datasets, we estimated nucleotide diversity (π), individual heterozygosity, and Tajima’s D for each population in 25 kb windows using VCFTOOLS (Danecek et al., 2011). To characterize genome-wide patterns of divergence between Gulf and Med populations, we calculated pairwise F*_ST_* values for individual SNPs and in 25 kb windows. To evaluate absolute genetic differentiation between the two populations, we estimated *d_xy_* in 25kb windows using PIXY version 1.2.7 (Korunes and Samuk, 2021). We compared heterozygosity in the Gulf vs. Med populations using the Mann-Whitney U test due to the non-normal distribution of the data and used Wilcoxon signed rank tests for comparing the mean heterozygosity and F*_ST_* of the full and neutral datasets. Differences were considered statistically significant at the *p* < 0.01 level. Data reported are mean ± SD unless otherwise specified.

### PCA and Admixture

To evaluate population structure between Gulf and Med ABT, we performed a principal component analysis (PCA) using the R package “SNPRelate” (Zheng et al. 2012) using both the full SNP dataset and the putatively neutral SNP dataset, (Full dataset: 11,181,223 SNPs, neutral dataset: 1,208,268 SNPs). To quantify the number of distinct genetic groups present in our dataset, we ran ADMIXTURE (Alexander et al. 2009) on the neutral dataset with the missing data threshold set to 90% (957,680 SNPs). ADMIXTURE estimates individual assuming the existence of distinct clusters, and we inferred the most parsimonious number of clusters by running ADMIXTURE assuming 1 to 6 clusters (K) and chose the best supported number of clusters using cross-validation (CV) error estimates.

### F_ST_-based Outlier Analysis

Peaks in *F*_ST_ were visualized using Manhattan plots constructed using the R package QQMAN (Turner, 2018). We defined outlier SNPs as those that exhibited elevated divergence between GULF and Med and, specifically, those that had *F*_ST_ estimates greater than four standard deviations above the weighted mean. For each outlier SNP, we extracted regions within within 40 kb of each SNP and used the R package GENOMICRANGES (Lawrence et al., 2013) to find overlapping annotations within the Atlantic bluefin tuna reference genome. We obtained information on the function of each gene using the UniProt database (http://www.uniprot.org/), and inspected individual outlier regions by plotting F*_ST_* with nucleotide diversity and Tajima’s D. Finally, we performed a PCA using just the outlier SNPs.

### GEMMA

We performed genome-wide associations to investigate whether differences in spawning timing, as well as oceanographic discrepancies between the Gulf and Med are associated with particular genomic regions. To do so, we used the Bayesian sparse linear mixed model (BSLMM) implemented in the package GEMMA (Zhou et al., 2013) to scan for SNPs associated with the behavioral trait of spawning timing, as well as oceanographic variables (detailed in section below). GEMMA detects SNPs that are associated with specified traits while controlling for population structure by incorporating a relatedness matrix as a covariate in the model.

We fit a linear BSLMM using MCMC with 5,000,000 burn-in iterations and 20,000,000 sampling iterations. Each SNP is assigned a posterior inclusion probability (PIP), which is the proportion of times a SNP has a non-zero effect on the trait of interest, obtained from the gamma values assigned to each SNP. We used a conservative threshold of PIP > 0.1 to estimate the significance of our genotype-behavior or genotype-environment associations following (Chaves et al., 2016; Armstrong et al., 2017).

### Oceanographic Data

SST, sea surface salinity (SSS), mixed layer depth (MLD), sea surface height (SSH), eastward velocity and northward velocity were obtained from the Global Ocean Physics Reanalysis (GLOBAL_MULTIYEAR_PHY_001_030) product (https://doi.org/10.48670/moi-00021) distributed by Copernicus Marine Service. The eastward and northward velocities were used to produce maps of eddy kinetic energy (EKE) with a spatial resolution of 0.083° × 0.083°.

On days where the posterior means of the longitude and latitude estimates were inside either of the two spawning grounds, the oceanographic variables above were extracted within the daily 99% likelihood surfaces and the mean value for each variable was computed. For each individual, the mean and standard deviation of the daily mean values were calculated and used in the GEMMA analyses described below.

### Demographic history

We used the PAIRWISE SEQUENTIALLY MARKOVIAN COALESCENT (PSMC) to estimate demographic history for each population (Li and Durbin, 2011). We followed the recommended data preparation for SNP calling using the mpileup command with a minimum mapping quality (“-q 30”) and minimum base quality (“-Q 30”) and generated a diploid sequence for each individual using “-c” in bcftools v.1.21 (Li, 2011). We ran PSMC using the default parameters, and for scaling, used a mutation rate of 3×10^-9^ and a generation length of 13 years. There are are a range of estimates for generation length of the two populations which ranges from 13-16 years for the Med and 13-19 for the GULF (Colette et al., 2021), so we selected to use 13 for both as to avoid potentially overestimating N_e_.

