## Supplemental Figures for "Ecological and evolutionary insights into the diversification of Atlantic bluefin tuna"

In preparation for *Current Biology*

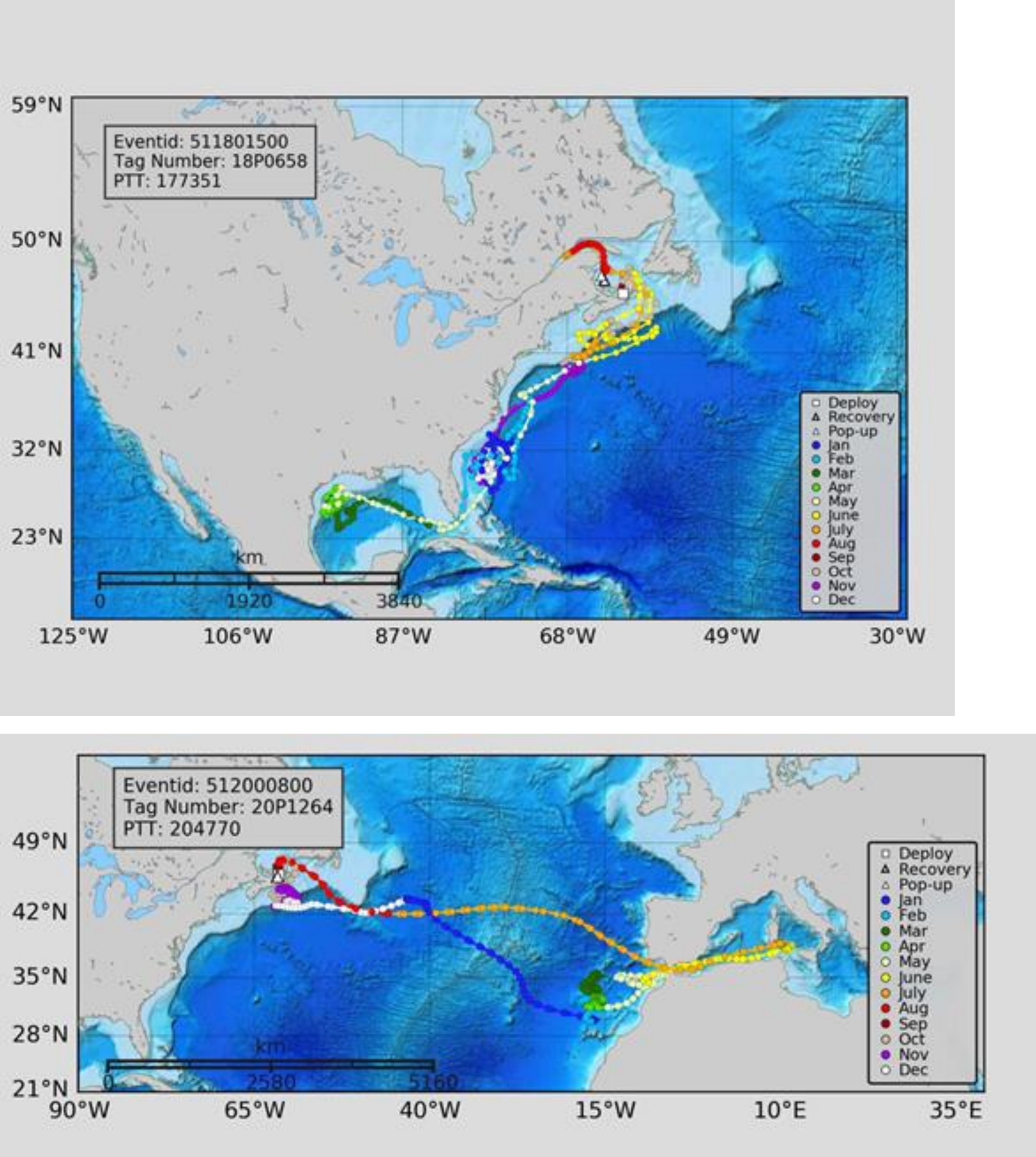


**Figure S1.** (Top panel): SSM track for Gulf-24. This individual was sequenced at high depth of coverage (58x). (Bottom panel) SSM track for Med-7. This individual was sequenced at high coverage (41.5x).

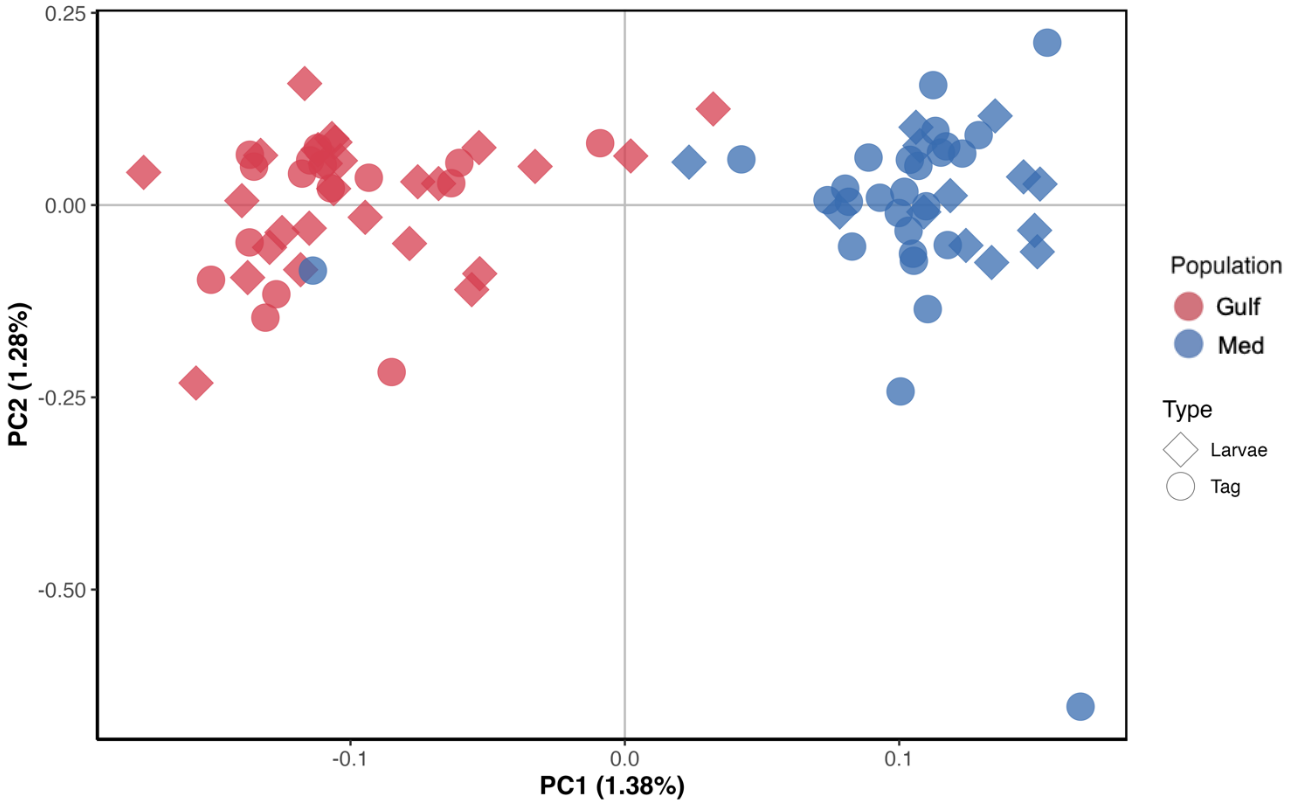


**Figure S2.** Principal component analysis of all samples using a putatively neutral dataset of 1,208,268 SNPs (dataset pruned for linkage and excluded SNPs mapped to or within 10kb of exons).

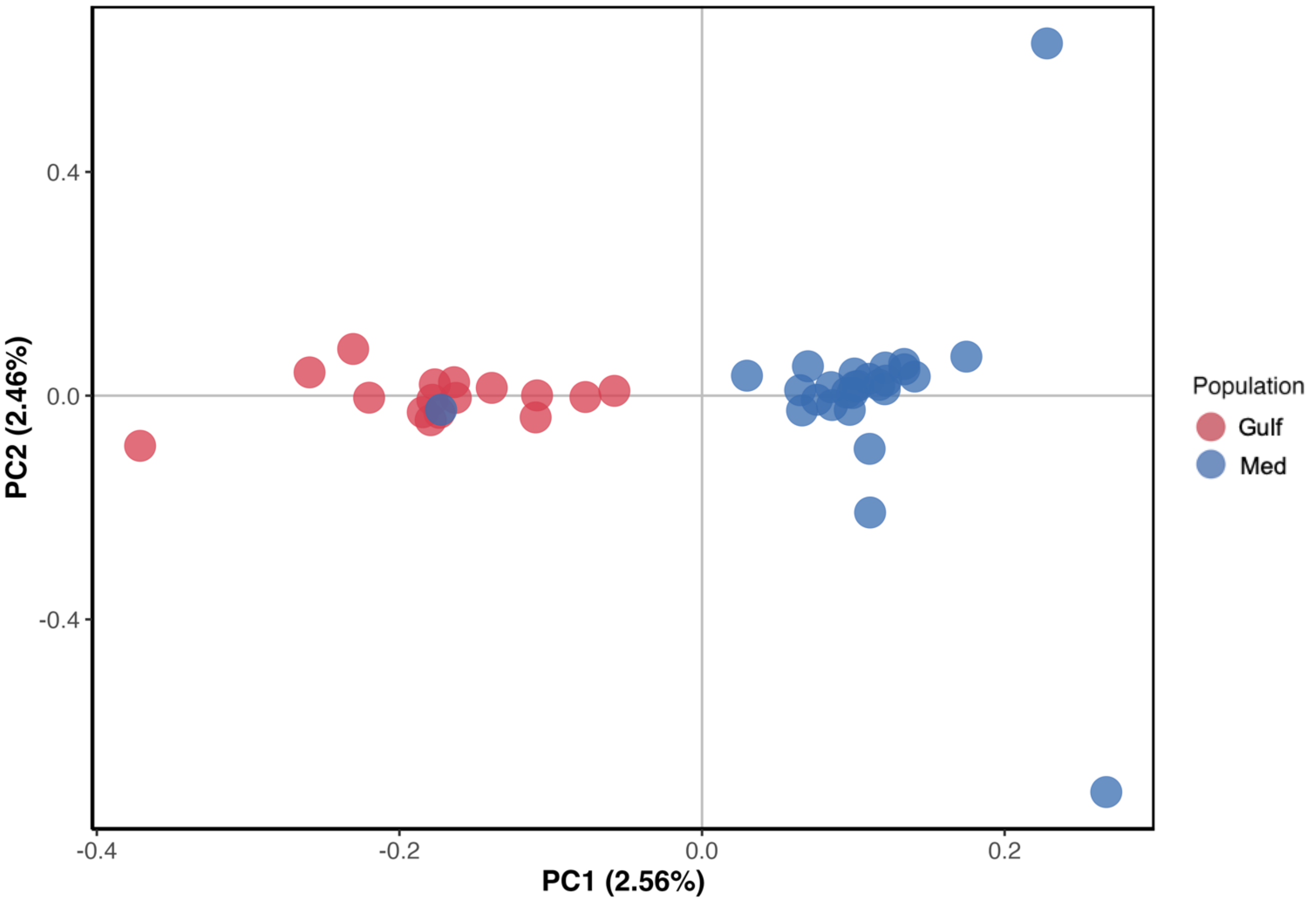


**Figure S3.** Principal component analysis of only tagged adult bluefin tuna using the full SNP dataset of 11,181,223 SNPs.

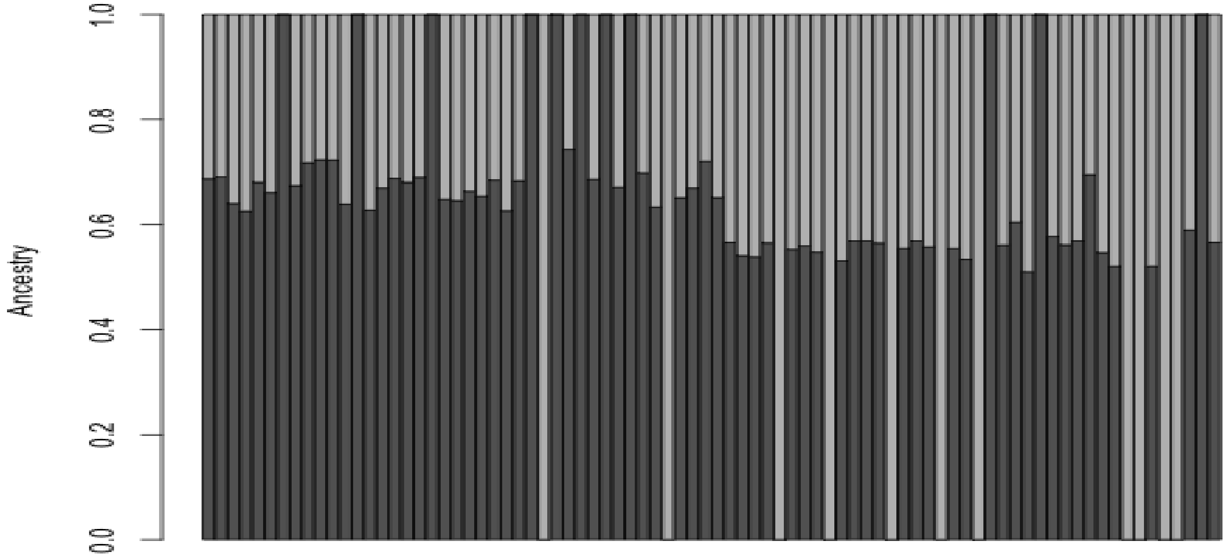


**Figure S3.** ADMIXTURE results using thinned neutral dataset with missing data threshold set to 90%.  Cross-validation error determined K=1 to be the most likely number of populations; we plotted K=2 for visual purposes.

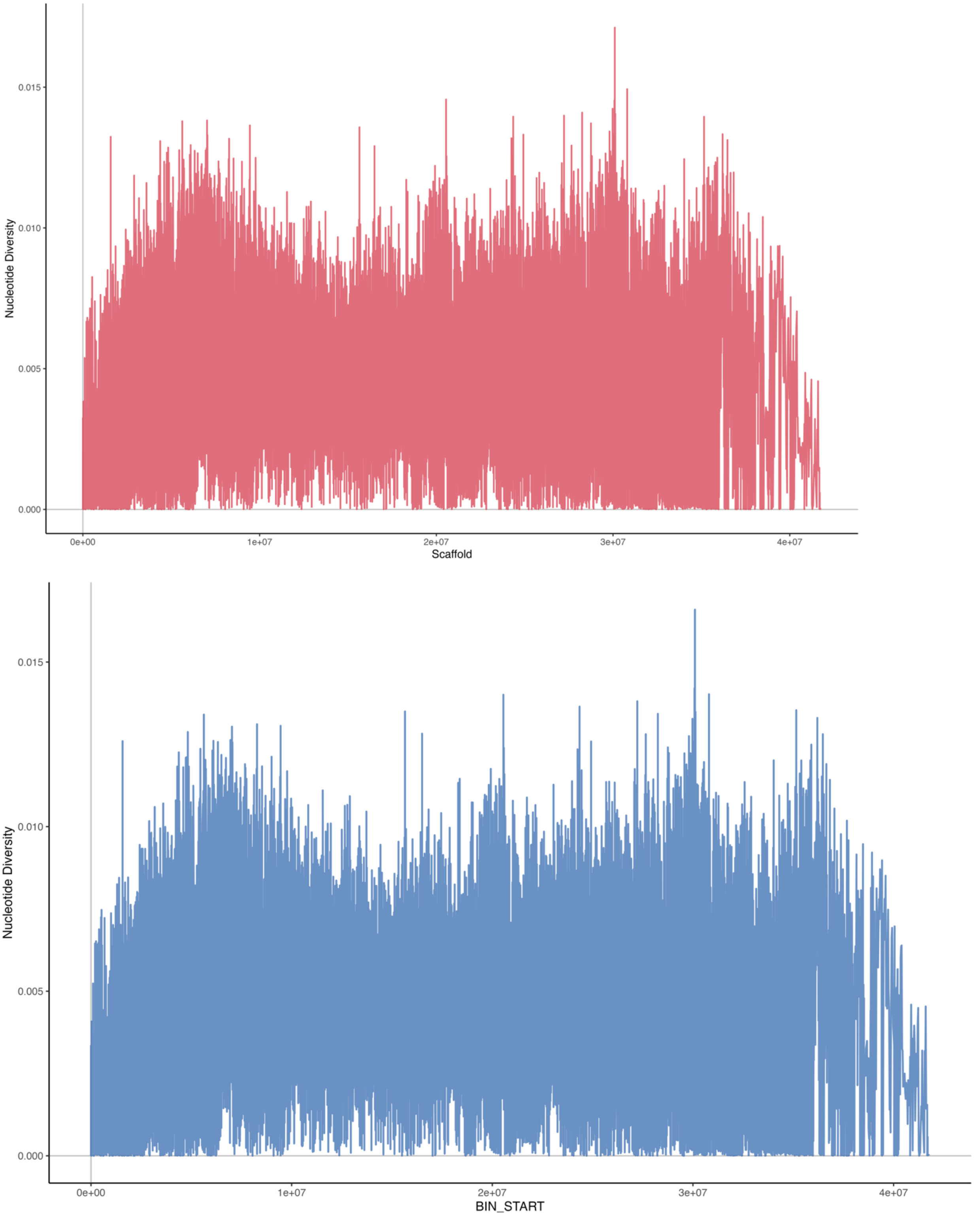


**Figure S5.** Genome-wide patterns of nucleotide diversity by population (Gulf= red, Med= blue).

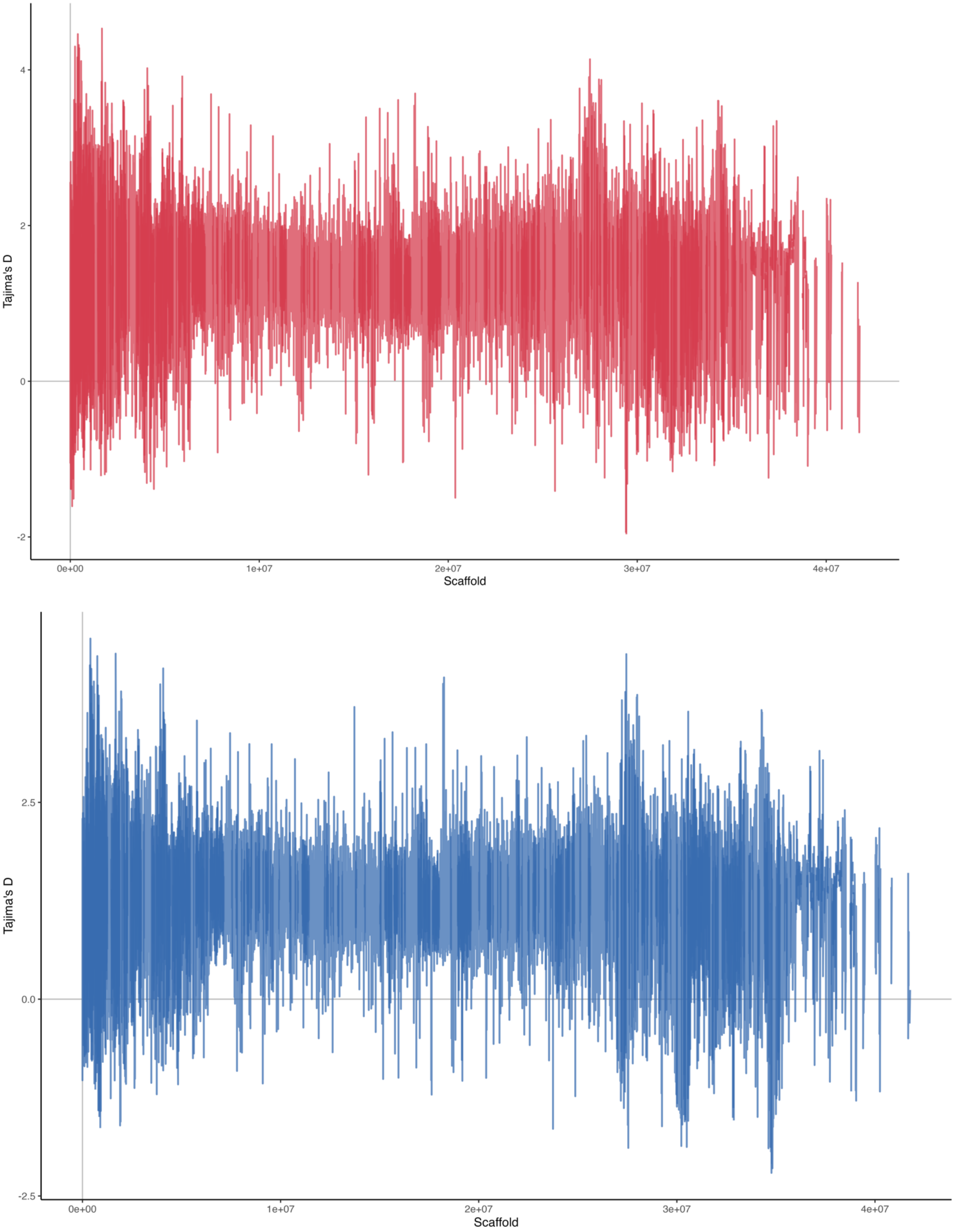


**Figure S6.** Genome-wide patterns of Tajima's D. (Gulf= red, Med= blue).

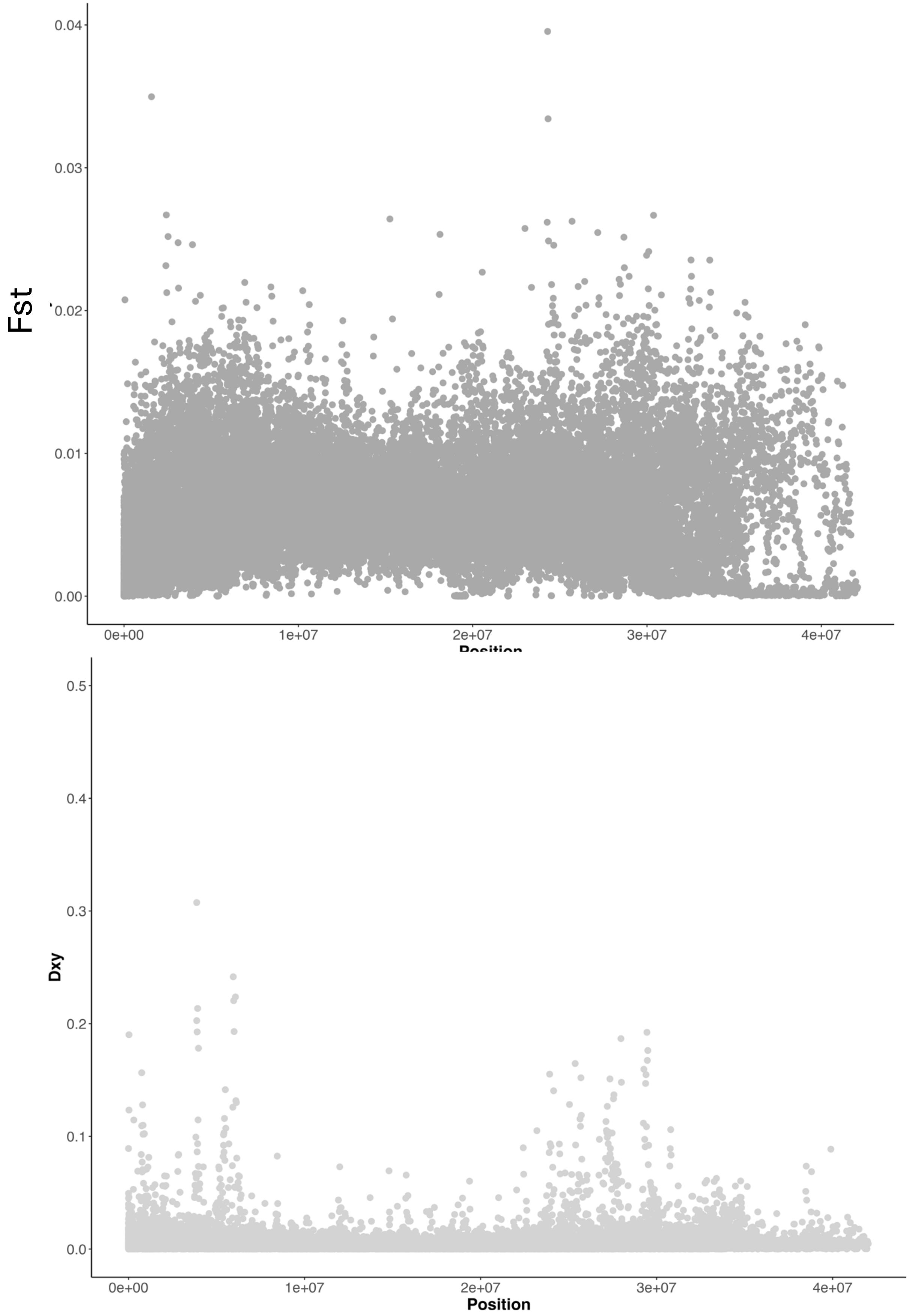


**Figure S7.** Genome-wide patterns of Fst (top) and Dxy (bottom).

Figure

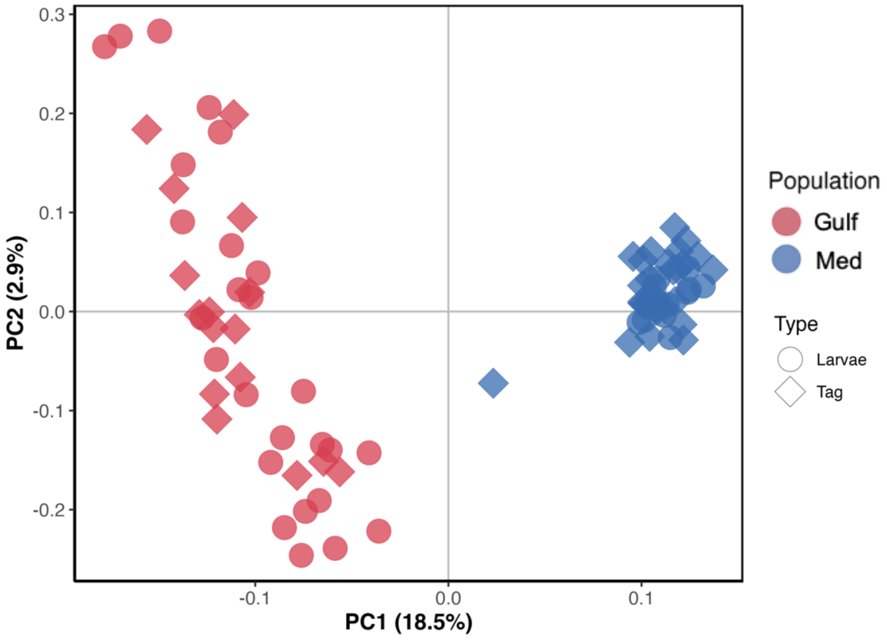


**Figure S8.** Principal component analysis using 298 outlier SNPs reveals clear separation of the two populations.

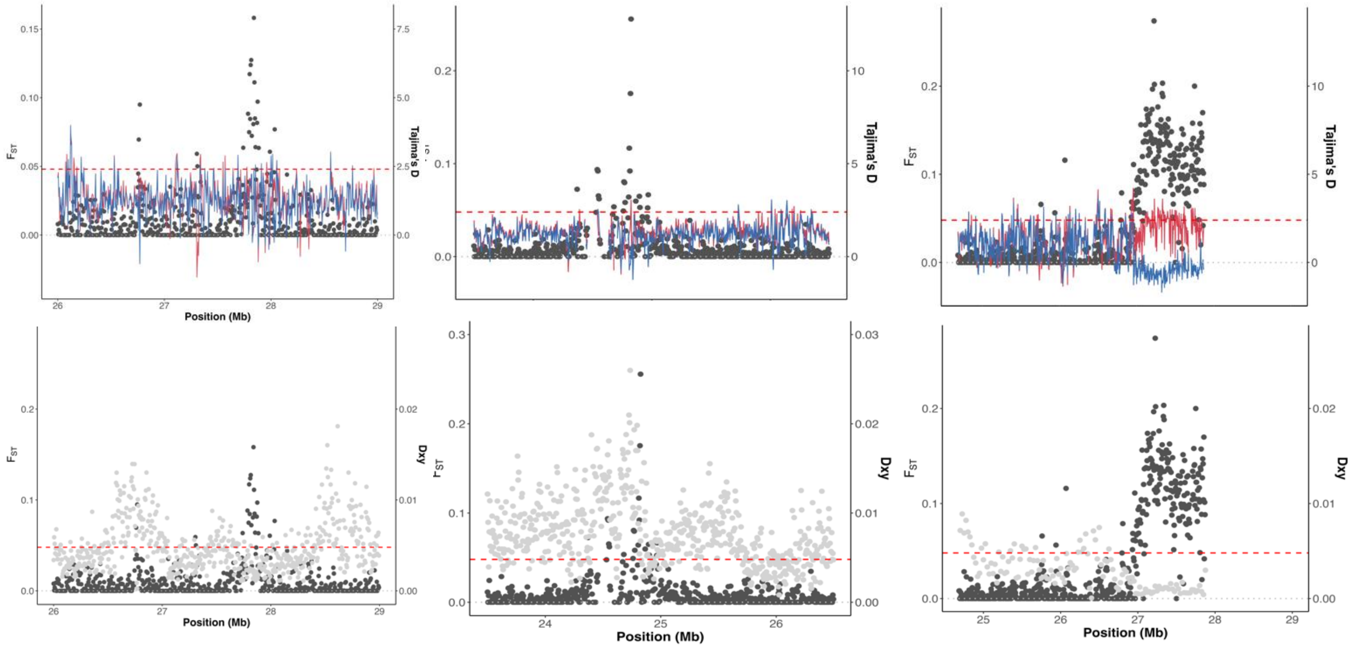


**Figure S9.** FST plotted in 5kb windows for zoomed in views of peaks on Chromosome 1, 8, and 21. Top panels: Fst plotted with nucleotide diversity by population (Gulf = red; Med = blue). Bottom panels: Fst plotted with Dxy (grey dots).

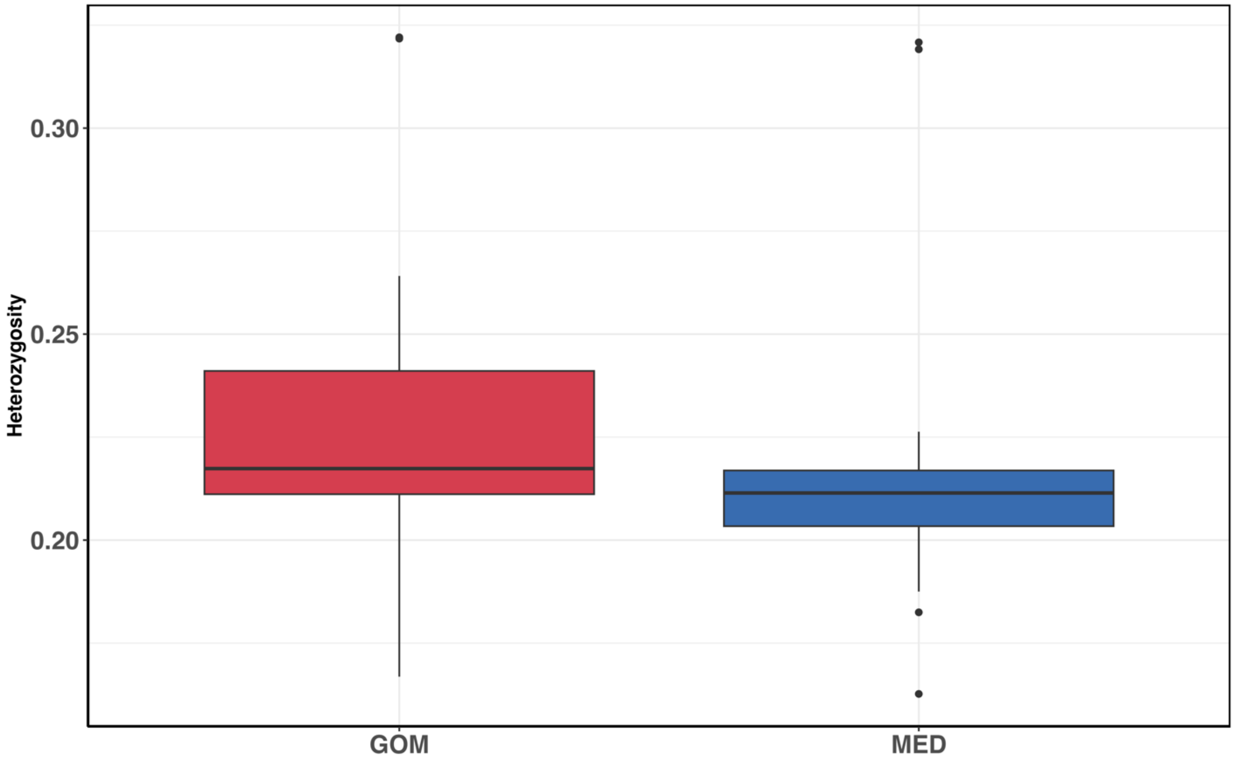


**Figure S10.** Individual heterozygosity for the neutral SNP dataset.

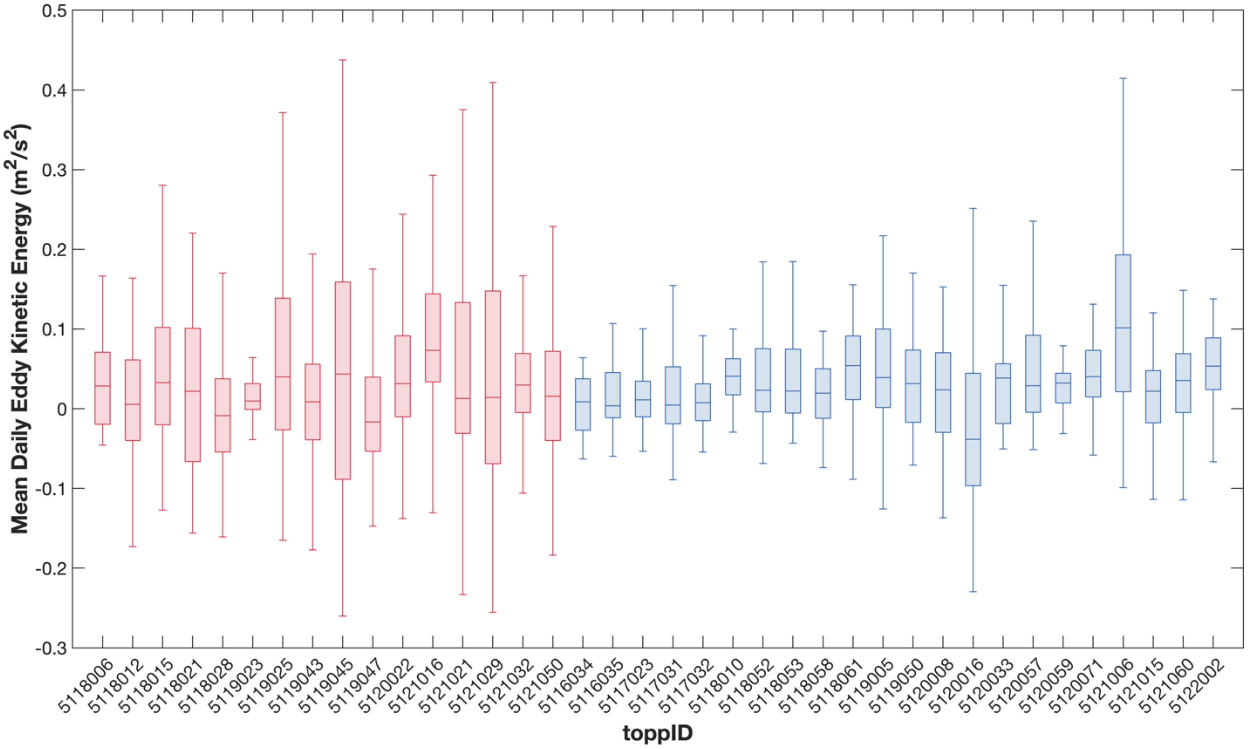


**Figure S11.** Mean daily eddy kinetic energy for adult bluefin tuna while occupying spawning grounds in Gulf of Mexico and Mediterranean Sea.

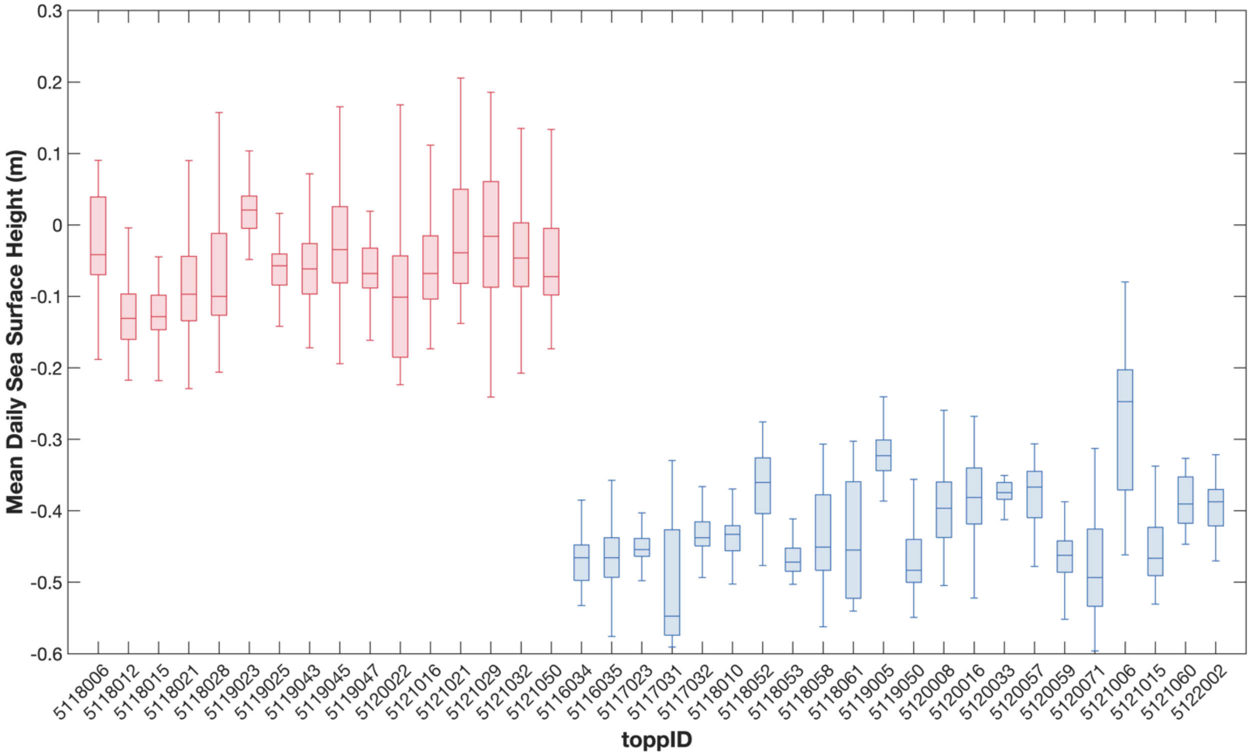


**Figure S12.** Mean daily sea surface height for adult bluefin tuna while occupying spawning grounds in Gulf of Mexico and Mediterranean Sea.

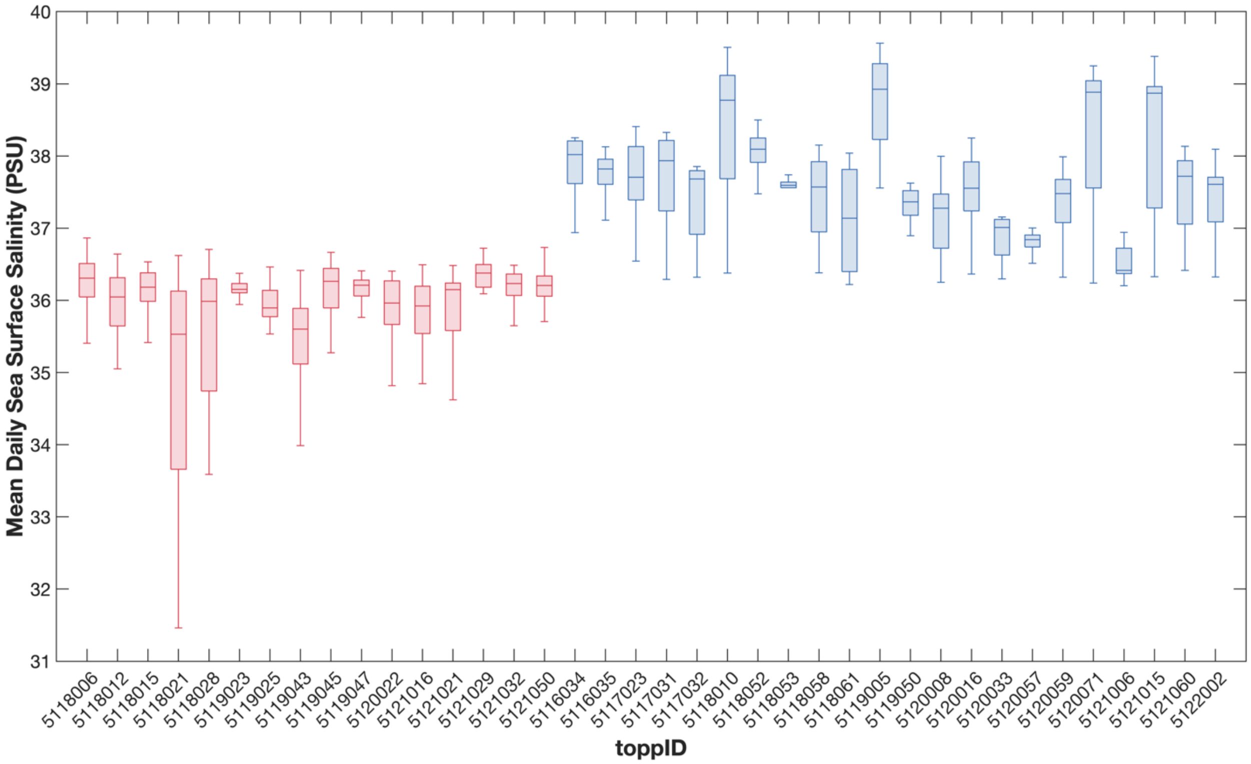


**Figure S13.** Mean daily sea surface salinity for adult bluefin tuna while occupying spawning grounds in Gulf of Mexico and Mediterranean Sea.

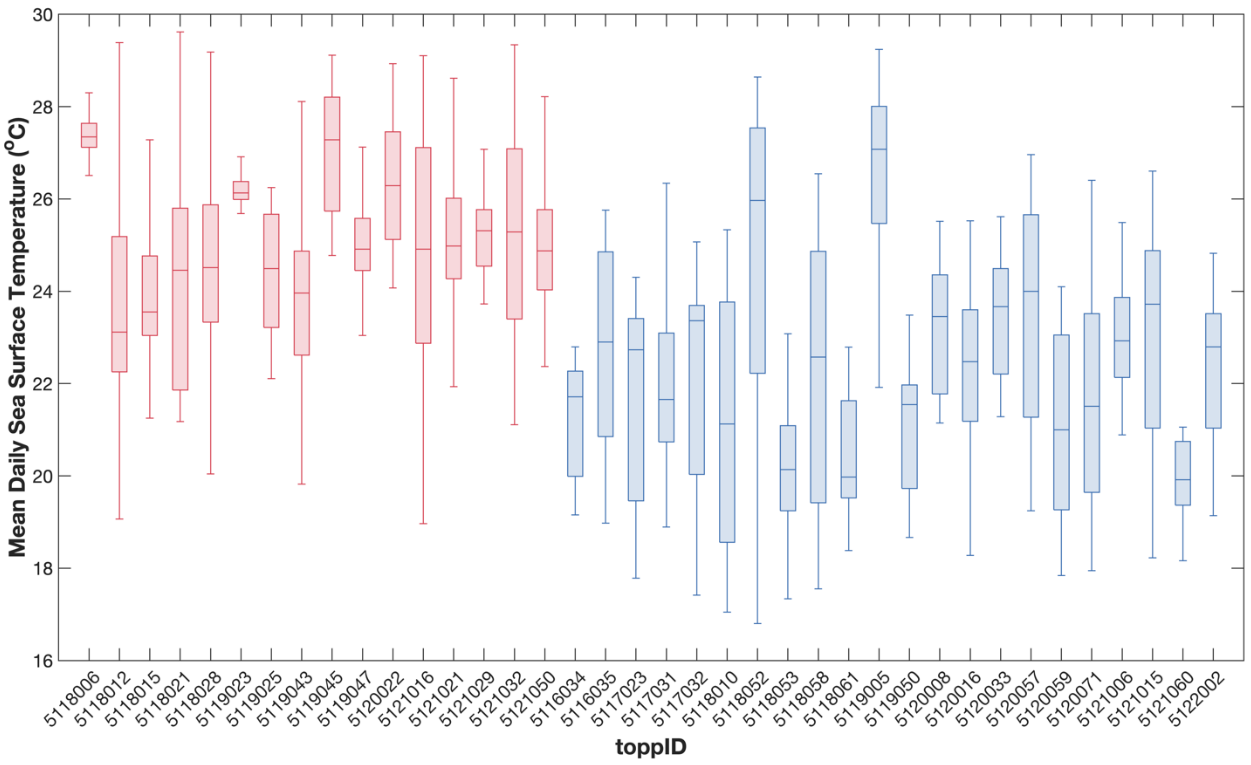


**Figure S14.** Mean daily sea surface temperature for adult bluefin tuna while occupying spawning grounds in Gulf of Mexico and Mediterranean Sea.

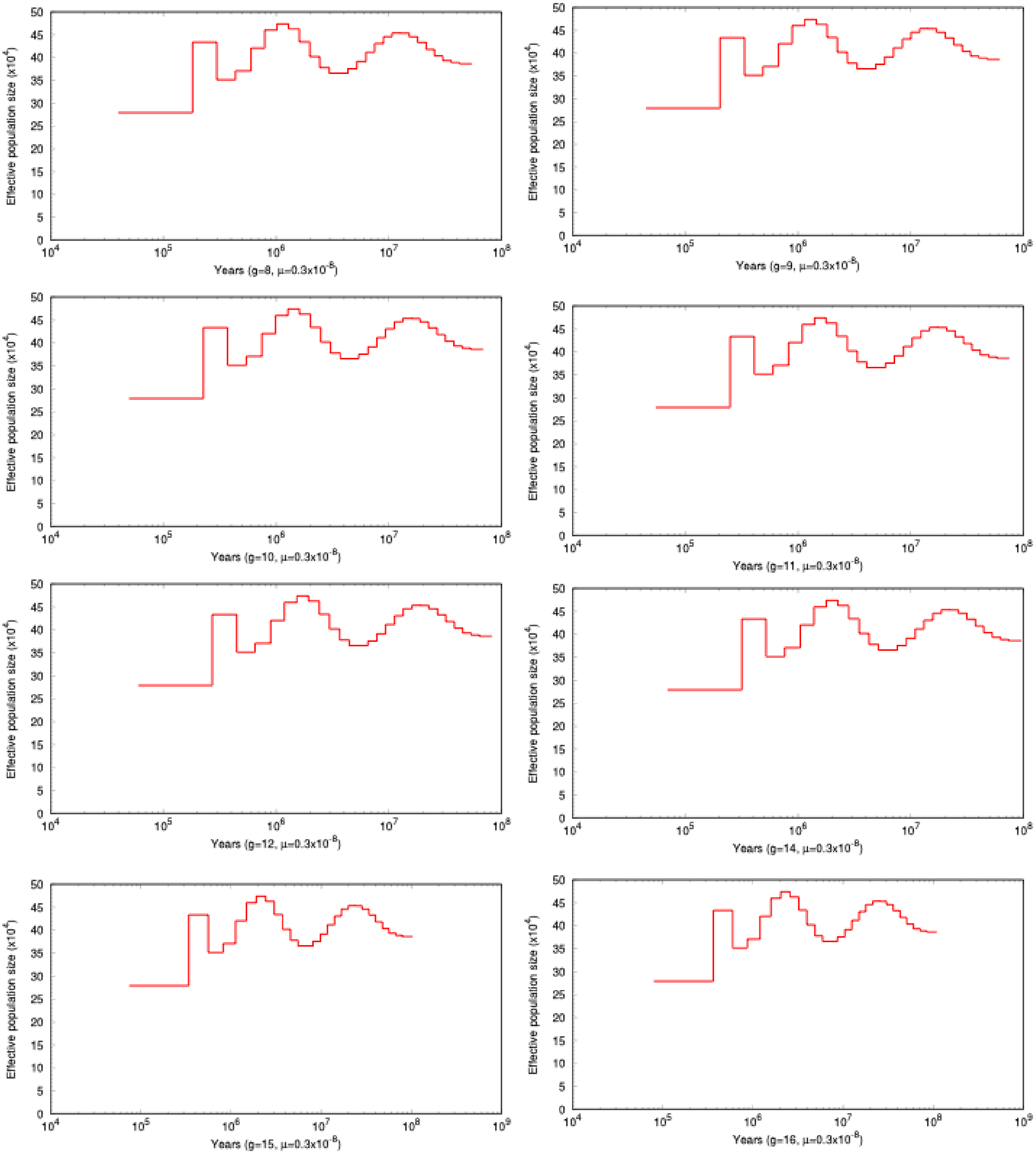


**Figure S15.** PSMC runs scaled for generation lengths of 8 – 16, for individual Med-7.

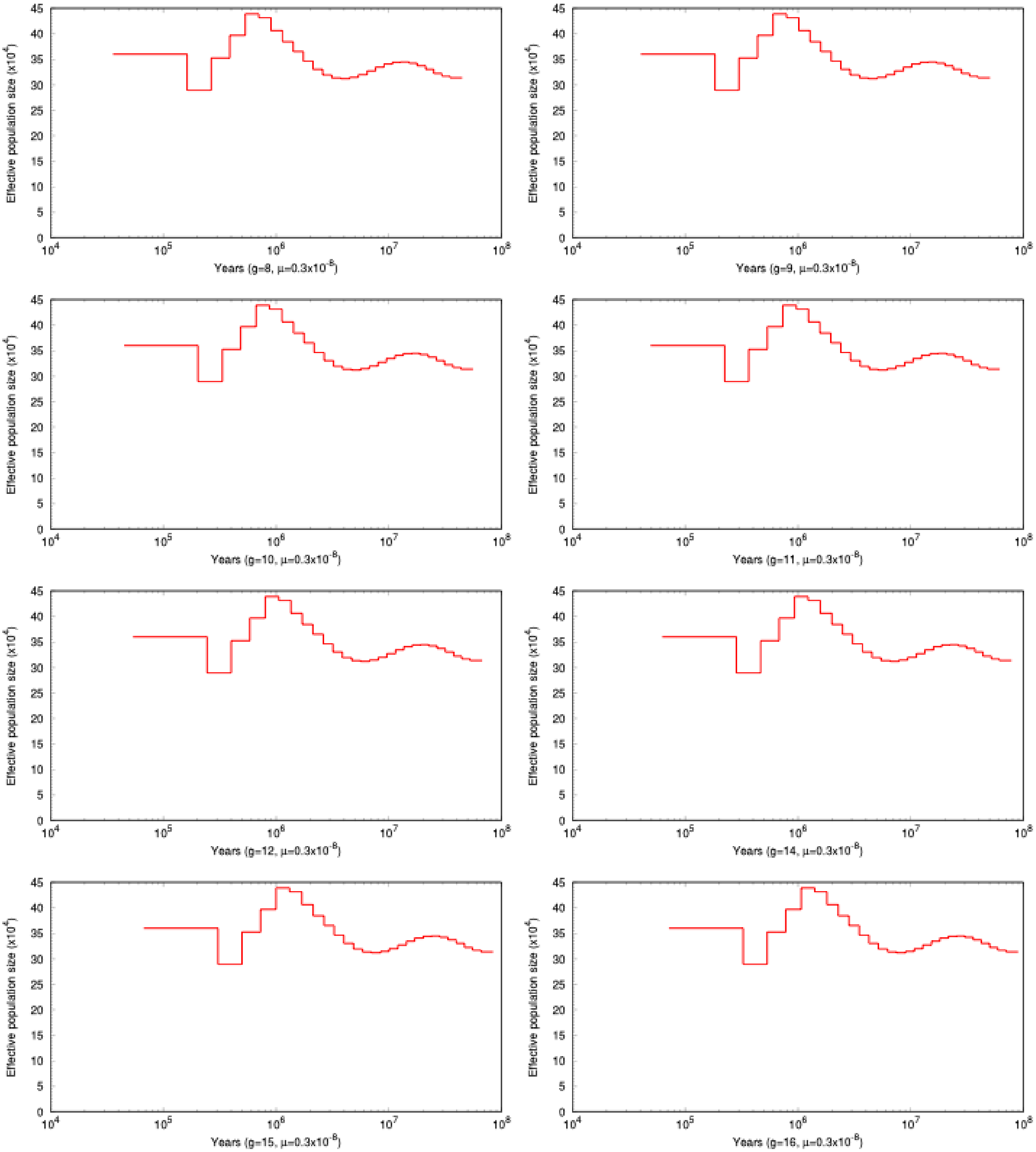


**Figure S15.** PSMC runs scaled for generation lengths of 8 – 16, for individual Gulf-24.
