## Supplemental Table 1 for "Ecological and evolutionary insights into the diversification of Atlantic bluefin tuna"

| ID | Population | Type | Tagcode | Barcode | Alignment Percentage | Mean coverage | Std. coverage | Missing Data | Heterozygosity | pc1 | pc2 | pc3 |
|---|---|---|---|---|---|---|---|---|---|---|---|---|
| Gulf-10 | Gulf | Larvae | NA | GTGCGCATTG+TGTCCACTGG | 99.56 | 9.8886 | 5.4181 | 0.0329076 | 0.181684416 | -0.1199456 | -0.0043971 | -0.0071344 |
| Gulf-11 | Gulf | Larvae | NA | GCGGTTTAGC+AGTAGCGTGG | 99.49 | 8.1279 | 4.5452 | 0.042914 | 0.167906386 | -0.1061994 | -0.0388062 | 0.07448653 |
| Gulf-12 | Gulf | Larvae | NA | AACCGGTAGC+CACTCCTTTC | 99.49 | 9.2513 | 5.4087 | 0.032998 | 0.181198912 | -0.0598532 | 0.02658885 | -0.0289534 |
| Gulf-13 | Gulf | Larvae | NA | ACTGCTCATG+ATTGTGAGGA | 99.28 | 8.4764 | 4.7523 | 0.0381494 | 0.173238088 | 0.01052908 | 0.04139599 | -0.0268424 |
| Gulf-14 | Gulf | Larvae | NA | ATAGTAACAC+GCATCCCAGT | 99.33 | 9.187 | 4.9306 | 0.0339738 | 0.178860873 | -0.1160893 | -0.0264238 | -0.0181352 |
| Gulf-19 | Gulf | Tag | 19P0507 | CCTAGTGCAG+ACGTGCAATT | 99.51 | 12.1309 | 6.1415 | 0.0148819 | 0.210059039 | -0.0972295 | 0.00375479 | -0.044518 |
| Gulf-2 | Gulf | Tag | 20P0957 | TCCATGACGG+AACGTCTCCG | 99.46 | 7.5485 | 5.4455 | 0.0893255 | 0.1321647 | -0.0612836 | -0.0070388 | 0.0775444 |
| Gulf-20 | Gulf | Tag | 19P0504 | TTAACCGCAA+TGTATCCTAA | 99.53 | 11.5443 | 6.135 | 0.0191108 | 0.202034697 | -0.1242903 | -0.004255 | -0.0106902 |
| Gulf-21 | Gulf | Tag | 19P0547 | AACTGATCCG+TATTAACCTA | 99.53 | 10.5225 | 5.5584 | 0.0237105 | 0.194886912 | -0.1353557 | -0.0319974 | -0.0515352 |
| Gulf-22 | Gulf | Tag | L340-2683 | TAATTGAGGT+GCTGCAAGTG | 99.47 | 8.042 | 4.5231 | 0.0478456 | 0.165543491 | -0.1776216 | 0.04599868 | 0.10860167 |
| Gulf-23 | Gulf | Tag | 19P0500 | CGCCATCTAT+GAACCTCGTC | 99.48 | 8.5872 | 4.8913 | 0.0430823 | 0.170601235 | -0.1233706 | 0.03459125 | 0.00702775 |
| Gulf-24 | Gulf | Tag | 18P0658 | TCCCTTCGTC+CCCAATACCA | 99.49 | 58.1679 | 24.3781 | 0.00159758 | 0.241083772 | -0.0645074 | 0.01862044 | -0.0232367 |
| Gulf-25 | Gulf | Tag | 18P0647 | GTTTATATCG+AGCCGGACCA | 99.54 | 8.3144 | 4.6982 | 0.0432563 | 0.170717778 | -0.0993712 | -0.1549084 | 0.1486155 |
| Gulf-26 | Gulf | Tag | 14P0401 | GTCCGGTTAC+AACCGTGGAT | 99.55 | 8.947 | 4.9334 | 0.0364411 | 0.177262411 | -0.0223364 | 0.01911485 | -0.0583446 |
| Gulf-27 | Gulf | Tag | L330-3454 | GATGAACTAC+ATGCACGTGG | 99.58 | 10.2906 | 6.0736 | 0.0301463 | 0.189301687 | -0.1317813 | 0.01556477 | 0.01088079 |
| Gulf-28 | Gulf | Tag | 21P0408 | CAGAAATCGC+TACAATTTGT | 99.54 | 9.1391 | 5.0714 | 0.0326307 | 0.18210143 | -0.1190189 | 0.04636515 | -0.009339 |
| Gulf-29 | Gulf | Tag | 21P0400 | GCAATAGCTA+CATCCCTTAG | 99.69 | 5.4174 | 8.021 | 0.0296392 | 0.194490887 | -0.1516308 | 0.01142798 | -0.0961446 |
| Gulf-3 | Gulf | Tag | 21P0348 | AAGTGACTTG+CATAAACGTC | 99.4 | 8.5356 | 5.5529 | 0.041728 | 0.171645703 | -0.1093419 | -0.0129006 | -0.0145773 |
| Gulf-30 | Gulf | Tag | 18P0624 | AGCACGCAGG+ACTCATTCCG | 99.54 | 8.7165 | 4.8929 | 0.04086 | 0.174591313 | -0.1285448 | -0.0216591 | 0.0368137 |
| Gulf-31 | Gulf | Tag | 18P0628 | AAGAGCTTAT+TAGTCGCTTT | 99.51 | 11.3134 | 6.3903 | 0.0244249 | 0.193097342 | -0.0679747 | 0.00793823 | -0.0361551 |
| Gulf-32 | Gulf | Larvae | NA | ACGAGCCAAA+CAGCATAATC | 99.45 | 59.0367 | 22.3581 | 0.00143784 | 0.240876094 | -0.0867102 | 0.00012494 | -0.0160225 |
| Gulf-33 | Gulf | Larvae | NA | TTTATAGATC+GCGCATTCTC | 99.23 | 9.9849 | 5.1935 | 0.0229641 | 0.193847415 | -0.1133341 | -0.0086384 | -0.033747 |
| Gulf-34 | Gulf | Larvae | NA | TGGTGGTTTC+TCTGCATGCA | 99.6 | 11.0765 | 5.5984 | 0.0166915 | 0.205389588 | -0.0927163 | 0.00438068 | 0.00145707 |
| Gulf-35 | Gulf | Larvae | NA | AGGAATCGCT+TAGACGGTAA | 99.52 | 11.3536 | 5.945 | 0.0185714 | 0.201926991 | -0.1311876 | -0.003164 | -0.0497078 |
| Gulf-36 | Gulf | Larvae | NA | GAATGTATGT+AACTCTGTTC | 99.52 | 11.3188 | 5.8129 | 0.0189367 | 0.203037187 | -0.0330928 | -0.0020555 | -0.0220082 |
| Gulf-37 | Gulf | Larvae | NA | TCTTCCAGTC+ATAAACTCCG | 99.57 | 9.9779 | 5.452 | 0.0290449 | 0.186532272 | -0.1084618 | 0.02895083 | -0.0210385 |
| Gulf-38 | Gulf | Larvae | NA | CATCCTGTAA+TTCTTAGGTC | 99.43 | 9.0037 | 5.0296 | 0.0338657 | 0.17860017 | -0.1023555 | -0.0227316 | -0.0282124 |
| Gulf-39 | Gulf | Larvae | NA | AATGGATTGC+CAGACCGCGG | 99.49 | 7.259 | 4.165 | 0.0562886 | 0.157019284 | -0.1727083 | 0.09424012 | 0.2462419 |
| Gulf-4 | Gulf | Tag | 21P0344 | CCCGCCTGAA+CAGTGGTCGG | 99.47 | 8.289 | 4.9352 | 0.0371011 | 0.174748728 | -0.1161831 | 0.00272557 | 0.00561863 |
| Gulf-40 | Gulf | Larvae | NA | GACCGCTCAC+AGGGAATCAT | 99.45 | 8.0253 | 4.4147 | 0.0449247 | 0.166493509 | -0.1016171 | 0.00805545 | 0.03096416 |
| Gulf-41 | Gulf | Larvae | NA | CCTCGTCACT+GACGACGCTC | 98.06 | 8.4439 | 4.7414 | 0.0415254 | 0.170200238 | -0.1234165 | 0.01307765 | -0.0093309 |
| Gulf-42 | Gulf | Larvae | NA | AGTCACGCCA+TAGGGCAGTC | 99.46 | 8.6742 | 4.8281 | 0.0383392 | 0.173220414 | -0.1468068 | 0.00826195 | -0.010428 |
| Gulf-43 | Gulf | Larvae | NA | TGTAGGATGG+CAAGTTTGTT | 99.59 | 8.0538 | 4.6135 | 0.044911 | 0.166633803 | -0.1026777 | -0.0032165 | -0.0469418 |
| Gulf-45 | Gulf | Larvae | NA | GCGAATGATA+CTTATGAACG | 98.98 | 8.4702 | 4.7115 | 0.0386762 | 0.173515361 | -0.1100718 | 0.02527984 | -0.041753 |
| Gulf-46 | Gulf | Larvae | NA | GCAAGGAACA+ACAGGTAGAT | 99.53 | 8.4005 | 4.841 | 0.0395433 | 0.17243941 | -0.14957 | 0.00774074 | -0.0029537 |
| Gulf-47 | Gulf | Larvae | NA | CGCATCCCGA+CTGTAGCCCA | 99.6 | 8.4312 | 4.8569 | 0.038063 | 0.175238098 | -0.1100132 | 0.04106102 | -0.046357 |
| Gulf-49 | Gulf | Larvae | NA | GAATATTAAT+AAGTAGGGTC | 99.58 | 7.6946 | 4.5154 | 0.0478366 | 0.165829601 | 0.01636692 | -0.4165938 | 0.32996539 |
| Gulf-5 | Gulf | Larvae | NA | GAGATCTATG+AGGTCGCGCG | 99.07 | 8.0419 | 4.4458 | 0.0415307 | 0.170871327 | -0.0593933 | -0.0352891 | 0.00681448 |
| Gulf-6 | Gulf | Larvae | NA | AAACCAATAC+CATAATTTCA | 99.15 | 8.2926 | 4.6213 | 0.0405547 | 0.171941755 | -0.1149688 | 0.00427371 | -0.0441549 |
| Gulf-7 | Gulf | Larvae | NA | TGATCACGTA+TACCGTCCCA | 99.49 | 9.0225 | 5.2606 | 0.0439049 | 0.16839907 | -0.0830852 | 0.01471609 | 0.01087195 |
| Gulf-8 | Gulf | Larvae | NA | GATGTGAGAT+TATATTTAGG | 99.54 | 8.2184 | 4.6769 | 0.0457886 | 0.166573046 | -0.0679511 | -0.0552173 | 0.07268124 |
| Gulf-9 | Gulf | Larvae | NA | TGGTTGGGTC+GCATGTCCTT | 99.56 | 8.0661 | 4.5904 | 0.0456237 | 0.16662607 | -0.051919 | 0.00340554 | 0.02218411 |
| Med-1 | Med | Tag | 21P1998 | ACAGGCACAA+ACTTATTAAT | 99.51 | 8.0618 | 5.2562 | 0.0529755 | 0.163695373 | 0.09436145 | 0.04317747 | -0.037614 |
| Med-10 | Med | Tag | 18P0893 | TCGGTGGAAC+GTGCAGTATA | 99.54 | 7.7141 | 4.6384 | 0.0511185 | 0.162801693 | 0.11994635 | -0.0388067 | -0.0126763 |
| Med-11 | Med | Tag | 17P0217 | TATATGTCTA+TGGGATTCCA | 99.43 | 8.4749 | 4.8189 | 0.0367332 | 0.176556525 | 0.0987081 | 0.04244439 | -0.0340505 |
| Med-12 | Med | Tag | L330-2973 | CACATACGGA+GTGATTGAAC | 99.56 | 8.2803 | 4.738 | 0.0376956 | 0.176259368 | 0.11009454 | -0.0386653 | 0.01934112 |
| Med-13 | Med | Tag | 20P1296 | TCACAACGTC+TACTCATGGG | 99.51 | 8.2015 | 4.6965 | 0.042525 | 0.171869951 | 0.13226505 | -0.1704481 | 0.16427179 |
| Med-14 | Med | Tag | 20P1340 | GTGGAGCGCT+GTTCAACAAA | 99.47 | 8.5625 | 4.8091 | 0.0369875 | 0.175481126 | 0.10036595 | 0.016649 | -0.0619148 |
| Med-15 | Med | Tag | 18P1500 | GATGAGCGAA+AACGCGGATT | 99.55 | 8.1891 | 4.6714 | 0.0443821 | 0.170636584 | 0.1858426 | -0.3809826 | 0.0738412 |
| Med-16 | Med | Tag | 18P0557 | TTCTCCTATC+CGTACATGCA | 99.52 | 8.4924 | 4.9522 | 0.0394182 | 0.175530284 | 0.12102217 | -0.0549478 | 0.02630011 |
| Med-17 | Med | Tag | 18P0555 | AATTAGATTG+GCATTTAGCG | 99.5 | 8.6066 | 5.1507 | 0.0417095 | 0.173769436 | 0.12768332 | 0.1503718 | -0.3042817 |
| Med-18 | Med | Tag | 18P0547 | TAGAGACCCA+CCGCTGATGG | 99.56 | 8.0684 | 4.5692 | 0.0450638 | 0.168413983 | 0.10749527 | -0.0115188 | -0.0089581 |
| Med-19 | Med | Tag | 18P0560 | GATACTTAGG+ATCCTGCATT | 99.55 | 8.4995 | 4.9178 | 0.0414808 | 0.172693485 | 0.08913712 | -0.0028632 | 0.0028301 |
| Med-2 | Med | Tag | 20P2947 | CTTTCGTCTT+TTCGGTTATT | 99.59 | 8.2152 | 4.8779 | 0.0473553 | 0.168820502 | 0.11037866 | -0.0704965 | 0.04910763 |
| Med-23 | Med | Larvae | NA | CGGTATGGCT+TCGTGTGGGT | 99.4 | 8.2672 | 5.033 | 0.0455595 | 0.167014362 | 0.09415575 | -0.0888815 | 0.02459826 |
| Med-24 | Med | Larvae | NA | AGGGCAATGA+CAAGGATTCT | 99.5 | 7.0635 | 4.9409 | 0.0966442 | 0.131959783 | 0.09510625 | 0.01418708 | -0.0323852 |
| Med-26 | Med | Larvae | NA | TTCTCCTCGA+GTCTGTTTAA | 99.46 | 8.5681 | 5.2332 | 0.0390665 | 0.174854225 | 0.10472478 | 0.00780104 | -0.0902595 |
| Med-27 | Med | Larvae | NA | GTTGTCATTT+AAATCTCTTA | 99.41 | 8.49 | 4.9067 | 0.0414332 | 0.170617805 | 0.11113996 | -0.0020449 | -0.028978 |
| Med-29 | Med | Larvae | NA | AAGTGGCCAC+TATGGGACTC | 99.39 | 8.9542 | 5.2618 | 0.0365414 | 0.176199164 | 0.11243711 | -0.0729658 | 0.06955153 |
| Med-3 | Med | Tag | L330-3482 | ACCTTTGGGC+CACCTCTCTT | 99.43 | 7.8601 | 4.5243 | 0.0492544 | 0.163392693 | 0.10634282 | 0.06447049 | -0.0727799 |
| Med-30 | Med | Larvae | NA | ACCTACAGAC+CACAGTCATG | 99.41 | 8.4072 | 5.3777 | 0.0448251 | 0.168144443 | 0.11587496 | -0.0686457 | -0.0178833 |
| Med-31 | Med | Larvae | NA | TGCCGCCCGC+ACGGCAGGAT | 99.35 | 8.7008 | 5.2359 | 0.0377332 | 0.173695975 | 0.11265012 | 0.02565727 | -0.055708 |
| Med-32 | Med | Larvae | NA | TACAGAGAAG+ACACCAACTG | 99.26 | 7.4094 | 4.5535 | 0.0631378 | 0.149847196 | 0.10248944 | 0.01848892 | -0.0079697 |
| Med-34 | Med | Larvae | NA | GCGCTATCAA+ACTATTCATG | 99.42 | 8.0562 | 4.8135 | 0.0519594 | 0.158452044 | 0.15935694 | -0.2271635 | 0.11965525 |
| Med-36 | Med | Larvae | NA | TCATCGTGGT+GACCAGCATC | 99.27 | 8.5275 | 5.1064 | 0.0415559 | 0.170110208 | 0.11192349 | 0.02960992 | -0.0530291 |
| Med-37 | Med | Larvae | NA | TGAAACGACA+AGTTGAGGTC | 99.56 | 40.4675 | 15.3404 | 0.00158638 | 0.241084877 | 0.0399432 | -0.0007496 | -0.0462743 |
| Med-38 | Med | Larvae | NA | TAATACGTTG+TTTGAGGGCG | 99.56 | 7.2728 | 4.3254 | 0.0591285 | 0.156547588 | 0.12358628 | 0.02894128 | -0.0759624 |
| Med-39 | Med | Larvae | NA | CCAAGGAAAT+AACCGATTAG | 99.47 | 8.477 | 4.9306 | 0.0390748 | 0.174480293 | 0.10527082 | -0.0089277 | -0.0272814 |
| Med-4 | Med | Tag | 21P0405 | GATCGACTAT+ATAGCAGGTG | 99.51 | 8.3612 | 5.0306 | 0.0396584 | 0.173479459 | 0.11474543 | 0.05155481 | -0.0950648 |
| Med-40 | Med | Tag | 36213 | GCCGTGAGCT+ATGCCCTTGT | 99.51 | 8.8891 | 4.8358 | 0.033249 | 0.180170462 | 0.08896293 | -0.0115811 | -0.069835 |
| Med-41 | Med | Tag | 18P0886 | AGGATCGGGC+CTGAAGGTAA | 99.24 | 8.5097 | 4.9043 | 0.0411416 | 0.171859457 | 0.11507222 | 0.17265957 | -0.3105998 |
| Med-42 | Med | Tag | 14P0396 | CACTCAGAAA+TAACACAGCT | 99.51 | 8.1215 | 5.0297 | 0.0433727 | 0.169884302 | -0.1184764 | 0.03391955 | -0.0123509 |
| Med-43 | Med | Tag | 13P0073 | GCTTCGCAAC+GTGCCAGGTC | 99.57 | 7.9608 | 4.6871 | 0.0557437 | 0.163895871 | 0.08952435 | 0.0291694 | -0.0817752 |
| Med-44 | Med | Tag | 13P0076 | TATAATTGTA+ATCAGTCTAC | 99.49 | 8.1289 | 4.6163 | 0.0448869 | 0.168067115 | 0.08933264 | 0.03804069 | -0.0207329 |
| Med-45 | Med | Tag | 1275516 | ACGTGACGGC+TCCAATTACG | 98.88 | 7.257 | 4.1738 | 0.0580279 | 0.154692291 | 0.18218749 | 0.62099826 | 0.57551081 |
| Med-46 | Med | Tag | 14P0031 | AACACCCAAT+TGAAGACCCT | 99.55 | 8.2009 | 7.7451 | 0.0460969 | 0.177769456 | 0.10071067 | 0.19162754 | 0.15372526 |
| Med-47 | Med | Tag | 14P0330 | TAATCTCCTC+AGAAAGCGAT | 99.52 | 8.2132 | 5.44 | 0.0393417 | 0.17656702 | 0.10559893 | 0.0435043 | 0.12832462 |
| Med-5 | Med | Tag | 20P1715 | AGCATAATCT+ATATCACCGG | 99.54 | 8.5006 | 4.8459 | 0.0373973 | 0.175313768 | 0.08873668 | 0.04169436 | -0.0576487 |
| Med-6 | Med | Tag | 20P1253 | TCTCGCTGAG+ACTGGTCGAG | 99.45 | 7.9206 | 4.595 | 0.047378 | 0.166480253 | 0.11991125 | -0.0858958 | 0.06049997 |
| Med-7 | Med | Tag | 20P1264 | CTCAGTATTG+CGCACCAGTG | 99.49 | 41.5539 | 18.0376 | 0.00166574 | 0.241442239 | 0.04731475 | -0.0070541 | -0.0575706 |
| Med-8 | Med | Tag | 18P0875 | GTCCAGGGAG+GTATCGGCCA | 99.05 | 7.5788 | 4.5463 | 0.0536256 | 0.161214831 | 0.14157889 | -0.0832676 | 0.04712896 |
| Med-9 | Med | Tag | 18P0878 | AATGAGTCGA+TAATGGTGAC | 99.52 | 9.0101 | 5.1521 | 0.0343136 | 0.180580296 | 0.10958825 | 0.0825768 | -0.23361 |
