## Supplemental Table 2 for "Ecological and evolutionary insights into the diversification of Atlantic bluefin tuna"

| n | Chromosome | Start pos | End pos | ID | Gene | Protein Name | Putative Function | Organism | Notes |
|---|---|---|---|---|---|---|---|---|---|
| 13618 | Chromosome10 | 42204 | 46789 | ThM00000013618 | fshr | thyrotropin receptor | receptor for TSH, plays role in gonadal development | Zebrafish | |
| 13625 | Chromosome10 | 147865 | 273378 | ThM00000013625 | PAM | peptidyl-glycine alpha-amidating monooxygenase A isoform X1 | Catalytic activity, activation of peptides | Zebrafish | |
| 13835 | Chromosome10 | 5673810 | 5681625 | ThM00000013835 | ANXA1 | Annexin A1-like | n the innate immune response; also identified as Ca and phosp | Human | |
| 13836 | Chromosome10 | 5684032 | 5695778 | ThM00000013836 | tmc2a | transmembrane channel-like protein 2-A | Probable ion channel required for hair cells; sensory response | Zebrafish | |
| 13837 | Chromosome10 | 5702903 | 5705098 | ThM00000013837 | zfand5b | AN1-type zinc finger protein 5b | Metal binding | Zebrafish | |
| 13838 | Chromosome10 | 5711500 | 5714507 | ThM00000013838 | cdc25b | G2/mitotic-specific cyclin-B1 | Essential control of the cell cycle at the mitosis transition | Zebrafish | |
| 13839 | Chromosome10 | 5740160 | 5740678 | ThM00000013839 | pmchl | Pro-melanin concentrating hormone-like protein | ays a role in skin pigmentation, induces malanin concentratio | Zebrafish | |
| 13840 | Chromosome10 | 5743645 | 5744017 | ThM00000013840 | NA | NA | NA | | |
| 14926 | Chromosome11 | 720012 | 741057 | ThM00000014926 | celf1-xp | CUGBP Elav-like family member 1 isoform X3 | mRNA binding, splicing | Zebrafish | |
| 14927 | Chromosome11 | 720012 | 741057 | ThM00000014927 | celf1-xp | CUGBP Elav-like family member 1 isoform X3 | mRNA binding, splicing | Zebrafish | |
| 14928 | Chromosome11 | 788633 | 796086 | ThM00000014928 | NA | NA | NA | | |
| 14929 | Chromosome11 | 788633 | 797280 | ThM00000014929 | NA | NA | NA | | |
| 14930 | Chromosome11 | 796245 | 797280 | ThM00000014930 | NA | NA | NA | | |
| 14933 | Chromosome11 | 1016755 | 1066443 | ThM00000014933 | ADAMTS17 | disintegrin and metalloproteinase with thrombospondin motifs 17 isoform X | Zinc ion binding cofactor | Sperm whale | |
| 14937 | Chromosome11 | 1393854 | 1394984 | ThM00000014937 | NA | NA | NA | NA | |
| 14938 | Chromosome11 | 1414817 | 1443214 | ThM00000014938 | cers3a | ceramide synthase 2 | Catalytic activity; lipid metabolism | Zebrafish | |
| 14939 | Chromosome11 | 1462098 | 1468433 | ThM00000014939 | cers3a | ceramide synthase 3 | Catalytic activity; lipid metabolism | Zebrafish | |
| 14948 | Chromosome11 | 1941555 | 2097379 | ThM00000014948 | nek10 | Serine/threonine-protein kinase Nek10 isoform X1 | ATP binding, animal organ development | Zebrafish | |
| 14949 | Chromosome11 | 1941555 | 2098254 | ThM00000014949 | nek10 | Serine/threonine-protein kinase Nek10 isoform X1 | ATP binding, animal organ development | Zebrafish | |
| 15887 | Chromosome11 | 27508256 | 27616225 | ThM00000015887 | loc121909861 | Protein diaphanous homolog 3 isoform X2 | Actin binding, cytoskeletal organization | Atlantic herring | |
| 15888 | Chromosome11 | 27621105 | 27636839 | ThM00000015888 | tdrkh | tudor domain-containing protein 3 isoform X1 | RNA binding, piRNA processing, spermatogenesis | Zebrafish | |
| 16098 | Chromosome11 | 32936223 | 32936423 | ThM00000016098 | NA | NA | NA | NA | |
| 16099 | Chromosome11 | 33043409 | 33132472 | ThM00000016099 | LUZP1 | leucine zipper protein 1 | artery development, neural fold bending, ventircular septum | Zebrafish | heart |
| 16702 | Chromosome12 | 5735749 | 5780713 | ThM00000016702 | arhgap44b | rho GTPase-activating protein 44-like isoform X4 | Neutotransmitter receptor transport; synaptic activity, | Zebrafish | |
| 16703 | Chromosome12 | 5735749 | 5780713 | ThM00000016703 | arhgap44b | rho GTPase-activating protein 44-like isoform X5 | Neutotransmitter receptor transport; synaptic activity, | Zebrafish | |
| 16704 | Chromosome12 | 5787316 | 5792713 | ThM00000016704 | tex47 | testis-expressed protein 47 | NA | Zebrafish | |
| 16705 | Chromosome12 | 5797345 | 5807978 | ThM00000016705 | ndel1a | nuclear distribution protein nudE-like 1-A isoform X3 | Microtubule binding, intracellular functions | Atlantic herring | |
| 16706 | Chromosome12 | 5813541 | 5815773 | ThM00000016706 | NA | NA | NA | NA | |
| 18925 | Chromosome14 | 67093 | 97303 | ThM00000018925 | senp3b | sentrin-specific protease 3b isoform X1 | Peptidase activity, proteolysis | Zebrafish | |
| 18926 | Chromosome14 | 106723 | 118603 | ThM00000018926 | EIF4A1 | eukaryotic translation initiation factor 4A1A | RNA helicase, required for mRNA binding to ribosome | Human | |
| 18927 | Chromosome14 | 107783 | 118603 | ThM00000018927 | EIF4A2 | eukaryotic translation initiation factor 4A1A | RNA helicase, required for mRNA binding to ribosome | Human | |
| 18957 | Chromosome14 | 771448 | 790652 | ThM00000018957 | pelp1 | proline glutamic acid- and leucine-rich protein 1 | Coactivator of estrogen receptor-mediated transcription | Human | |
| 18958 | Chromosome14 | 790888 | 795133 | ThM00000018958 | MED11 | mediator of RNA polymerase II transcription subunit 11 | Involved in regulated trnacription of RNA poly-II dependent | Human | |
| 18959 | Chromosome14 | 797754 | 811630 | ThM00000018959 | ARR3 | arrestin red cell-like | Retina specific signal transduction / Photoreceptor required | Striped mullet | |
| 18960 | Chromosome14 | 797754 | 811630 | ThM00000018960 | ARR4 | arrestin red cell-like | Retina specific signal transduction / Photoreceptor required | Striped mullet | |
| 18961 | Chromosome14 | 814217 | 829659 | ThM00000018961 | cd99l2 | CD99 antigen-like protein 2 | May function as a homophilic adhesion molecule | Zebrafish | |
| 18962 | Chromosome14 | 814217 | 829659 | ThM00000018962 | cd99l3 | CD99 antigen-like protein 3 | May function as a homophilic adhesion molecule | Zebrafish | |
| 18963 | Chromosome14 | 839601 | 850313 | ThM00000018963 | MTMR1 | myotubularin-related protein 1a isoform X1 | Lipid phosphatase; catalytic activity | Human | |
| 20420 | Chromosome15 | 3670497 | 3763412 | ThM00000020420 | scube1 | signal peptide CUB and EGF-like domain-containing protein 1 isoform X4 | Calcium ion binding, animal organ development | Zebrafish | |
| 20421 | Chromosome15 | 3774582 | 3790230 | ThM00000020421 | scube1 | signal peptide CUB and EGF-like domain-containing protein 1 isoform X4 | Calcium ion binding, animal organ development | Zebrafish | |
| 20422 | Chromosome15 | 3795034 | 3801215 | ThM00000020422 | samm50b | sorting and assembly machinery component 50 homolog B isoform X1 | Maintenance of the mitochondrial cristae | Zebrafish | |
| 20423 | Chromosome15 | 3806140 | 3830183 | ThM00000020423 | aldh1l2 | mitochondrial 10-formyltetrahydrofolate dehydrogenase isoform X1 | Metabolic process | Zebrafish | |
| 20424 | Chromosome15 | 3836608 | 3836992 | ThM00000020424 | NA | NA | NA | NA | |
| 20425 | Chromosome15 | 3837028 | 3837535 | ThM00000020426 | NA | NA | NA | NA | |
| 20426 | Chromosome15 | 3837028 | 3838228 | ThM00000020425 | NA | NA | NA | NA | |
| 20427 | Chromosome15 | 3841792 | 3842241 | ThM00000020427 | NA | NA | NA | NA | |
| 20428 | Chromosome15 | 3842304 | 3846079 | ThM00000020428 | no confident match | acidic repeat-containing protein-like isoform X2 | NA | NA | |
| 20429 | Chromosome15 | 3850760 | 3853029 | ThM00000020429 | nopchap1 | Uncharacterized protein C12orf45 homolog | box C/D snoRNP assembly | Zebrafish | |
| 20430 | Chromosome15 | 3855494 | 3893458 | ThM00000020430 | SLC41a2b | solute carrier family 41 member 3 | Magnesium transport | Zebrafish | *SLC gene |
| 20431 | Chromosome15 | 3929833 | 3950998 | ThM00000020431 | chst11 | carbohydrate sulfotransferase 11 isoform X2 | Catalyzes transfer for sulfate | Zebrafish | |
| 20432 | Chromosome15 | 4014002 | 4016553 | ThM00000020432 | NA | NA | NA | NA | |
| 20433 | Chromosome15 | 4017550 | 4027861 | ThM00000020433 | TXNRD1 | thioredoxin reductase 1 cytoplasmic-like isoform X2 | Cell redox homeostasis | Atlantic salmon | |
| 20434 | Chromosome15 | 4017550 | 4027861 | ThM00000020434 | TXNRD1 | thioredoxin reductase 1 cytoplasmic-like isoform X2 | Cell redox homeostasis | Atlantic salmon | |
| 20435 | Chromosome15 | 4017550 | 4027861 | ThM00000020435 | TXNRD1 | thioredoxin reductase 1 cytoplasmic-like isoform X2 | Cell redox homeostasis | Atlantic salmon | |
| 20436 | Chromosome15 | 4031348 | 4035343 | ThM00000020436 | nfybb | nuclear transcription factor Y beta a | DNA binding transcription activity | Zebrafish | |
| 20437 | Chromosome15 | 4036010 | 4054560 | ThM00000020437 | HAL | histidine ammonia-lyase-like | Histidine catabolic process | Human | |
| 20438 | Chromosome15 | 4036010 | 4054560 | ThM00000020438 | HAL | histidine ammonia-lyase-like | Histidine catabolic process | Human | |
| 20439 | Chromosome15 | 4064410 | 4083850 | ThM00000020439 | podxl2 | podocalyxin isoform X2 | Membrane protein | Sperm whale | |
| 20440 | Chromosome15 | 4092690 | 4118249 | ThM00000020440 | MKLN1 | muskelin 1 | cytoskeleton organization, cell-matrix adhesion, regulation | Domestic ferret | |
| 21142 | Chromosome15 | 22421608 | 22457350 | ThM00000021142 | srgap2 | SLIT-ROBO Rho GTPase-activating protein 1-like isoform X2 | Neurogenesis; regulates synapse maturation and density | Zebrafish | |
| 21143 | Chromosome15 | 22460141 | 22463453 | ThM00000021143 | Kics2 | KICSTOR subunit 2 | Cellular response to amino acid, glucose starvation | Zebrafish | |
| 21144 | Chromosome15 | 22471228 | 22483499 | ThM00000021144 | TMEM19 | transmembrane protein 19 | cellular component of membrane | Human | |
| 21145 | Chromosome15 | 22489352 | 22496145 | ThM00000021145 | LOC108679122 | Rab-3A-interacting protein isoform X1 | exocytosis | Amphipod | |
| 23433 | Chromosome17 | 29701425 | 29706720 | ThM00000023433 | COQ7 | 5-demethoxyubiquinone hydroxylase mitochondrial isoform X1 | NA | Leopard gecko | |
| 23434 | Chromosome17 | 29701425 | 29706720 | ThM00000023434 | COQ7 | 5-demethoxyubiquinone hydroxylase mitochondrial isoform X2 | NA | Leopard gecko | |
| 23435 | Chromosome17 | 29717007 | 29723171 | ThM00000023435 | notum2 | carboxylesterase notum2 | mediates deacylation of target proteins | Zebrafish | |
| 23436 | Chromosome17 | 29724886 | 29730093 | ThM00000023436 | rps15a | 40S ribosomal protein S15a | Component of small ribosomal unit | Zebrafish | |
| 23437 | Chromosome17 | 29769725 | 29803747 | ThM00000023437 | xxylt1 | xylosyltransferase 1 isoform X2 | O-glycan processing | Zebrafish | |
| 24117 | Chromosome18 | 4179003 | 4253860 | ThM00000024117 | fbln2 | fibulin-2 | Calcium ion binding, animal organ development | Zebrafish | |
| 24118 | Chromosome18 | 4253884 | 4256759 | ThM00000024118 | NA | NA | NA | NA | |
| 24119 | Chromosome18 | 4292139 | 4316324 | ThM00000024119 | hdac11 | histone deacetylase 11 isoform X1 | histone deacetylase activity | Zebrafish | |
| 24143 | Chromosome18 | 5069730 | 5091891 | ThM00000024143 | NA | Nuclear casein kinase and cyclin-dependent kinase substrate 1a | DNA binding | Freshwater eel | |
| 24144 | Chromosome18 | 5101845 | 5114204 | ThM00000024144 | SLC45 | solute carrier family 45 member 3 | sucrose transmembrane transporter activity | Zebrafish | |
| 24145 | Chromosome18 | 5122183 | 5125417 | ThM00000024145 | klf15 | Krueppel-like factor 15 | regulation of transcription by RNA polymerase | Zebrafish | |
| 24146 | Chromosome18 | 5140495 | 5147817 | ThM00000024146 | elk4 | ETS domain-containing protein Elk-4 | regulation of transcription by RNA polymerase | Zebrafish | |
| 25370 | Chromosome18 | 27492854 | 27512071 | ThM00000025370 | alpl | alkaline phosphatase tissue-nonspecific isozyme | metal ion binding, bone mineralization | Zebrafish | |
| 25371 | Chromosome18 | 27545304 | 27547646 | ThM00000025371 | GUCY2C | guanylin-like | guanylate cyclase activity | Human | |
| 25372 | Chromosome18 | 27551902 | 27552704 | ThM00000025372 | GUCY2C | guanylin-like | guanylate cyclase activity | Human | |
| 25373 | Chromosome18 | 27570231 | 27572171 | ThM00000025373 | NA | NA | NA | NA | |
| 25374 | Chromosome18 | 27573155 | 27577147 | ThM00000025374 | nmur3 | neuromedin-U receptor 2 | circadian behavior, neuropeptide receptor activity | Zebrafish | |
| 25375 | Chromosome18 | 27585059 | 27601813 | ThM00000025375 | lactb | putative beta-lactamase-like 1 isoform X2 | Peptidase activity, proteolysis | Zebrafish | |
| 25376 | Chromosome18 | 27585059 | 27601813 | ThM00000025376 | lactb | putative beta-lactamase-like 1 isoform X3 | Peptidase activity, proteolysis | Zebrafish | |
| 25377 | Chromosome18 | 27632625 | 27640594 | ThM00000025377 | NA | NA | NA | NA | |
| 25488 | Chromosome18 | 30102805 | 30108227 | ThM00000025488 | c1qc | C1qc protein | complement activation, classical pathway | Zebrafish | |
| 25489 | Chromosome18 | 30130529 | 30136698 | ThM00000025489 | no confident match | NA | NA | NA | |
| 25490 | Chromosome18 | 30142500 | 30148065 | ThM00000025490 | LOC108182724 | Butyrophilin-like protein 10 | plasma membrane | Zebrafish | |
| 25491 | Chromosome18 | 30154922 | 30157886 | ThM00000025491 | LOC110438744 | Erythroid membrane-associated protein-like | membrane component | Zebrafish | |
| 27169 | Chromosome19 | 29185375 | 29194391 | ThM00000027169 | slc16a1 | monocarboxylate transporter 1-like | monocarboxylic acid transmembrane transporter activity | Zebrafish | SLC gene |
| 27170 | Chromosome19 | 29201326 | 29208447 | ThM00000027170 | avpr2ab | vasopressin V2 receptor-like | regulation of systemic arterial blood pressure by vasopressin | Zebrafish | |
| 27171 | Chromosome19 | 29218179 | 29222759 | ThM00000027171 | NA | socitrate dehydrogenase [NAD] subunit gamma mitochondrial-like isoform X | tricarboxylic acid cycle; mitochondrion | Eastern oyster | |
| 27172 | Chromosome19 | 29218179 | 29226288 | ThM00000027172 | NA | socitrate dehydrogenase [NAD] subunit gamma mitochondrial-like isoform X | tricarboxylic acid cycle; mitochondrion | Eastern oyster | |
| 27173 | Chromosome19 | 29229469 | 29246391 | ThM00000027173 | fam120c | constitutive coactivator of PPAR-gamma-like protein 2 | cellular component of nucleus | Zebrafish | |
| 27174 | Chromosome19 | 29262113 | 29268833 | ThM00000027174 | loc12298230 | GTPase IMAP family member 4-like | GTP binding | Zebrafish | |
| 27181 | Chromosome19 | 29331378 | 29333035 | ThM00000027181 | NA | CMRF35-like molecule 1 isoform X2 | Cellular component of plasma membrane | Atlantic salmon | |
| 27182 | Chromosome19 | 29350518 | 29354498 | ThM00000027182 | NA | NA | NA | NA | |
| 27183 | Chromosome19 | 29363619 | 29372638 | ThM00000027183 | NA | NA | NA | NA | |
| 27184 | Chromosome19 | 29454399 | 29460954 | ThM00000027184 | NA | NA | NA | NA | |
| 27185 | Chromosome19 | 29467908 | 29474218 | ThM00000027185 | NA | NA | NA | NA | |
| 27186 | Chromosome19 | 29481105 | 29494449 | ThM00000027186 | NA | NA | NA | NA | |
| 27187 | Chromosome19 | 29498193 | 29506867 | ThM00000027187 | NA | NA | NA | NA | |
| 27188 | Chromosome19 | 29512360 | 29517162 | ThM00000027188 | NA | NA | NA | NA | |
| 27189 | Chromosome19 | 29519090 | 29523233 | ThM00000027189 | NA | NA | NA | NA | |
| 27190 | Chromosome19 | 29528977 | 29536614 | ThM00000027190 | NA | NA | NA | NA | |
| 27191 | Chromosome19 | 29528977 | 29536614 | ThM00000027191 | NA | NA | NA | NA | |
| 27192 | Chromosome19 | 29543831 | 29574807 | ThM00000027192 | cables1 | CDK5 and ABL1 enzyme substrate 2 | Nervous system development, regulation of cell cycle | Zebrafish | |
| 27193 | Chromosome19 | 29579946 | 29594778 | ThM00000027193 | ccm2l | cerebral cavernous malformations 2 protein-like isoform X2 | NA | Zebrafish | |
| 27194 | Chromosome19 | 29579946 | 29594778 | ThM00000027194 | ccm2l | cerebral cavernous malformations 2 protein-like isoform X2 | NA | Zebrafish | |
| 27195 | Chromosome19 | 29611247 | 29623208 | ThM00000027195 | xkr7b | XK-related protein 7 | apoptopic processes | Zebrafish | |
| 27196 | Chromosome19 | 29688439 | 29689050 | ThM00000027196 | xkr7b | XK-related protein 7 | apoptopic processes | Zebrafish | |
| 27197 | Chromosome19 | 29740827 | 29741114 | ThM00000027197 | NA | NA | NA | NA | |
| 27198 | Chromosome19 | 29762031 | 29766597 | ThM00000027198 | sox1 | transcription factor SOX-1 | transcription regulator, cell differentiation and anatomical | Zebrafish | |
| 27199 | Chromosome19 | 29822341 | 29838953 | ThM00000027199 | tcea3 | transcription elongation factor A protein 2 isoform X1 | Zinc ion binding | Zebrafish | |
| 27200 | Chromosome19 | 29822341 | 29838953 | ThM00000027200 | tcea3 | transcription elongation factor A protein 2 isoform X2 | Zinc ion binding | Zebrafish | |
| 27201 | Chromosome19 | 29850826 | 29872292 | ThM00000027201 | rgs19 | regulator of G-protein signaling 19 isoform X3 | NA | Zebrafish | |
| 27202 | Chromosome19 | 29872338 | 29883048 | ThM00000027202 | rgs19 | regulator of G-protein signaling 19 isoform X3 | NA | Zebrafish | |
| 27203 | Chromosome19 | 29927295 | 29932275 | ThM00000027203 | loc121896063 | Ig-like V-type domain-containing protein FAM187A | Membrane protein | Atlantic herring | |
| 27204 | Chromosome19 | 29965012 | 30061849 | ThM00000027204 | oprl1 | nociceptin receptor | Neuropeptide binding, signaling pathways | Zebrafish | |
| 19 | Chromosome1 | 452344 | 473989 | ThM00000000019 | smad4a | Mothers against decapentaplegic homolog | cardiac muscle tissue development, skeletal muscle tissue | Zebrafish | heart |
| 20 | Chromosome1 | 477020 | 491268 | ThM00000000020 | LOC108080282 | cidic leucine-rich nuclear phosphoprotein 32 family member E-like isoform X | NA | Fruit fly | |
| 21 | Chromosome1 | 500387 | 516052 | ThM00000000021 | no confident match | NA | NA | NA | |
| 1548 | Chromosome1 | 26747178 | 26751861 | ThM00000001548 | ENY2 | Transcription and mRNA export factor ENY2 | protein transport, regulation of transcription by RNA pol II | Zebra finch | |
| 1549 | Chromosome1 | 26754513 | 26763903 | ThM00000001549 | NA | NA | NA | NA | |
| 1591 | Chromosome1 | 27748672 | 27768421 | ThM00000001591 | no confident match | NA | NA | NA | |
| 1592 | Chromosome1 | 27770343 | 27801073 | ThM00000001592 | KLHL | kelch-like protein 32 | Protein binding | Human | |
| 1593 | Chromosome1 | 27804645 | 27811969 | ThM00000001593 | FZD6 | frizzled-6-like | Receptor for Wnt proteins, inner ear and midbrain | Human | |
| 1594 | Chromosome1 | 27813884 | 27819403 | ThM00000001594 | Cthrc1 | collagen triple helix repeat-containing protein 1-like | Inhibits collagen expression | Rat | |
| 1595 | Chromosome1 | 27820597 | 27825297 | ThM00000001595 | loc121905676 | mitochondrial folate transporter/carrier-like | transmembrane transport; mitochondrial inner membrane | Atlantic herring | SLC gene |
| 1596 | Chromosome1 | 27827199 | 27839289 | ThM00000001596 | dcaf13 | DDB1- and CUL4-associated factor 13 | rRNA processing, protein ubiwuitination, oocyte growth | Zebrafish | |
| 27392 | Chromosome20 | 5482 | 11888 | ThM00000027392 | col12a1b | Collagen alpha-1(XII) chain isoform X2 | endodermal cell differentiation, regeneration | Zebrafish | |
| 27393 | Chromosome20 | 35464 | 41389 | ThM00000027393 | NA | NA | NA | NA | |
| 27394 | Chromosome20 | 47403 | 85655 | ThM00000027394 | lamb4 | laminin subunit beta-4 isoform X1 | NA | Atlantic salmon | |
| 27395 | Chromosome20 | 47403 | 85655 | ThM00000027395 | lamb4 | laminin subunit beta-4 isoform X1 | NA | Atlantic salmon | |
| 27396 | Chromosome20 | 110828 | 146938 | ThM00000027396 | lamb4 | laminin subunit beta-4 isoform X1 | NA | Atlantic salmon | |
| 27441 | Chromosome20 | 1185297 | 1193702 | ThM00000027441 | ftsj1 | utative tRNA (cytidine(32)/guanosine(34)-2'-O)-methyltransferase isoform X | tRNA methylation | Zebrafish | |
| 29142 | Chromosome21 | 4327346 | 4410424 | ThM00000029142 | plekhg5b | pleckstrin homology-like domain family B member 1 isoform X1 | angiogenesis, blood vessel endothelial cell migration, Rho | Zebrafish | |
| 29143 | Chromosome21 | 4327346 | 4410424 | ThM00000029143 | plekhg5b | pleckstrin homology-like domain family B member 1 isoform X1 | angiogenesis, blood vessel endothelial cell migration, Rho | Zebrafish | |
| 29144 | Chromosome21 | 4414668 | 4446580 | ThM00000029144 | a2ml | alpha-2-macroglobulin isoform X2 | liver development; serine-type endopeptidase inhibitor | Zebrafish | |
| 29145 | Chromosome21 | 4414668 | 4446580 | ThM00000029145 | a2ml | alpha-2-macroglobulin isoform X2 | liver development; serine-type endopeptidase inhibitor | Zebrafish | |
| 29146 | Chromosome21 | 4455214 | 4455799 | ThM00000029146 | ALP1 | protein ALP1-like | metal ion binding | Zebrafish | |
| 29147 | Chromosome21 | 4461236 | 4465642 | ThM00000029147 | NA | NA | NA | NA | |
| 29148 | Chromosome21 | 4465661 | 4468275 | ThM00000029148 | NA | NA | NA | NA | |
| 29986 | Chromosome21 | 23417193 | 23419379 | ThM00000029986 | hic1l | Hic1l protein | regulation of cytokine production, immune system processes | Zebrafish | |
| 30002 | Chromosome21 | 23923194 | 24016931 | ThM00000030002 | pitpnm2 | membrane-associated phosphatidylinositol transfer protein 3 isoform X3 | metal ion binding, phospholipid transporter activity | Zebrafish | |
| 30003 | Chromosome21 | 23923194 | 24016931 | ThM00000030003 | pitpnm2 | membrane-associated phosphatidylinositol transfer protein 3 isoform X3 | metal ion binding, phospholipid transporter activity | Zebrafish | |
| 30004 | Chromosome21 | 23923194 | 24016931 | ThM00000030004 | pitpnm2 | membrane-associated phosphatidylinositol transfer protein 3 isoform X3 | metal ion binding, phospholipid transporter activity | Zebrafish | |
| 30005 | Chromosome21 | 24022635 | 24042449 | ThM00000030005 | kiaa0753 | protein moonraker isoform X1 | cranial skeletal system development, centriole replication | Zebrafish | |
| 30006 | Chromosome21 | 24022635 | 24042449 | ThM00000030006 | kiaa0753 | protein moonraker isoform X1 | cranial skeletal system development, centriole replication | Zebrafish | |
| 30007 | Chromosome21 | 24043133 | 24045546 | ThM00000030007 | txndc17 | thioredoxin domain-containing protein 17 | disulfide reductase; peroxidase activity | Zebrafish | |
| 30008 | Chromosome21 | 24046621 | 24050641 | ThM00000030008 | NXNL2 | nucleoredoxin isoform X1 | sensory perception | Cheetah | |
| 30125 | Chromosome21 | 26878283 | 26967787 | ThM00000030125 | GBE1 | alpha-glucan-branching enzyme | carbohydrate binding, cation binding, glycogen metabolic | Human | |
| 30126 | Chromosome21 | 26982092 | 26985889 | ThM00000030126 | loc121898712 | high affinity cationic amino acid transporter 1-like isoform X1 | transmembrane activity | Atlantic herring | SLC like |
| 30127 | Chromosome21 | 26982092 | 26988370 | ThM00000030127 | loc121898712 | high affinity cationic amino acid transporter 1-like isoform X1 | transmembrane activity | Atlantic herring | SLC like |
| 30128 | Chromosome21 | 26982092 | 26988370 | ThM00000030128 | loc121898712 | high affinity cationic amino acid transporter 1-like isoform X1 | transmembrane activity | Atlantic herring | SLC like |
| 30129 | Chromosome21 | 26985471 | 26988370 | ThM00000030129 | loc121898712 | high affinity cationic amino acid transporter 1-like isoform X1 | transmembrane activity | Atlantic herring | SLC like |
| 30130 | Chromosome21 | 26985928 | 26988370 | ThM00000030130 | loc121898712 | high affinity cationic amino acid transporter 1-like isoform X1 | transmembrane activity | Atlantic herring | SLC like |
| 30131 | Chromosome21 | 26996524 | 27016535 | ThM00000030131 | mtus2a | microtubule-associated tumor suppressor candidate 2 isoform X5 | microtubule binding | Zebrafish | |

| | | | | | | | | | |
|---|---|---|---|---|---|---|---|---|---|
| 30132 | Chromosome21 | 26996524 | 27016535 | ThM00000030132 | mtus2a | microtubule-associated tumor suppressor candidate 2 isoform X6 | microtubule binding | Zebrafish | |
| 30133 | Chromosome21 | 26996524 | 27016535 | ThM00000030133 | mtus2a | microtubule-associated tumor suppressor candidate 2 isoform X6 | microtubule binding | Zebrafish | |
| 30134 | Chromosome21 | 27019217 | 27023111 | ThM00000030134 | stoml3a | stomatin (EPB72)-like 3a isoform X1 | plasma membrane | Zebrafish | |
| 30135 | Chromosome21 | 27019217 | 27023111 | ThM00000030136 | stoml3a | stomatin (EPB72)-like 3a isoform X1 | plasma membrane | Zebrafish | |
| 30136 | Chromosome21 | 27019217 | 27026585 | ThM00000030135 | stoml3a | stomatin (EPB72)-like 3a isoform X1 | plasma membrane | Zebrafish | |
| 30137 | Chromosome21 | 27028538 | 27054090 | ThM00000030137 | foxo1a | forkhead box protein O1-A | regulates homeostasis in response to oxidative stress; may | Zebrafish | cardiac |
| 30138 | Chromosome21 | 27086555 | 27108860 | ThM00000030138 | cog6 | conserved oligomeric Golgi complex subunit 6 | vesicular trafficking between ER and golgi | Zebrafish | |
| 30139 | Chromosome21 | 27112918 | 27157268 | ThM00000030139 | lhfpl6 | LHFPL tetraspan subfamily member 6 protein | Membrane protein | Zebrafish | |
| 30140 | Chromosome21 | 27163794 | 27173040 | ThM00000030140 | nek3 | serine/threonine-protein kinase Nek3 isoform X1 | ATP binding, protein kinase activity, phosphorylation | Atlantic herring | |
| 30141 | Chromosome21 | 27168099 | 27173040 | ThM00000030141 | nek3 | serine/threonine-protein kinase Nek3 isoform X2 | ATP binding, protein kinase activity, phosphorylation | Atlantic herring | |
| 30142 | Chromosome21 | 27184886 | 27185932 | ThM00000030142 | lpar6a | lysophosphatidic acid receptor 6a | Receptor signaling pathway, angiogenesis | Killifish | |
| 30143 | Chromosome21 | 27199175 | 27205957 | ThM00000030143 | rcbtb1 | RCC1 and BTB domain-containing protein 2 isoform X1 | retina vasculature development in camera-type eye; blood | Zebrafish | interesting |
| 30144 | Chromosome21 | 27215127 | 27239854 | ThM00000030144 | fndc3a | fibronectin type-III domain-containing protein 3A isoform X1 | medial fin development | Zebrafish | interesting |
| 30145 | Chromosome21 | 27242495 | 27260526 | ThM00000030145 | fndc3a | fibronectin type-III domain-containing protein 3A isoform X1 | medial fin development | Zebrafish | interesting |
| 30146 | Chromosome21 | 27266469 | 27269748 | ThM00000030146 | MLNR | motilin receptor | calcium mediated signaling using extracellular calcium | Zebrafish | calcium |
| 30147 | Chromosome21 | 27275186 | 27291914 | ThM00000030147 | zgc:153184 | capZ-interacting protein isoform X3 | NA | NA | |
| 30148 | Chromosome21 | 27294217 | 27298025 | ThM00000030148 | loc121898809 | TPA-induced transmembrane protein homolog | membrane | NA | |
| 30149 | Chromosome21 | 27302861 | 27305863 | ThM00000030149 | guca3e | guanylyl cyclase-activating protein 3 | calcium ion binding, ferrous iron binding, response to | Zebrafish | lcium and vision |
| 30150 | Chromosome21 | 27307935 | 27308465 | ThM00000030150 | loc121899026 | mid1-interacting protein 1-B-like | gamma-tubulin complex binding, epethelial tube formation, | Ciclid | |
| 30151 | Chromosome21 | 27308952 | 27311172 | ThM00000030151 | ndufc2 | NADH dehydrogenase [ubiquinone] 1 subunit C2 | mitochondrial electron transport, respiration | Rainbow trout | intersting |
| 30152 | Chromosome21 | 27311531 | 27314927 | ThM00000030152 | rab30 | ras-related protein Rab-30 isoform X1 | Rab protein signal transduction; GTPase activity | Channel catfish | |
| 30153 | Chromosome21 | 27313463 | 27314927 | ThM00000030153 | rab30 | ras-related protein Rab-30 isoform X1 | Rab protein signal transduction; GTPase activity | Channel catfish | |
| 30154 | Chromosome21 | 27318641 | 27327772 | ThM00000030154 | prcp | lysosomal Pro-X carboxypeptidase | negative regulation of systemic arterial blood pressure, | Zebrafish | |
| 30155 | Chromosome21 | 27331971 | 27333203 | ThM00000030155 | fam181b | FAM181B | NA | Zebrafish | |
| 30156 | Chromosome21 | 27449312 | 27569554 | ThM00000030156 | tenm4 | teneurin-4 isoform X1 | protein heterodimerization activity, neuron development, | Zebrafish | |
| 30157 | Chromosome21 | 27584480 | 27588583 | ThM00000030157 | nars2 | probable asparagine--tRNA ligase mitochondrial | ATP binding, nucleic acid binding | Zebrafish | |
| 30158 | Chromosome21 | 27589694 | 27624509 | ThM00000030158 | gab1 | GRB2-associated-binding protein 2 isoform X1 | signaling adaptor activity, apiboly, signal transduction | Zebrafish | |
| 30159 | Chromosome21 | 27631693 | 27671994 | ThM00000030159 | MAML2 | mastermind-like protein 2 | positive regulation of transcription of Notch receptor target | Chinook salmon | |
| 30160 | Chromosome21 | 27678841 | 27700557 | ThM00000030160 | yap1 | transcriptional coactivator YAP1 | embryonic development and organ development | Zebrafish | |
| 30161 | Chromosome21 | 27705417 | 27714688 | ThM00000030161 | cep126 | centrosomal protein of 126 kDa | spindle organization, cilium assembly | Zebrafish | |
| 30162 | Chromosome21 | 27715599 | 27723541 | ThM00000030162 | trpc6a | short transient receptor potential channel 6 isoform X2 | calcium channel activity | Atlantic herring | Calcium |
| 30163 | Chromosome21 | 27715599 | 27723541 | ThM00000030163 | trpc6a | short transient receptor potential channel 6 isoform X2 | calcium channel activity | Atlantic herring | Calcium |
| 30164 | Chromosome21 | 27726181 | 27731363 | ThM00000030164 | pgr | progesterone receptor | zinc ion binding, ovulation, LH signaling pathway | Zebrafish | |
| 30165 | Chromosome21 | 27733902 | 27793636 | ThM00000030165 | loc121898900 | rho GTPase-activating protein 42 isoform X2 | signal transduction, cell development | Zebrafish | |
| 30166 | Chromosome21 | 27816113 | 27894270 | ThM00000030166 | cntnap5l | conactin-5 isoform X4 | anatomical structure development | Zebrafish | |
| 30167 | Chromosome21 | 27816113 | 27894270 | ThM00000030167 | cntnap5l | conactin-5 isoform X3 | anatomical structure development | Zebrafish | |
| 30168 | Chromosome21 | 27816113 | 27894270 | ThM00000030168 | cntnap5l | conactin-5 isoform X1 | anatomical structure development | Zebrafish | |
| 30169 | Chromosome21 | 27843002 | 27878787 | ThM00000030169 | cntnap5l | conactin-5 isoform X1 | anatomical structure development | Zebrafish | |
| 30593 | Chromosome22 | 3306223 | 3421211 | ThM00000030593 | spata16 | spermatogenesis-associated protein 16-like isoform X1 | spermatogenesis | Atlantic salmon | reproduction |
| 30594 | Chromosome22 | 3416137 | 3421211 | ThM00000030594 | spata16 | spermatogenesis-associated protein 16-like isoform X1 | spermatogenesis | Atlantic salmon | reproduction |
| 30616 | Chromosome22 | 4973808 | 5017029 | ThM00000030616 | slc1 | Amino acid transporter | L-aspartate import across plasma membrane, metal ion | Zebrafish | SLC gene |
| 30617 | Chromosome22 | 5022137 | 5031102 | ThM00000030617 | podn | podocan | extracellular space | Zebrafish | |
| 30618 | Chromosome22 | 5045547 | 5056603 | ThM00000030618 | scp2b | sterol carrier protein 2b isoform X2 | cytosol | Zebrafish | |
| 30619 | Chromosome22 | 5045547 | 5056988 | ThM00000030619 | scp2b | sterol carrier protein 2b isoform X2 | cytosol | Zebrafish | |
| 30620 | Chromosome22 | 5062111 | 5066294 | ThM00000030620 | cpt-2 | Choline/carnitine acyltransferase domain-containing protein | fatty acid beta-oxidation | C. elegans | |
| 30621 | Chromosome22 | 5067111 | 5069880 | ThM00000030621 | MAGOHB | Protein mago nashi homolog 2 | regulation of mRNA processing | Human | |
| 30622 | Chromosome22 | 5070693 | 5072342 | ThM00000030622 | AATF | protein AATF | part of small subunit processome, generate RNA folding and | Human | |
| 30623 | Chromosome22 | 5074996 | 5082538 | ThM00000030623 | no confident match | NA | NA | NA | |
| 30624 | Chromosome22 | 5084110 | 5089672 | ThM00000030624 | CALCOCO1 | Calcium-binding and coiled-coil domain-containing protein 1 | positive regulation of gene expression, signal transduction | Human | |
| 30625 | Chromosome22 | 5092417 | 5097807 | ThM00000030625 | aldh9a1a | 4-trimethylaminobutyraldehyde dehydrogenase A | cartinin biosynthetic process, cellular aldehyde metabolic | Zebrafish | |
| 30710 | Chromosome22 | 6959671 | 7045882 | ThM00000030710 | lpp | lipoma-preferred partner | metal ion binding, convergent extension involved in | Zebrafish | |
| 31047 | Chromosome22 | 11947450 | 12024881 | ThM00000031046 | celf5a | CUGBP Elav-like family member 5a isoform X8 | RNA binding | Atlantic herring | |
| 31048 | Chromosome22 | 11947450 | 12024881 | ThM00000031048 | celf5a | CUGBP Elav-like family member 5a isoform X8 | RNA binding | Atlantic herring | |
| 31049 | Chromosome22 | 12043018 | 12045182 | ThM00000031049 | ALP1 | protein ALP1-like | metal ion binding | Zebrafish | |
| 31621 | Chromosome22 | 23740012 | 23745799 | ThM00000031621 | | | | | |
| 31622 | Chromosome22 | 23754464 | 23761833 | ThM00000031622 | NA | NA | NA | NA | |
| 31623 | Chromosome22 | 23761988 | 23814399 | ThM00000031623 | adamts10 | disintegrin and metalloproteinase with thrombospondin motifs 10 isoform X | proteolysis, retina layer formation | Zebrafish | |
| 31624 | Chromosome22 | 23761988 | 23814399 | ThM00000031624 | adamts10 | disintegrin and metalloproteinase with thrombospondin motifs 10 isoform X | proteolysis, retina layer formation | Zebrafish | |
| 31667 | Chromosome22 | 24933406 | 25031922 | ThM00000031667 | raver2 | ribonucleoprotein PTB-binding 2 | RNA binding | Zebrafish | |
| 31668 | Chromosome22 | 25036285 | 25124862 | ThM00000031668 | cachd1 | VWFA and cache domain-containing protein 1 | voltage-gated calcium channel activity | Zebrafish | calcium |
| 31697 | Chromosome22 | 26035681 | 26065055 | ThM00000031697 | AP4M1 | AP-4 complex subunit mu-1 | protein localization, protein targeting to lysosome, vesicle- | Human | |
| 31698 | Chromosome22 | 26068327 | 26071072 | ThM00000031698 | NA | Type III endosome membrane protein TEMP | may be involved in membrane trafficking between | Mouse | |
| 32151 | Chromosome23 | 6104757 | 6107348 | ThM00000032151 | slitrk4 | SLIT and NTRK-like protein 4 isoform X1 | regulation of synapse activity | Zebrafish | |
| 32152 | Chromosome23 | 6121457 | 6135180 | ThM00000032152 | NA | NA | NA | NA | |
| 33069 | Chromosome23 | 25208049 | 25279214 | ThM00000033069 | fgfr1a | fibroblast growth factor receptor-like 1a | Fin pattern formation, olfactory nerve formation, fin | Zebrafish | interesting |
| 33070 | Chromosome23 | 25292479 | 25299825 | ThM00000033070 | RNF212 | Probable E3 SUMO-protein ligase RNF212 isoform X1 | metal ion binding, protein sumoylation, meitotic | White tailed deer | |
| 33073 | Chromosome23 | 25354202 | 25376463 | ThM00000033073 | CTBP1 | C-terminal-binding protein 1 | chromatin binding | Human | |
| 33074 | Chromosome23 | 25387603 | 25399912 | ThM00000033074 | maea | macrophage erythroblast attacher | cell cycle, plasma membrane, metal ion binding | Zebrafish | |
| 33075 | Chromosome23 | 25404449 | 25405348 | ThM00000033075 | hnrnpa0l | heterogeneous nuclear ribonucleoprotein A0-like | mRNA splicing | Zebrafish | |
| 33076 | Chromosome23 | 25405470 | 25409210 | ThM00000033076 | NA | NA | NA | NA | |
| 33077 | Chromosome23 | 25415479 | 25430093 | ThM00000033077 | klhl29 | kelch-like protein 3 isoform X2 | NA | Zebrafish | |
| 33078 | Chromosome23 | 25415479 | 25430093 | ThM00000033078 | klhl29 | kelch-like protein 3 isoform X2 | NA | Zebrafish | |
| 33079 | Chromosome23 | 25440352 | 25441173 | ThM00000033079 | NA | NA | NA | NA | |
| 33080 | Chromosome23 | 25442543 | 25467372 | ThM00000033080 | fam13b | protein FAM13B isoform X1 | cell development, signal transduction | Zebrafish | |
| 33081 | Chromosome23 | 25442543 | 25467372 | ThM00000033082 | fam13b | protein FAM13B isoform X2 | cell development, signal transduction | Zebrafish | |
| 33082 | Chromosome23 | 25442543 | 25495778 | ThM00000033081 | fam13b | protein FAM13B isoform X2 | cell development, signal transduction | Zebrafish | |
| 33083 | Chromosome23 | 25478109 | 25495778 | ThM00000033083 | fam13b | protein FAM13B isoform X5 | cell development, signal transduction | Zebrafish | |
| 33084 | Chromosome23 | 25495795 | 25500244 | ThM00000033084 | NA | NA | NA | NA | |
| 33085 | Chromosome23 | 25523895 | 25530547 | ThM00000033085 | cxcl14 | C-X-C motif chemokine 14 | cellular responses, cemotaxis, cell differentiation | Zebrafish | |
| 33086 | Chromosome23 | 25719941 | 25721782 | ThM00000033086 | lingo1b | eat and immunoglobulin-like domain-containing nogo receptor-inter | neuron development, myelination | Zebrafish | |
| 33087 | Chromosome23 | 25720402 | 25721782 | ThM00000033087 | lingo1b | eat and immunoglobulin-like domain-containing nogo receptor-inter | neuron development, myelination | Zebrafish | |
| 33088 | Chromosome23 | 26215443 | 26219889 | ThM00000033088 | eif4ea | Eukaryotic translation initiation factor 4E-1A | regulation of translation | Zebrafish | |
| 33089 | Chromosome23 | 26215443 | 26219946 | ThM00000033089 | eif4ea | Eukaryotic translation initiation factor 4E-1A | regulation of translation | Zebrafish | |
| 33090 | Chromosome23 | 26221422 | 26225572 | ThM00000033090 | ADH5 | Alcohol dehydrogenase class-3 | catalyzes oxidation of long-chain prmary alcohols; functions | Human | |
| 33091 | Chromosome23 | 26226968 | 26229652 | ThM00000033091 | dkc1 | H/ACA ribonucleoprotein complex subunit DKC1 | Definitive hemopoiesis, mrNA processing and pseudouridine | Zebrafish | |
| 33092 | Chromosome23 | 26229721 | 26234882 | ThM00000033092 | ADH5 | Alcohol dehydrogenase class-3 | catalyzes oxidation of long-chain prmary alcohols; functions | Human | |
| 33322 | Chromosome24 | 12918 | 28319 | ThM00000033323 | olfm2a | noelin-like | neural crest cell development, signal transduction | Zebrafish | |
| 33323 | Chromosome24 | 36803 | 37729 | ThM00000033324 | c9h9orf116 | UPF0691 protein C9orf116 | cytoskeleton | NA | |
| 33324 | Chromosome24 | 45631 | 53025 | ThM00000033326 | fnbp1a | formin-binding protein 1a isoform X2 | plasma membrane, signal transduction | Atlantic herring | |
| 33325 | Chromosome24 | 45631 | 73649 | ThM00000033325 | fnbp1a | formin-binding protein 1a isoform X2 | plasma membrane, signal transduction | Atlantic herring | |
| 33326 | Chromosome24 | 45631 | 73649 | ThM00000033327 | fnbp1a | formin-binding protein 1a isoform X2 | plasma membrane, signal transduction | Atlantic herring | |
| 33327 | Chromosome24 | 143452 | 151157 | ThM00000033328 | med27 | mediator of RNA polymerase II transcription subunit 27 | camera type eye development, retinal cone cell | Zebrafish | eyes |
| 33328 | Chromosome24 | 156607 | 195468 | ThM00000033329 | ntng2a | netrin-G2 | animal organ morphogenesis, tissue development | Zebrafish | |
| 33329 | Chromosome24 | 198188 | 203260 | ThM00000033330 | no confident match | NA | NA | NA | |
| 33330 | Chromosome24 | 204058 | 217111 | ThM00000033331 | TTF1 | transcription termination factor 1-like | negative regulation of DNA replication | Human | |
| 33331 | Chromosome24 | 223211 | 223834 | ThM00000033332 | mylk4a | myosin light chain cardiac muscle-like | myosin light chain binding, cardiac muscle contraction | Human | cardiac |
| 33332 | Chromosome24 | 225244 | 251381 | ThM00000033333 | rgs12a | regulator of G-protein signaling 3-like isoform X4 | signal transduction | Zebrafish | |
| 33366 | Chromosome24 | 905978 | 915326 | ThM00000033367 | NA | NA | NA | NA | |
| 33367 | Chromosome24 | 915357 | 931209 | ThM00000033368 | loc122969386 | zinc finger MIZ domain-containing protein 2-like isoform X1 | Zinc ion binding | Atlantic salmon | |
| 33368 | Chromosome24 | 915357 | 931209 | ThM00000033369 | loc122969386 | zinc finger MIZ domain-containing protein 2-like isoform X1 | Zinc ion binding | Atlantic salmon | |
| 33369 | Chromosome24 | 941484 | 944743 | ThM00000033370 | ppia | peptidyl-prolyl cis-trans isomerase A | Protein folding, platelet aggregation, positive regulation of | Human | |
| 33370 | Chromosome24 | 950176 | 954166 | ThM00000033371 | NA | NA | NA | NA | |
| 33371 | Chromosome24 | 950176 | 954166 | ThM00000033372 | NA | NA | NA | NA | |
| 33372 | Chromosome24 | 954478 | 961190 | ThM00000033373 | slc18a1 | chromaffin granule amine transporte | transmembrane transporter activity | Zebrafish | SLC gene |
| 33373 | Chromosome24 | 966862 | 983272 | ThM00000033374 | slc18a1 | chromaffin granule amine transporte | transmembrane transporter activity | Zebrafish | SLC gene |
| 33376 | Chromosome24 | 1034784 | 1135379 | ThM00000033377 | cntrl | centriolin isoform X1 | chromosome condensation, chromatin binding | Zebrafish | |
| 33377 | Chromosome24 | 1137133 | 1188321 | ThM00000033378 | cntrl | centriolin isoform X1 | chromosome condensation, chromatin binding | Zebrafish | |
| 33526 | Chromosome24 | 3791721 | 3795658 | ThM00000033527 | loc122968105 | natterin-3-like isoform X2 | membrane | Atlantic herring | |
| 33527 | Chromosome24 | 3791721 | 3795667 | ThM00000033528 | loc122968105 | natterin-3-like isoform X2 | membrane | Atlantic herring | |
| 33528 | Chromosome24 | 3800243 | 3800763 | ThM00000033529 | harbi1 | putative nuclease HARBI1 | metal ion binding | Zebrafish | |
| 33529 | Chromosome24 | 3807575 | 3810573 | ThM00000033530 | loc122971239 | natterin-3-like | NA | Zebrafish | |
| 33530 | Chromosome24 | 3816436 | 3818155 | ThM00000033531 | loc122971239 | natterin-3-like | NA | Zebrafish | |
| 33531 | Chromosome24 | 3821250 | 3827984 | ThM00000033532 | ptrh1 | probable peptidyl-tRNA hydrolase isoform X2 | RNA binding | Human | |
| 33532 | Chromosome24 | 3825678 | 3827984 | ThM00000033533 | ptrh1 | probable peptidyl-tRNA hydrolase isoform X2 | RNA binding | Human | |
| 33533 | Chromosome24 | 3846583 | 3866646 | ThM00000033534 | tor2a | prosalusin | endocrine and/or paracrine factors able to increase | Human | |
| 33534 | Chromosome24 | 3868169 | 3876680 | ThM00000033535 | sh2d3cb | SH2 domain containing 3Cb isoform X2 | GTPase mediated signal transduction | Human | |
| 33536 | Chromosome24 | 3879936 | 3881516 | ThM00000033537 | ugt2b1 | UDP-glucuronosyltransferase 2A1-like | glucuronosyltransferase activity | Zebrafish | |
| 33537 | Chromosome24 | 3886274 | 3887854 | ThM00000033538 | ugt2b1 | UDP-glucuronosyltransferase 2A1-like | glucuronosyltransferase activity | Zebrafish | |
| 33538 | Chromosome24 | 3889523 | 3912053 | ThM00000033539 | sh2d3cb | SH2 domain containing 3Cb isoform X2 | GTPase mediated signal transduction | Human | |
| 33539 | Chromosome24 | 3964444 | 3964806 | ThM00000033541 | ZNF682 | zinc finger protein 862-like | metal ion binding, transcription regulation | Human | |
| 33540 | Chromosome24 | 3964444 | 3965089 | ThM00000033540 | ZNF682 | zinc finger protein 862-like | metal ion binding, transcription regulation | Human | |
| 33541 | Chromosome24 | 3965211 | 3966401 | ThM00000033542 | NA | NA | NA | NA | |
| 33542 | Chromosome24 | 3986890 | 4036657 | ThM00000033543 | LOC104949344 | Spermatid perinuclear RNA-binding protein-like isoform X1 | siRNA binding, regulation of regulatory ncRNA processing, | Black rockcod | |
| 2651 | Chromosome2 | 22642840 | 22643874 | ThM00000002651 | sox1a | Transcription factor Sox-1a | brain development, negative regulation of transcription, | Zebrafish | |
| 2670 | Chromosome2 | 23194866 | 23197628 | ThM00000002670 | filip1l | Filamin A-interacting protein 1-like | cytoplasm | Zebrafish | |
| 2671 | Chromosome2 | 23210036 | 23211827 | ThM00000002671 | filip1l | Filamin A-interacting protein 1-like | cytoplasm | Zebrafish | |
| 2672 | Chromosome2 | 23222334 | 23226265 | ThM00000002672 | csms1 | protein CMSS1 isoform X1 | nucleus, 90S preribosome | Zebrafish | |
| 2673 | Chromosome2 | 23227046 | 23229253 | ThM00000002673 | tmem30c | transmembrane protein 30C | plasma membrane, phospholipid translocation | Mouse | |
| 2674 | Chromosome2 | 23232243 | 23250200 | ThM00000002674 | dcbld2 | discoidin CUB AND LCCL domain-containing protein 2 isoform X2 | signaling receptor activity, wound healing, regulation of | Polar bear /Atlantic herring | |
| 2881 | Chromosome2 | 27103463 | 27255817 | ThM00000002880 | karlna | kalirin isoform X1 | cytoplasm, kinase activity, intracellular signal transduction | Atlantic herring | |
| 2883 | Chromosome2 | 27149942 | 27255817 | ThM00000002883 | karlna | kalirin isoform X1 | cytoplasm, kinase activity, intracellular signal transduction | Atlantic herring | |
| 3059 | Chromosome2 | 31881315 | 31882296 | ThM00000003059 | NA | NA | NA | NA | |
| 3060 | Chromosome2 | 31891043 | 31910181 | ThM00000003060 | FRZB | secreted frizzled-related protein 3 | modulator of Wnt signaling through direct interaction with | Polar bear/ Zebrafish | |
| 3061 | Chromosome2 | 31920299 | 31924709 | ThM00000003061 | dusp19b | dual specificity protein phosphatase 19-like | cytoplasm, kinase activity, dephosphorylation | Zebrafish | |
| 3062 | Chromosome2 | 31941634 | 31971368 | ThM00000003062 | pde1a | calcium/calmodulin-dependent-cyclic nucleotide phosphodiesterase 1A; | calmodulin binding, metal ion binding, cAMP-mediated | Mouse | |
| 3169 | Chromosome3 | 1345499 | 1458315 | ThM00000003169 | KIAA1217 | sickle tail protein homolog isoform X3 | NA | Atlantic salmon | |
| 3170 | Chromosome3 | 1468642 | 1477691 | ThM00000003170 | Paxip1 | PAX-interacting protein 1 | Adipose tissue development, chroion development, | Mouse | |
| 3198 | Chromosome3 | 1913003 | 1913957 | ThM00000003198 | no confident match | Junctophilin-1-like isoform X2 | NA | NA | |
| 3199 | Chromosome3 | 1933053 | 1958747 | ThM00000003199 | no confident match | Junctophilin-1-like isoform X2 | NA | NA | |
| 4170 | Chromosome3 | 18953700 | 18956594 | ThM00000004170 | MYH13 | Myosin-13 | Mediates cobntractions of striated muscles; muscle | Human | |
| 4171 | Chromosome3 | 18964441 | 18999638 | ThM00000004171 | crhr1 | Corticotropin-releasing factor receptor 1 | receptor signaling pathway, cell surface receptor pathway, | Zebrafish | |
| 4172 | Chromosome3 | 19009437 | 19016837 | ThM00000004172 | cd59b | CD59 glycoprotein isoform X2 | potassium ion import across plasma membrane, regulation | Zebrafish | |
| 5062 | Chromosome4 | 4503083 | 4582948 | ThM00000005062 | galnt10 | polypeptide N-acetylgalactosaminyltransferase 10-like | golgi membrane, carbohydrate binding, glycosylation | Zebrafish | |
| 5063 | Chromosome4 | 4877359 | 4940730 | ThM00000005063 | galnt10 | polypeptide N-acetylgalactosaminyltransferase 10-like | golgi membrane, carbohydrate binding, glycosylation | Zebrafish | |
| 5064 | Chromosome4 | 5104065 | 5105485 | ThM00000005064 | NA | NA | NA | NA | |
| 5065 | Chromosome4 | 5111041 | 5118464 | ThM00000005065 | sap30l | histone deacetylase complex subunit SAP30L | DNA, histone, and nucleosome binding; heart looping, | Zebrafish | |
| 5066 | Chromosome4 | 5125671 | 5127538 | ThM00000005066 | oc121910685-xp | heart- and neural crest derivatives-expressed protein 1-like | protein dimerization, axon, forebrain, and sensory organ | Octopus | |
| 5067 | Chromosome4 | 5190546 | 5193002 | ThM00000005068 | loc121910757 | cornifelin homolog B-like | membrane cellular component | Zebrafish | |
| 5068 | Chromosome4 | 5190546 | 5195800 | ThM00000005067 | loc121910757 | cornifelin homolog B-like | membrane cellular component | Zebrafish | |
| 5069 | Chromosome4 | 5200338 | 5201522 | ThM00000005069 | NA | NA | NA | NA | |
| 5070 | Chromosome4 | 5204661 | 5208378 | ThM00000005070 | NA | NA | NA | NA | |
| 5071 | Chromosome4 | 5211299 | 5215783 | ThM00000005071 | hnrnpl | heterogeneous nuclear ribonucleoprotein A/B-like isoform X1 | mRNA binding, regulation of RNA splicing | Zebrafish | |
| 5072 | Chromosome4 | 5211299 | 5215783 | ThM00000005072 | hnrnpl | heterogeneous nuclear ribonucleoprotein A/B-like isoform X1 | mRNA binding, regulation of RNA splicing | Zebrafish | |
| 5073 | Chromosome4 | 5211299 | 5215783 | ThM00000005073 | hnrnpl | heterogeneous nuclear ribonucleoprotein A/B-like isoform X1 | mRNA binding, regulation of RNA splicing | Zebrafish | |
| 5074 | Chromosome4 | 5223900 | 5227264 | ThM00000005074 | btg4 | protein BTG4 | negative regulation of cell cycle | Zebrafish | |
| 5075 | Chromosome4 | 5239975 | 5255504 | ThM00000005075 | SLC25A14 | brain mitochondrial carrier protein 1 isoform X1 | component of mitochondrial envelope, chloride | Polar bear | **SLC gene |
| 5076 | Chromosome4 | 5259878 | 5264328 | ThM00000005076 | tnnt3 | troponin T skeletal muscle | troponin C, I binding, regulation of striated muscle | Zebrafish | |

| | | | | | | | | | |
|---|---|---|---|---|---|---|---|---|---|
| 5077 | Chromosome4 | 5266135 | 5268676 | TthM00000005077 | Gpr119 | glucose-dependent insulinotropic receptor isoform X1 | G protein-coupled receptor activity, phosphatidylcholine | Mouse | |
| 5078 | Chromosome4 | 5273014 | 5289107 | TthM00000005078 | Clk4 | dual specificity protein kinase CLK4-like | protein kinase activity, phosphorylation, regulation of RNA | Mouse | |
| 5079 | Chromosome4 | 5273014 | 5289107 | TthM00000005079 | Clk4 | dual specificity protein kinase CLK4-like | protein kinase activity, phosphorylation, regulation of RNA | Mouse | |
| 5080 | Chromosome4 | 5296575 | 5306000 | TthM00000005080 | Arhgap22 | rho GTPase-activating protein 22-like | GTPase activator, angiogenesis, cell differentiation, signal | Mouse | |
| 5081 | Chromosome4 | 5313704 | 5319610 | TthM00000005081 | p33monox | putative monooxygenase p33MONOX | oxidoreductase activity | Zebrafish | |
| 5082 | Chromosome4 | 5313704 | 5319610 | TthM00000005082 | p33monox | putative monooxygenase p33MONOX | oxidoreductase activity | Zebrafish | |
| 5083 | Chromosome4 | 5319636 | 5322401 | TthM00000005083 | NA | NA | NA | NA | |
| 5084 | Chromosome4 | 5324164 | 5329481 | TthM00000005084 | pof1b | protein POF1B isoform X2 | actin filament binding and organization, bicellular tight | Lake trout | |
| 5085 | Chromosome4 | 5343274 | 5360917 | TthM00000005085 | pof1b | protein POF1B isoform X2 | actin filament binding and organization, bicellular tight | Lake trout | |
| 5086 | Chromosome4 | 5400192 | 5401100 | TthM00000005086 | SPRY3 | protein sprouty homolog 3 | Inhibits neurite branching, arbor length and neurite | Mouse | |
| 5307 | Chromosome4 | 11467849 | 11610647 | TthM00000005307 | fstl4 | Follistatin-like 4 | calcium ion binding, cell differentiation, regulation of BMP | Zebrafish | |
| 5308 | Chromosome4 | 11467849 | 11610647 | TthM00000005308 | fstl4 | Follistatin-like 4 | calcium ion binding, cell differentiation, regulation of BMP | Zebrafish | |
| 5702 | Chromosome4 | 19353454 | 19382618 | TthM00000005702 | nrxn2a | neurexin 2a isoform X5; | axon guidance, chemical synaptic transmission, motor | Zebrafish | |
| 5703 | Chromosome4 | 19353454 | 19449682 | TthM00000005703 | nrxn2a | neurexin 2a isoform X5; | axon guidance, chemical synaptic transmission, motor | Zebrafish | |
| 5704 | Chromosome4 | 19410386 | 19449682 | TthM00000005704 | nrxn2a | neurexin 2a isoform X5; | axon guidance, chemical synaptic transmission, motor | Zebrafish | |
| 5705 | Chromosome4 | 19410386 | 19449682 | TthM00000005705 | nrxn2a | neurexin 2a isoform X5; | axon guidance, chemical synaptic transmission, motor | Zebrafish | |
| 6517 | Chromosome5 | 5399248 | 5403081 | TthM00000006517 | oc122997364 | FAM161A-like | cytoskeleton protein | Atlantic herring | |
| 6518 | Chromosome5 | 5404782 | 5415230 | TthM00000006518 | NA | NA | NA | NA | |
| 6519 | Chromosome5 | 5451591 | 5523044 | TthM00000006520 | cntn1a | contactin-associated protein-like 4 | Mediates cell surface interactions during nervous system | Zebrafish | |
| 6520 | Chromosome5 | 5451591 | 5523196 | TthM00000006519 | cntn1a | contactin-associated protein-like 4 | Mediates cell surface interactions during nervous system | Zebrafish | |
| 6521 | Chromosome5 | 5581713 | 5584380 | TthM00000006521 | CIAO2B | Cytosolic iron-sulfur assembly component 2B | Chromosome segregation, iron-sulfur cluster assembly, | Human | |
| 6522 | Chromosome5 | 5581713 | 5584380 | TthM00000006522 | CIAO2B | Cytosolic iron-sulfur assembly component 2B | Chromosome segregation, iron-sulfur cluster assembly, | Human | |
| 6523 | Chromosome5 | 5593702 | 5607803 | TthM00000006523 | zbed4 | Zinc finger BED domain-containing protein 4 | DNA binding, metal ion binding, protein dimerization activity | Zebrafish | |
| 7716 | Chromosome6 | 2069132 | 2073998 | TthM00000007716 | NA | NA | NA | NA | |
| 7717 | Chromosome6 | 2159755 | 2165705 | TthM00000007717 | Rnf133 | E3 ubiquitin-protein ligase RING2-A-like isoform X1 | Metal ion binding, protein ligase activity, protein catabolic | Mouse | |
| 7718 | Chromosome6 | 2159755 | 2165705 | TthM00000007718 | Rnf133 | E3 ubiquitin-protein ligase RING2-A-like isoform X1 | Metal ion binding, protein ligase activity, protein catabolic | Mouse | |
| 7719 | Chromosome6 | 2169299 | 2173789 | TthM00000007719 | RPS18 | 40S ribosomal protein S18 | Component of small ribosomal unit | Human | |
| 7720 | Chromosome6 | 2174102 | 2191700 | TthM00000007720 | NA | NA | NA | NA | |
| 7721 | Chromosome6 | 2174102 | 2191700 | TthM00000007721 | NA | NA | NA | NA | |
| 9012 | Chromosome7 | 24575 | 32929 | TthM00000009012 | cinp | Cyclin-dependent kinase 2-interacting protein isoform X1 | kinase activity | Atlantic salmon | |
| 9013 | Chromosome7 | 34747 | 46662 | TthM00000009013 | pclob | Protein piccolo isoform X2 | metal ion binding, presynaptic active zone assembly, protein | Zebrafish | |
| 9014 | Chromosome7 | 55773 | 59348 | TthM00000009014 | tecpr1a | Tectonin beta-propeller repeat-containing protein 1 isoform X2 | intracellular membrane-bound organelle | Zebrafish | |
| 9015 | Chromosome7 | 55773 | 88182 | TthM00000009015 | tecpr1a | Tectonin beta-propeller repeat-containing protein 1 isoform X2 | intracellular membrane-bound organelle | Zebrafish | |
| 9016 | Chromosome7 | 55773 | 88182 | TthM00000009016 | tecpr1a | Tectonin beta-propeller repeat-containing protein 1 isoform X2 | intracellular membrane-bound organelle | Zebrafish | |
| 9017 | Chromosome7 | 89389 | 90219 | TthM00000009017 | ANK1 | Ankyrin-1 | exocytosis, protein localization to plasma membrane, signal | Human | |
| 9018 | Chromosome7 | 90381 | 91315 | TthM00000009018 | NA | NA | NA | NA | |
| 11075 | Chromosome8 | 13707758 | 13722207 | TthM00000011075 | AKAP7 | A-kinase anchor protein 7 isoform gamma | protein kinase binding, modulation of chemical synaptic | Human | |
| 11076 | Chromosome8 | 13736166 | 13747706 | TthM00000011076 | AKAP7 | A-kinase anchor protein 7 isoform gamma | protein kinase binding, modulation of chemical synaptic | Human | |
| 11077 | Chromosome8 | 13749250 | 13749819 | TthM00000011078 | NA | NA | NA | NA | |
| 11078 | Chromosome8 | 13749250 | 13752246 | TthM00000011077 | NA | NA | NA | NA | |
| 11079 | Chromosome8 | 13751134 | 13752246 | TthM00000011079 | NA | NA | NA | NA | |
| 11080 | Chromosome8 | 13763776 | 13766123 | TthM00000011080 | NA | NA | NA | NA | |
| 11081 | Chromosome8 | 13766388 | 13766653 | TthM00000011081 | LOC110439791 | G2/M phase-specific E3 ubiquitin-protein ligase-like | ubiquitin-protein transferase activity | Zebrafish | |
| 11082 | Chromosome8 | 13766388 | 13767187 | TthM00000011082 | LOC110439791 | G2/M phase-specific E3 ubiquitin-protein ligase-like | ubiquitin-protein transferase activity | Zebrafish | |
| 11656 | Chromosome8 | 24495614 | 24510978 | TthM00000011656 | loc121882320 | dystrobrevin beta-like isoform X3 | Zinc ion binding | Atlantic salmon | nds to dystrophin |
| 11657 | Chromosome8 | 24529655 | 24583410 | TthM00000011657 | dnmt3aa | DNA (cytosine-5)-methyltransferase 3A-like isoform X1 | DNA methylation, cellular response to hypoxia; epigenetic | Zebrafish | |
| 11658 | Chromosome8 | 24529655 | 24583410 | TthM00000011658 | dnmt3aa | DNA (cytosine-5)-methyltransferase 3A-like isoform X1 | DNA methylation, cellular response to hypoxia; epigenetic | Zebrafish | |
| 11659 | Chromosome8 | 24529655 | 24583410 | TthM00000011659 | dnmt3aa | DNA (cytosine-5)-methyltransferase 3A-like isoform X1 | DNA methylation, cellular response to hypoxia; epigenetic | Zebrafish | |
| 11660 | Chromosome8 | 24529655 | 24583410 | TthM00000011660 | dnmt3aa | DNA (cytosine-5)-methyltransferase 3A-like isoform X1 | DNA methylation, cellular response to hypoxia; epigenetic | Zebrafish | |
| 11661 | Chromosome8 | 24587456 | 24620987 | TthM00000011661 | loc115574212 | dihydropyrimidinase-related protein 5-like | Hydrolase activity on carbon-nitrogen bonds | Brown trout | |
| 11667 | Chromosome8 | 24701792 | 24797303 | TthM00000011667 | scara5 | scavenger receptor class A member 5 | Ferritin receptor that mediates deliver of iron to the cell; | Zebrafish | |
| 11668 | Chromosome8 | 24805450 | 24806310 | TthM00000011668 | samd12 | sterile alpha motif domain-containing protein 12-like | Chromatin organization | Zebrafish | |
| 11669 | Chromosome8 | 24805450 | 24807406 | TthM00000011669 | samd12 | sterile alpha motif domain-containing protein 12-like | Chromatin organization | Zebrafish | |
| 11670 | Chromosome8 | 24832941 | 24834966 | TthM00000011670 | ext1a | exostosin-1 | Metal ion binding, protein glycosylation | Zebrafish | |
