## Supplemental Table 3 for "Ecological and evolutionary insights into the diversification of Atlantic bluefin tuna"

| n | Chromosome | Start pos | End pos | ID | Gene | Protein Name | utative functi | Organism | Notes | Variable |
|---|---|---|---|---|---|---|---|---|---|---|
| 4679 | Chromosome 3 | 30953264 | 30962736 | TthM00000004679 | vcpip1 | uitinating protein | plasic reticulu | Zebrafish | | MLD |
| 4680 | Chromosome 3 | 30969537 | 30971735 | TthM00000004680 | NA | NA | NA | NA | | MLD |
| 6716 | Chromosome 5 | 12112775 | 12120732 | TthM00000006716 | grem2a | Gremlin-2 | IP from recept | Zebrafish | | MLD |
| 6717 | Chromosome 5 | 12121071 | 12169870 | TthM00000006717 | rgs7bpb | G-protein signal | pathway; neg | Zebrafish | | MLD |
| 7388 | Chromosome 5 | 25190734 | 25231084 | TthM00000007388 | gabrp | butyric acid rece | ansmembrane | Zebrafish | Cl ion | MLD |
| 12303 | Chromosome 9 | 8733310 | 8743309 | TthM00000012303 | xpp3 | minopeptidase 3 | proteolysis | Atlantic salmon | | MLD |
| 12304 | Chromosome 9 | 8749704 | 8781828 | TthM00000012304 | confident ma | NA | NA | NA | | MLD |
| 12676 | Chromosome 9 | 14703383 | 14710591 | TthM00000012676 | SLC5A3 | ositol cotranspor | port, inositol | Sea lamprey | | MLD |
| 12677 | Chromosome 9 | 14712390 | 14725555 | TthM00000012677 | arhgap17b | ctivating protein | al transductio | Zebrafish | | MLD |
| 12678 | Chromosome 9 | 14712390 | 14736955 | TthM00000012678 | | | | | | MLD |
| 12679 | Chromosome 9 | 14740695 | 14742445 | TthM00000012679 | aqp8b | Aquaporin-8b | vater transpo | Zebrafish | egulation, rep | MLD |
| 12680 | Chromosome 9 | 14746191 | 14750498 | TthM00000012680 | metrnl | eteorin-like prote | onse to cold, | Zebrafish | oregulation, e | MLD |
| 14541 | Chromosome 10 | 22195372 | 22200971 | TthM00000014541 | nlrc3l | main-containing | evelopment, | Zebrafish | | MLD |
| 14542 | Chromosome 10 | 22206883 | 22209668 | TthM00000014542 | ankra2 | repeat family A p | on of gene exp | Atlantic salmon | | MLD |
| 14543 | Chromosome 10 | 22212117 | 22215209 | TthM00000014543 | CH211-157C1 | scription factor | nal transduct | Killifish | odium chann | MLD |
| 14817 | Chromosome 10 | 28134970 | 28175497 | TthM00000014817 | cse1l | Exportin-2 | m nucleus, bo | Zebrafish | | MLD |
| 14818 | Chromosome 10 | 28180915 | 28186521 | TthM00000014818 | pbdc1 | Protein PBDC1 | cytoplasm | Black rockcod | | MLD |
| 14820 | Chromosome 10 | 28198698 | 28206933 | TthM00000014820 | dmtn | Dematin | ization, lame | Zebrafish | | MLD |
| 22283 | Chromosome 17 | 3781050 | 3816138 | TthM00000022283 | fat1a | otocadherin Fat | sure, proneph | Zebrafish | um, vision, ki | MLD |
| 28236 | Chromosome 20 | 17196514 | 17203808 | TthM00000028236 | SLC28A1 | ucleoside cotran | ter, transmen | Rat | *SLC | MLD |
| 28237 | Chromosome 20 | 17205943 | 17218227 | TthM00000028237 | alpk3a | otein kinase 3 is | NA | Zebrafish | | MLD |
| 28238 | Chromosome 20 | 17220682 | 17227905 | TthM00000028238 | malt3 | oid tissue lymph | proteolysis | Zebrafish | | MLD |
| 9892 | Chromosome 7 | 17271368 | 17288344 | TthM00000009892 | kif26ab | e protein KIF26A | ubule-based n | Zebrafish | | SSH |
| 20751 | Chromosome 15 | 11821573 | 11842105 | TthM00000020751 | nrcama | adhesion molecu | dhesion, brai | Zebrafish | Swimming | SSH |
| 20753 | Chromosome 15 | 11849692 | 11859206 | TthM00000020753 | cntn1b | ntactin 1b isofor | dance, cell-ce | Channel catfish | | SSH |
| 20874 | Chromosome 15 | 14952492 | 15023700 | TthM00000020874 | chchd3a | coil-helix domai | dria membra | Zebrafish | | SSH |
| 26847 | Chromosome 19 | 23705553 | 23709839 | TthM00000026848 | dlgap4b | ssociated protei | al synaptic tr | Zebrafish | | SSH |
| 29646 | Chromosome 21 | 17376819 | 17403304 | TthM00000029646 | Gosr2 | receptor comple | tein transport | Rat | | SSH |
| 29648 | Chromosome 21 | 17407653 | 17425168 | TthM00000029649 | cpda | arboxypeptidase | ic process, pr | Zebrafish | | SSH |
| 31027 | Chromosome 22 | 11671842 | 11676530 | TthM00000031027 | OC10652664 | ein ligase RNF12 | ein ubiquitina | Killifish | | SSH |
| 31028 | Chromosome 22 | 11679688 | 11690208 | TthM00000031029 | bdg | Basigin isoform X | ll-cell adhesi | Atlantic herring | | SSH |
| 31031 | Chromosome 22 | 11694623 | 11733897 | TthM00000031031 | hcn2b | tion-activated c | ne depolariza | Zebrafish | annel, swimm | SSH |
| 31301 | Chromosome 22 | 16545398 | 16567221 | TthM00000031301 | lrb | nsulin receptor | positive regul | Carp | | SSH |
| 33086 | Chromosome 23 | 25719941 | 25721782 | TthM00000033086 | lingo1b | ke domain-conta | on, neuron dev | Zebrafish | | SSH |
| 10287 | Chromosome 7 | 25336239 | 25341936 | TthM00000010287 | fosl1a | ted antigen 1a is | transcription | Zebrafish | | SST |
| 10289 | Chromosome 7 | 25352126 | 25353415 | TthM00000010289 | kcnk13b | annel, subfamily | nsport, regula | Zebrafish | * ion transpor | SST |
| 20574 | Chromosome 15 | 7388851 | 7393144 | TthM00000020574 | optn | Optineurin | acterium, pro | Zebrafish | * | EntryDay |
| 20575 | Chromosome 15 | 7395799 | 7403991 | TthM00000020575 | mcm10 | ein MCM10 hom | onse, DNA rep | Zebrafish | | EntryDay |
| 20576 | Chromosome 15 | 7406890 | 7413159 | TthM00000020576 | UCMAB | plate and cartila | NA | Killifish | | EntryDay |
| 20577 | Chromosome 15 | 7419057 | 7424934 | TthM00000020577 | phyh | noyl-CoA dioxyg | cid alpha-oxi | Zebrafish | | EntryDay |
| 22821 | Chromosome 17 | 15767063 | 15800994 | TthM00000022821 | lef1 | ncer-binding fac | NA | Zebrafish | | EntryDay |
